# Linking Oestradiol Timing and Tempo, Brain Development, and Mental Health in Adolescent Females

**DOI:** 10.1101/2025.04.23.650187

**Authors:** Muskan Khetan, Nandita Vijayakumar, Ye Ella Tian, Sarah Whittle

## Abstract

**Background:** Earlier timing and faster tempo of puberty have been associated with altered brain development and increased mental health symptoms in adolescents, particularly females. However, the role of oestradiol (E2) in these associations is unclear.

**Methods:** Using longitudinal data from the US-based Adolescent Brain Cognitive Development^SM^ Study (ABCD Study*®)*, we investigated whether, in females (N ∼ 3k), E2 timing (at age 10) and tempo (rate of change from age 10 to 12) were prospectively associated with mental health symptoms at age 13 via structural brain development from age 10 to 12. Linear mixed-effects models and Bayesian mediation models were fitted to investigate the aims of the study.

**Results:** Findings showed that E2 timing was not associated with mental health symptoms. However, earlier E2 timing was associated with a greater reduction in total cortical volume, total surface area, and surface area in the superior and middle temporal cortex over time. Further, a faster E2 tempo was associated with an increase in mental health symptoms, and this association was mediated by a faster reduction in total cortical volume and total surface area over time.

**Conclusion:** Findings suggest that earlier E2 timing and faster E2 tempo contribute to accelerated development of gray matter structure in adolescent females, and for E2 tempo, such associated brain changes may partly contribute to increased mental health risk.

## 1. INTRODUCTION

Puberty brings numerous biological and psychological changes, and individual differences in both the timing of pubertal events and the progression (or tempo) of pubertal changes have been linked to mental health risks, particularly in females. Early pubertal timing in females has consistently been associated with increased risk of internalising, externalising, and other psychopathologies (e.g., antisocial behaviour, substance abuse, and eating pathologies; Copeland et al., 2019; Horvath et al., 2020; Mendle et al., 2007; Ullsperger & Nikolas, 2017). While pubertal tempo is less studied, emerging evidence suggests faster tempo may also heighten psychopathology risk in female adolescents (Deardorff et al., 2021; Kowalski et al., 2021; Marceau et al., 2011; Nolen-Hoeksema & Girgus, 1994). Most research to date has relied on physical markers of pubertal development (e.g., Marshall & Tanner, 1969, 1970; Petersen et al., 1988; Platt et al., 2017), while the role of hormonal changes remains less clear. Given that the rise in hormones often precedes physical signs of puberty (Deardorff et al., 2021), examining how the timing and tempo of hormone exposure influence mental health may be important for prevention strategies. Additionally, examining their impact on neurobiological development could clarify mechanisms underlying adolescent mental health risk.

Among females, oestradiol (E2, the strongest of the naturally occurring oestrogens) plays a key role in puberty-related physical changes (e.g., breast development) and has been consistently associated with mental health in adult females, particularly during transitional stages of life, such as menopause and postpartum (Hara et al., 2015; Navarro-Pardo et al., 2018; Rieder et al., 2020). However, research in adolescent females is limited and mixed, with studies reporting positive, negative, or no effects of E2 on mental health (including internalising and externalising symptoms; Luo et al., 2024). Out of these, only a few studies controlled for age (and thus measured relative E2 timing), and most used cross-sectional designs and small sample sizes. Notably, no studies, to our knowledge, have investigated the effects of E2 change over time (i.e., tempo) on mental health. Thus, the effects of E2 timing and tempo on the mental health of adolescent females remain largely unexplored.

A possible mechanism by which pubertal timing and tempo contribute to various mental health could be via effects on the developing brain. E2 levels influence the developing brain through E2 receptors (Bixo et al., 1986, 1995a, 1997; Koss et al., 2015; Sisk & Zehr, 2005), and changes in E2-sensitive brain regions have been linked to adolescent mental health (Frodl & Skokauskas, 2012; Holtzman et al., 2013). There is evidence that pubertal processes (both physical and hormonal changes) affect brain development beyond chronological age (Dehestani et al., 2023; Vijayakumar et al., 2021), with early timing and faster tempo of physical pubertal changes related to accelerated patterns of normative development in brain structure over time—such as greater reductions in gray matter (GM) and increased white matter (WM) integrity (Beck et al., 2023; Chahal et al., 2018; Herting et al., 2014).

However, research on the effects of E2 timing and tempo on brain development is limited and inconsistent. For instance, higher E2 levels for one’s age (i.e., earlier timing of E2 exposure) have been associated with both larger and smaller frontal and temporal, smaller parietal and larger middle occipital GM densities (Brouwer et al., 2015; Peper et al., 2009), a smaller volume of the anterior cingulate cortex (Koolschijn et al., 2014) and lesser WM integrity (i.e., fractional anisotropy; FA) of the superior longitudinal fasciculus and angular gyrus (Herting et al., 2012). However, some studies found no significant association between E2 timing and WM FA among females (Peper et al., 2014). Longitudinal studies have reported that higher E2 levels for one’s age predict increases in FA in the uncinate fasciculus, whole-brain WM volume and amygdala volume over time (Herting et al., 2012). Regarding E2 tempo, the only study, to our knowledge, that has investigated associations between E2 changes over time and brain development in adolescent females found that a faster increase in E2 levels over two years was associated with a faster reduction in surface area of the middle temporal lobe (Herting et al., 2015). Notably, none of these studies investigated whether E2 timing or tempo was indirectly associated with mental health via brain changes.

This study aimed to longitudinally explore the total and indirect relationship between E2 timing and tempo and mental health symptoms during early-to mid-adolescence. In order to gain a comprehensive understanding of the effect of E2 timing/tempo on brain development, prior to testing indirect effects, we first examined associations between E2 timing/tempo and structural brain development across early adolescence. We then tested whether these brain changes mediated associations between E2 timing/tempo and mental health symptoms (refer to Figure 1). Based on existing studies of physical pubertal maturation and preliminary findings of E2 timing/tempo, we hypothesised that earlier E2 timing and faster E2 tempo would be associated with increased mental health symptoms, indirectly via an accelerated trajectory of GM and WM development. Given E2’s potentially broad influence on the brain (Goddings et al., 2019; Luo et al., 2024), we investigated both global and regional brain measures. We did not have specific hypotheses about brain regions or types of mental health symptoms that would be implicated.

**Figure 1.**
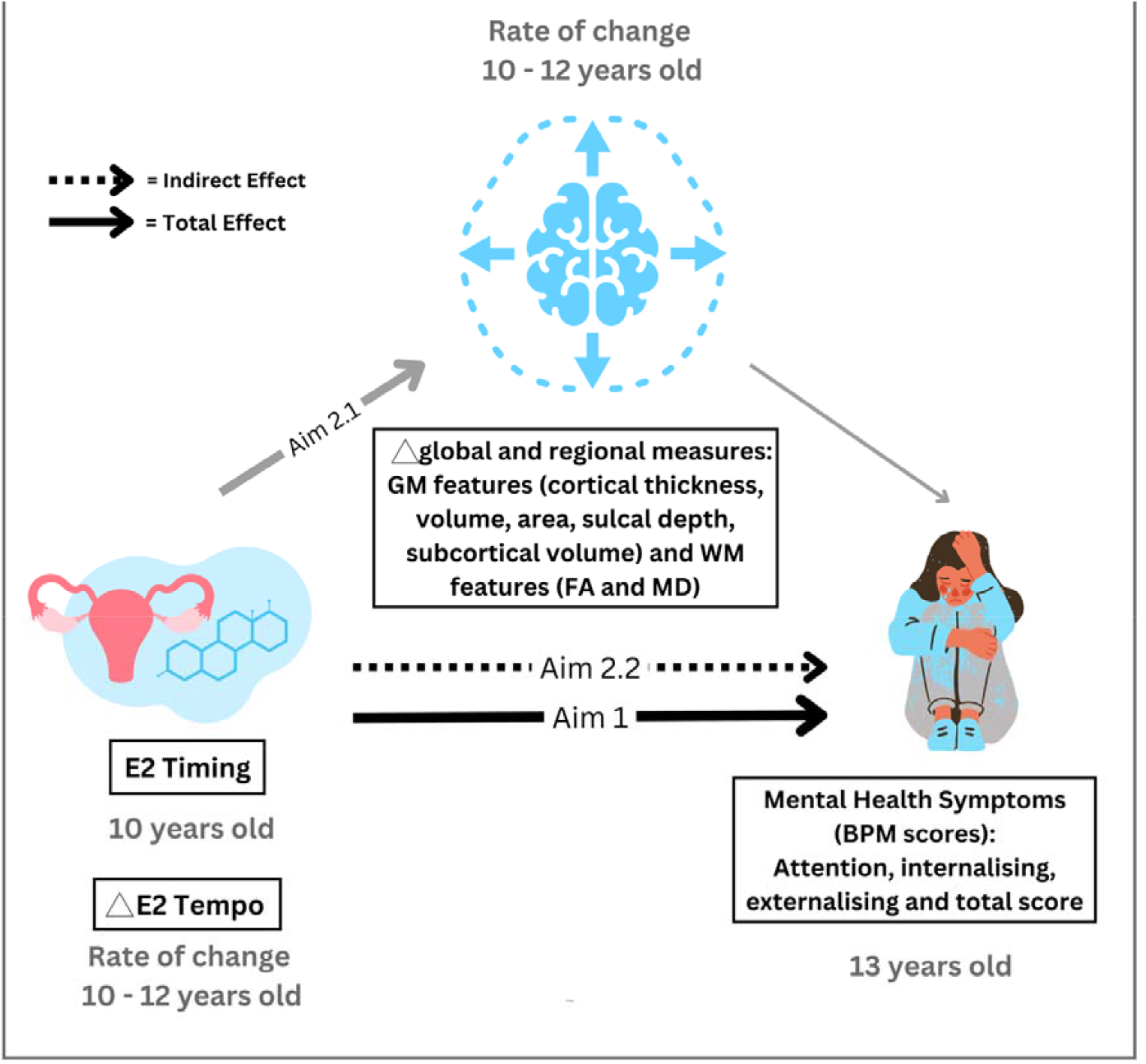
A Mediation model: E2 Timing and E2 tempo are the predictors, brain features are the mediators (Δ represents the rate of change), and mental health symptoms are the outcome. Mentioned age is the mean age of each group

Figure 1. : Bold arrow represents aim 1 of the study – the total effect of E2 timing and tempo and mental health symptoms. Grey and bold arrow represent aim 2.1 of the study – an association between E2 timing and tempo and brain structure. Dashed arrow represents aim 2.2 of the study, the indirect effect of E2 Timing and E2 tempo on mental health symptoms via brain changes. Mentioned age is the mean age at each wave. Δrepresents the rate of change.

## 2. Material and methods

Study hypotheses and analyses were preregistered (https://doi.org/10.17605/OSF.IO/7SNB4). Any changes from the preregistered plan are described below.

### 2.1 Participants

This study utilised data from the Adolescent Brain Cognitive DevelopmentSM Study (ABCD Study*®;* open-science dataset version 5.1; released 2024), a longitudinal study of adolescent development that includes data from 22 sites across the United States (www.ABCDStudy.org). For recruitment details, see Garavan et al., 2018. For the current study, we used structural and diffusion Magnetic Resonance Imaging (MRI) and E2 data from baseline (age 9-11 years) and Year 2 (age 11-14 years) for female participants. Parent-reported mental health symptoms at baseline and adolescent self-reports at Year 3 (age 11– 15 years) were also included.

Sample sizes for Aim 1 and Aim 2 varied based on available data (pairwise deletion was used). We excluded those taking oral contraceptives (N = 22). Imaging data were filtered using ABCD study recommendations (abcd_imgincl01) for T1-weighted structural MRI (sMRI) and diffusion-weighted MRI (dMRI). See Table 1 and Supplementary Figure 1 for details on sub-analysis sample sizes.

**Table 1:**
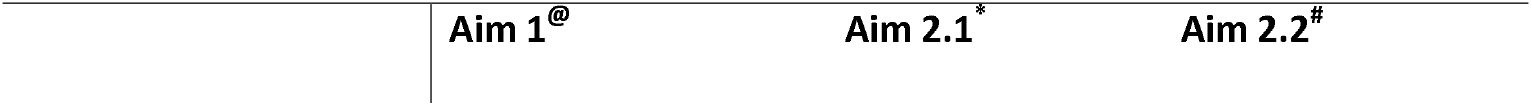
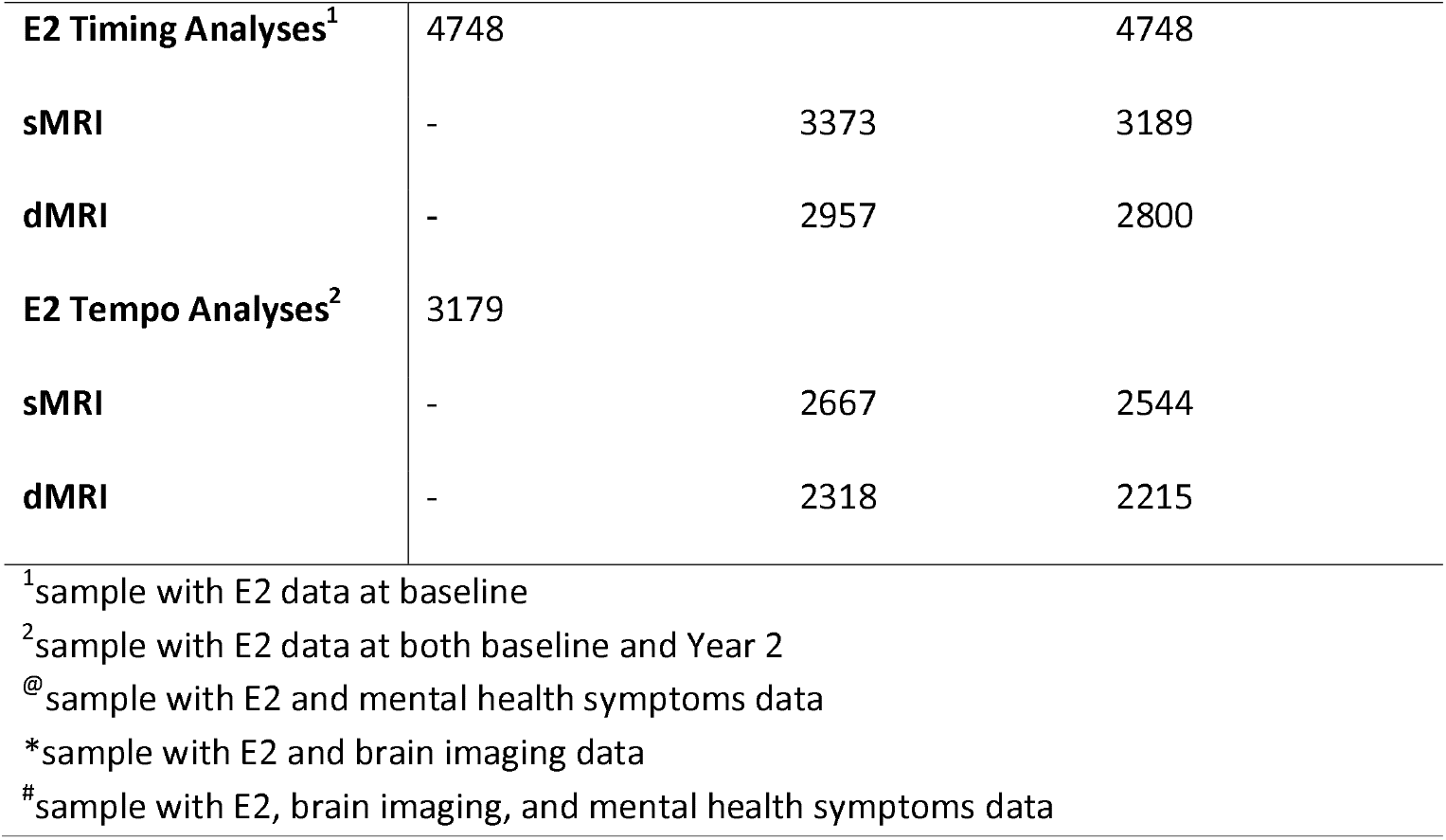
Sample sizes (N’s) used across different analyses.

### 2.2 Measures

#### E2 Hormone

E2 levels were measured from saliva collected on the MRI assessment day, following the protocol by Granger et al., 2012. Participants refrained from eating/drinking for at least 1 hour before collection. Samples were either frozen (−80°C to -20°C) or stored in a cooler (4°C) before being shipped on dry ice to Salimetrics (Carlsbad, CA) for ELISA-based assay in duplicate. E2 levels were cleaned and averaged from the duplicate assays according to the protocol by Herting et al., 2021.

#### Neuroimaging

Imaging data were collected using three different 3T MRI scanner platforms (*Siemens Prisma, General Electric (GE) 750 and Philips*). Further details on scanning parameters and imaging processing have been documented by the ABCD study investigators (Casey et al., 2018; Hagler et al., 2019).

For sMRI, cortical regions were parcellated using the FreeSurfer v7.1.1 Desikan atlas (Dale et al., 1999; Fischl et al., 1999), which has 34 cortical regions per hemisphere. Four cortical metrics, thickness, surface area, sulcal depth, and volume were examined. Seven subcortical regions were parcellated using FreeSurfer’s subcortical segmentation (Fischl et al., 2002). For the current analyses, given no hypotheses of lateralised effects, left and right regions were averaged across both hemispheres. However, hemisphere-specific analyses are reported in *Supplementary Materials*. Global measures of mean cortical thickness, total surface area, sulcal depth, and total cortical and subcortical volume, were also investigated.

From dMRI, 17 WM fibre tracts across the whole brain were parcellated using the AtlasTrack atlas (Dale et al., 1999; Fischl et al., 1999). FA and mean diffusivity (MD) were computed within these parcellated tracts (Hagler et al., 2019), averaging values across hemispheres. Only inner shell dMRI measures were included in this analysis (i.e., b = 500 & b = 1000 were included for tensor fit).

#### Multi-site scanner effects

To control for variance due to different scanners, a longitudinal harmonisation technique was implemented using the *LongComBat* package in R (Beer et al., 2020). This technique was implemented separately for different analyses, with the predictors from each analysis included in the harmonisation model.

#### Mental Health Symptoms

Mental health was assessed using the adolescent-reported Brief Problem Monitor (BPM; Achenbach et al., 2011) at Year 3. Three subscales (attention, internalising, externalising) in addition to a “total symptoms” score, were examined.

#### Covariates

For analyses involving mental health symptoms, we included scores from corresponding scales from the parent-reported Child Behaviour Checklist (CBCL; (Hagler et al., 2009) at baseline as covariates, as the BPM was not administered at baseline. Socio-demographic characteristics were included as covariates in sensitivity analyses, including BMI z-score (calculated according to the CDC 2000 Growth Charts; Gray et al., 2020), race/ethnicity, and SES (highest parental education and combined family income). See Supplementary Table 2 for further details.

### 2.3 Statistical analysis

#### E2 Timing

We calculated an individual’s E2 timing as the residual from regressing E2 on age at baseline:

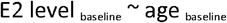

Residual = actual E2 level – predicted E2 level

#### E2 Tempo

E2 tempo was calculated as the rate of change in E2 levels from baseline to two-year follow-up:

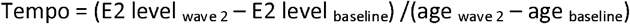

#### Brain development

Brain development was operationalised as the rate of change in brain structure from baseline to the two-year follow-up:

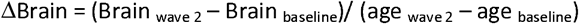

##### Aim 1: Relationship between E2 timing/tempo and mental health symptoms

Linear mixed effect (LME) models (R package; nlme) were fitted to compute the association between E2 timing and tempo and mental health symptoms. Separate models were run for each predictor (E2 timing or tempo) and each symptom, totalling eight models. Respective parent-reported baseline mental health symptoms were covariates in each model. Site, scanner type and family identity were included as nested random effects.

##### Aim 2: Indirect relationship between E2 timing/tempo and mental health symptoms via brain development

###### Aim 2.1 Relationship between E2 timing/tempo and brain development

The association between E2 timing and tempo and brain development was computed using separate LME models with the same random nested effects as the model for Aim 1. Separate models were run for each brain feature, N _models_ = 184 (148 GM and 36 WM).

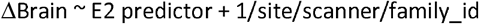

Here, E2 timing and E2 tempo were used as predictors in separate models. 1/site/scanner/family_id represents nested random effects

FDR correction (p_FDR_ < 0.05) was applied for the number of brain ROIs for each of the structural metrics (i.e., 34 tests for each cortical metric, 7 tests for subcortical volumes, 17 tests for each WM microstructure property).

###### Aim 2.2 Mediating role of brain development

Bayesian mediation models (brms, R) examined whether brain development mediated the relationship between E2 timing/tempo and mental health symptoms. The corresponding parent-reported baseline symptom was included as a covariate, and random effects matched the LME models. Indirect effects were computed using “bayestestR”. Mediation models were only run for significant E2 timing/tempo to brain associations and total effect models from previous analyses.

### 2.4. Replicability and sensitivity analyses

To assess replicability, we conducted a split-half analysis for aim 1 and aim 2, ensuring demographic matching (BMI, race/ethnicity, parent education, and income) across both the split-half samples via t-tests. We then repeated the main analyses on each split sample and examined whether effects for significant findings from the full sample remained consistent in both splits.

As per our pre-registration and consistent with other studies on E2 levels and brain development (Brouwer et al., 2015; Herting et al., 2014), we present the results from models without any socio-demographic covariates included in models. In sensitivity analyses, we re-ran models a) including sociodemographic covariates (i.e., BMI, race/ethnicity, parent education, and parent combined income) and regressing out saliva-related saliva covariates (e.g., caffeine intake, physical activity, and collection time) from E2 levels prior to calculating E2 timing/tempo, and b) winsorising statistical outliers (mean +/-3SD). Results of sensitivity analyses are in Supplementary Materials.

### 2.5. Exploratory analyses

Although not pre-registered, we conducted an additional analysis to investigate the total and indirect effects of the E2 timing and tempo interaction on mental health symptoms, given that an interactive effect of pubertal timing and tempo on mental health has been found in prior literature (Mendle et al., 2010).

## 3. RESULTS

### 2.1. Descriptives

Demographic and pubertal characteristics of participants are presented in Table 2, while the distribution of E2 timing and tempo values is shown in Supplementary Figure 1. E2 timing and tempo were negatively correlated (r = -0.61, p < 0.001).

**Table 2:**
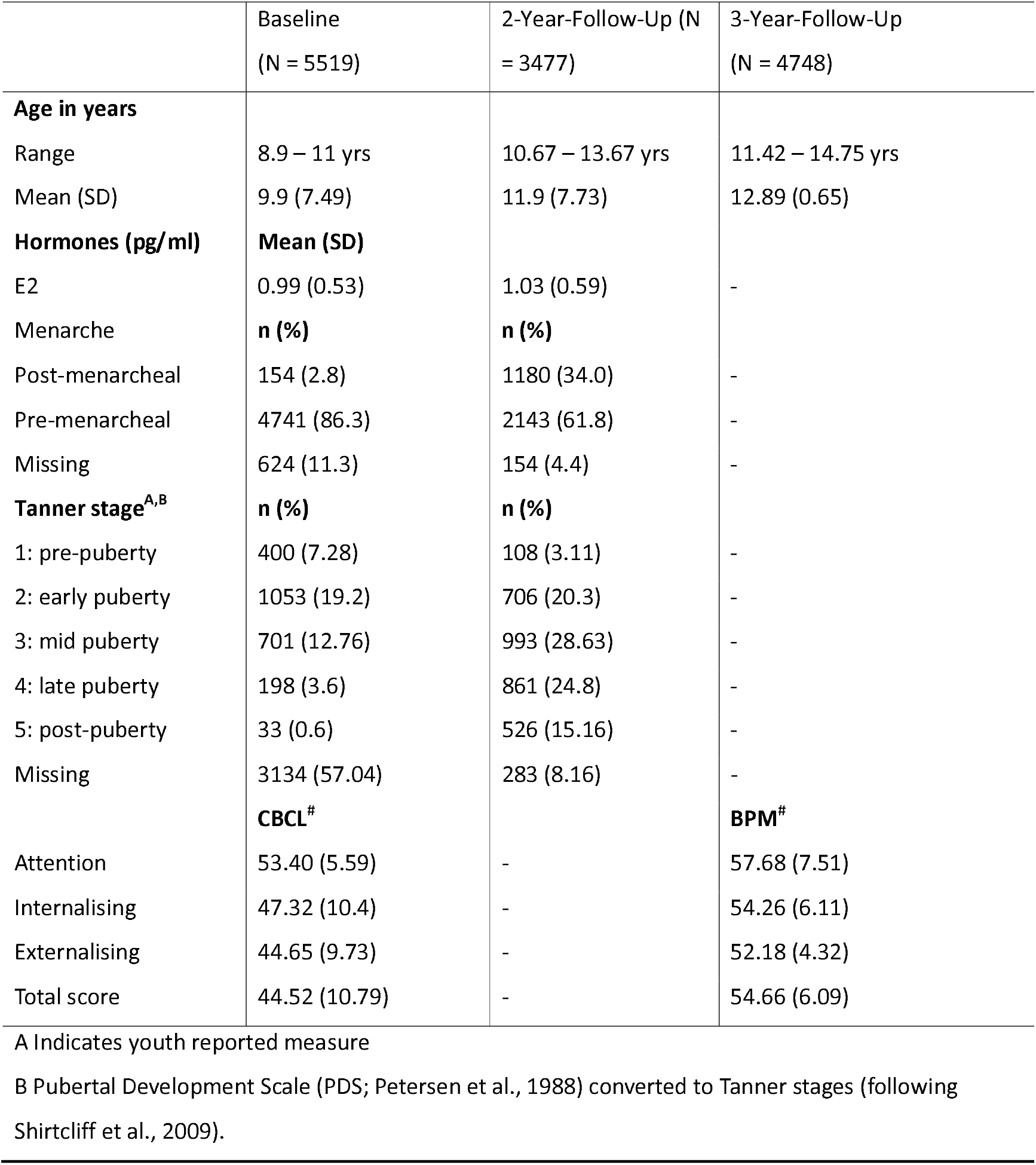
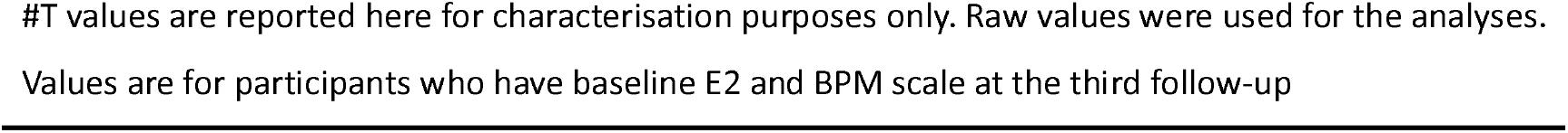
Descriptives at Each Wave.

T-scores for parent-reported CBCL and adolescent-reported BPM subscales are also in Table 2. Covariate details, along with their correlations with mental health symptoms, and E2 timing and tempo, are provided in Supplementary Table 2 and Figure 2.

**Figure 2.**
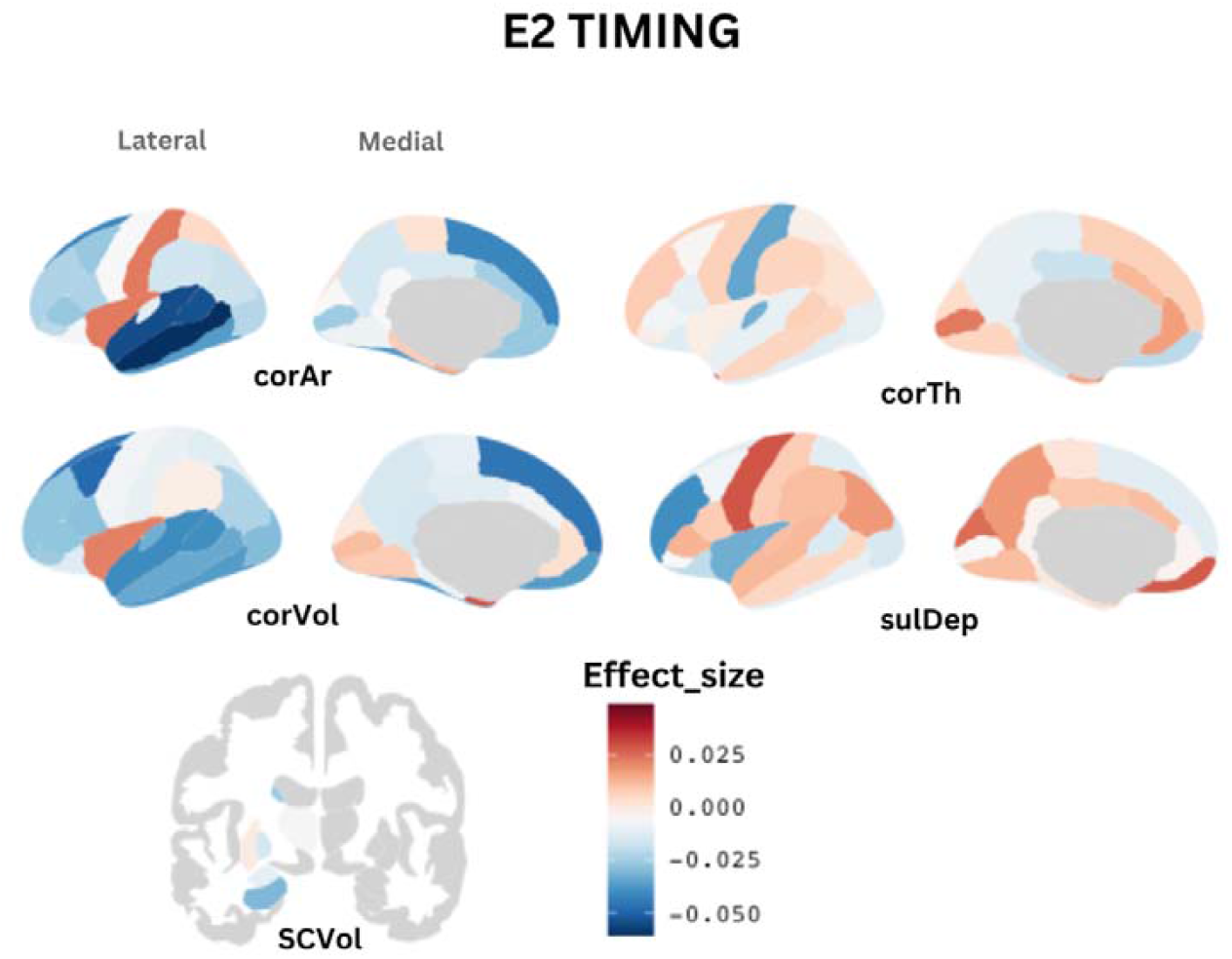
Effect size (Cohen’s d) of the association between E2 timing and each of the structural matrices across the whole brain. Abbv: corAr = cortical area; corTh = cortical thickness; corVol = cortical volume; sulDep: sulcal depth; SCVol = subcortical volume

Analyses revealed significant differences in demographic characteristics (race/ethnicity and socioeconomic status) between included and excluded participants. Excluded individuals were those with missing E2 data at baseline or follow-up, or those with poor-quality imaging data, even if hormone data were available (see Supplementary Table 3).

### 3.2. E2 Timing/tempo and mental health symptoms

E2 timing was not significantly associated with any of the mental health symptoms at the three-year follow-up. E2 tempo was significantly positively associated with attention (B = 4.85, p = 0.011, Cohen’s d = 0.05), internalising (B = 3.99, p = 0.028, Cohen’s d =0.044), and total symptoms (B = 11.93, p = 0.004, Cohen’s d = 0.056).

### 3.3 E2 Timing and Brain Development

In the whole sample, earlier E2 timing predicted greater reductions in global and regional brain structure over time. Specifically, it was associated with reductions in total cortical volume (B = -26.17, p = 0.018, Cohen’s d = -0.041), total surface area (B = -6.57, p = 0.016, Cohen’s d = -0.042), and surface area in the superior temporal cortex (B = -0.315, p_FDR_ = 0.022, Cohen’s d = -0.055), banks of the superior temporal sulcus (B = -0.122, p_FDR_ = 0.022, Cohen’s d = -0.054), and middle temporal cortex (B = -0.331, p_FDR_ = 0.042, Cohen’s d = - 0.056). However, no regional findings remained significant after controlling for total surface area, suggesting that effects may not be regionally specific.

No significant associations were found between E2 timing and FA/MD in any white matter tracts. Coefficients for GM and WM tracts are shown in Figures 2 and 4A, with effect sizes and p-values detailed in Supplementary Tables 4-6.

### 3.3 E2 Tempo and Brain Development

Faster E2 tempo was significantly associated with a greater reduction in total cortical volume (B = -1236.27, p = 0.004, Cohen’s d = -0.055) and total surface area (B = -242.28, p = 0.027, Cohen’s d = -0.043) over two years.

No significant associations were found between E2 tempo and FA/MD in any white matter tracts. Coefficient values for GM and WM are shown in Figures 3 and 4B, with effect sizes and p-values detailed in Supplementary Tables 7-9.

**Figure 3.**
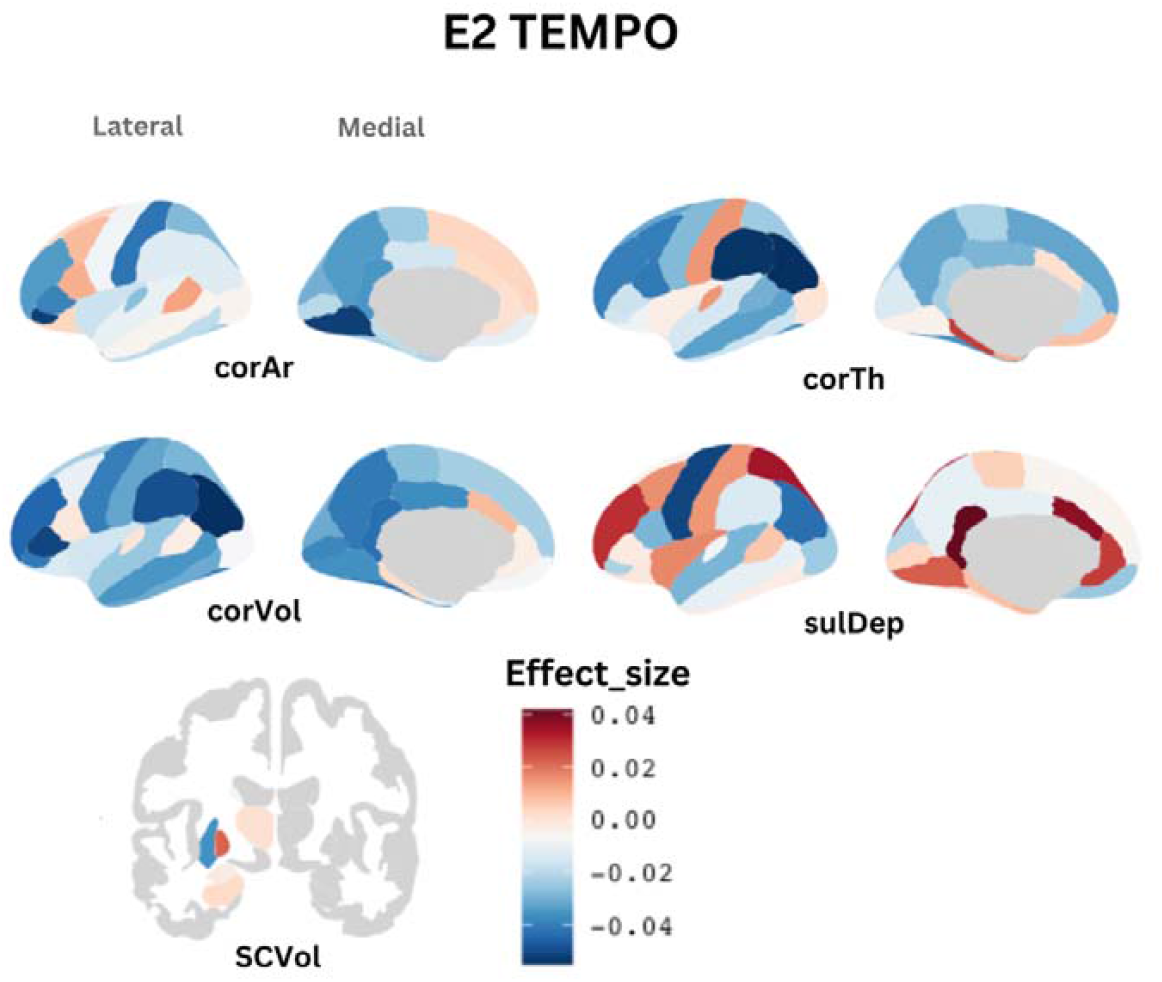
Effect size (Cohen’s d) of the association between E2 tempo and each of the structural matrices across the whole brain. Abbv: corAr = cortical area; corTh = cortical thickness; corVol = cortical volume; sulDep: sulcal depth; SCVol = subcortical volume

**Figure 4.**
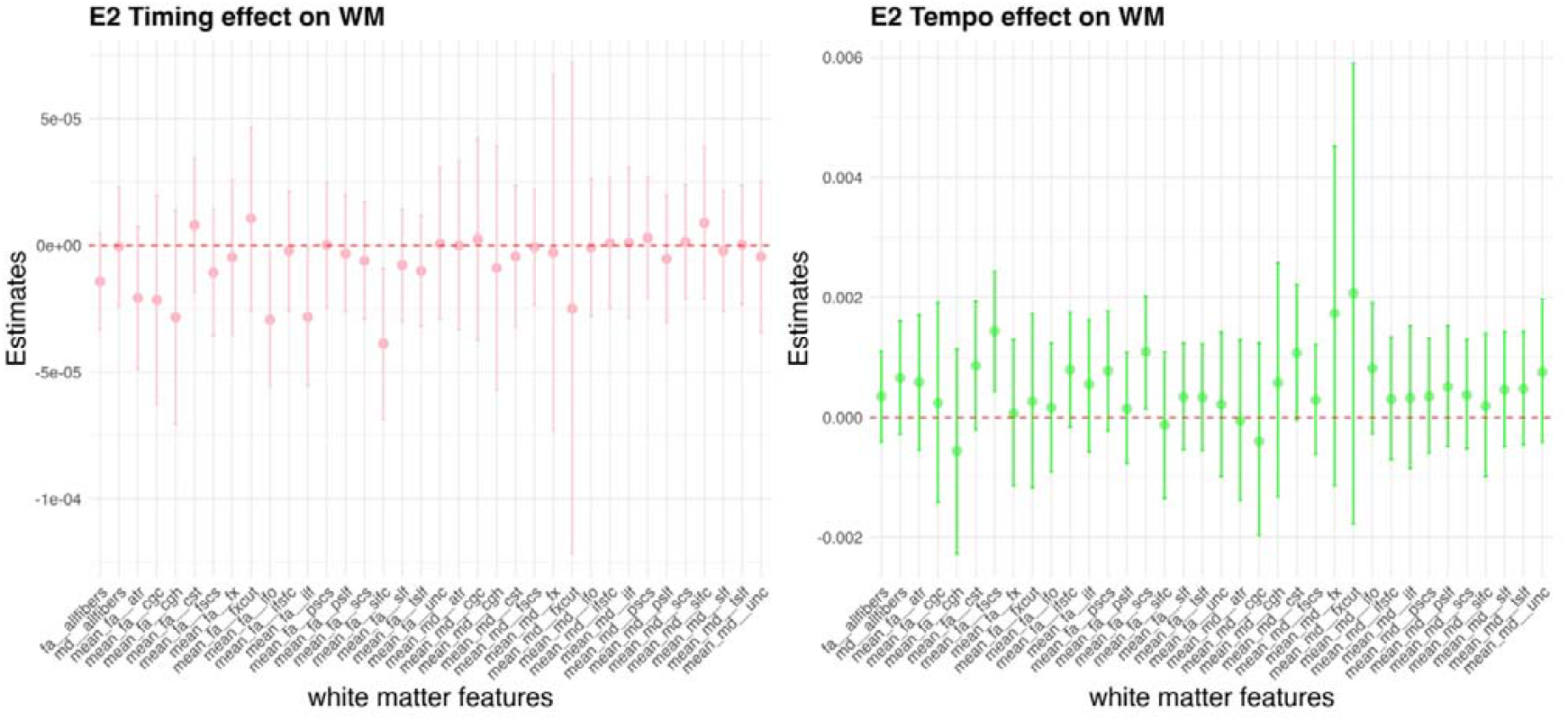
Beta coefficient (with ± 95 CI) of the association between A) E2 Timing and white matter FA and MD, and B) E2 Tempo and WM FA and MD. Red dashed line represents the zero point. FA/MD measures that do not cross the red dashed line were significant before FDR-correction

### 3.4 Mediation Analysis

Mediation analyses examined whether changes in total cortical volume and surface area mediated the association between E2 tempo and mental health symptoms. Using Bayesian models, a greater reduction in total cortical volume significantly mediated the relationship between faster E2 tempo and total symptoms (B = 0.524, CI_95_ = 0.043, 1.354; Figure 5C). Additionally, a greater reduction in total surface area mediated the association between E2 tempo and attention symptoms (B = 0.283, CI_95_ = 0.044, 0.676; Figure 5A) as well as total symptoms (B = 0.606, CI_95_ = 0.080, 1.483; Figure 5B).

**Figure 5.**
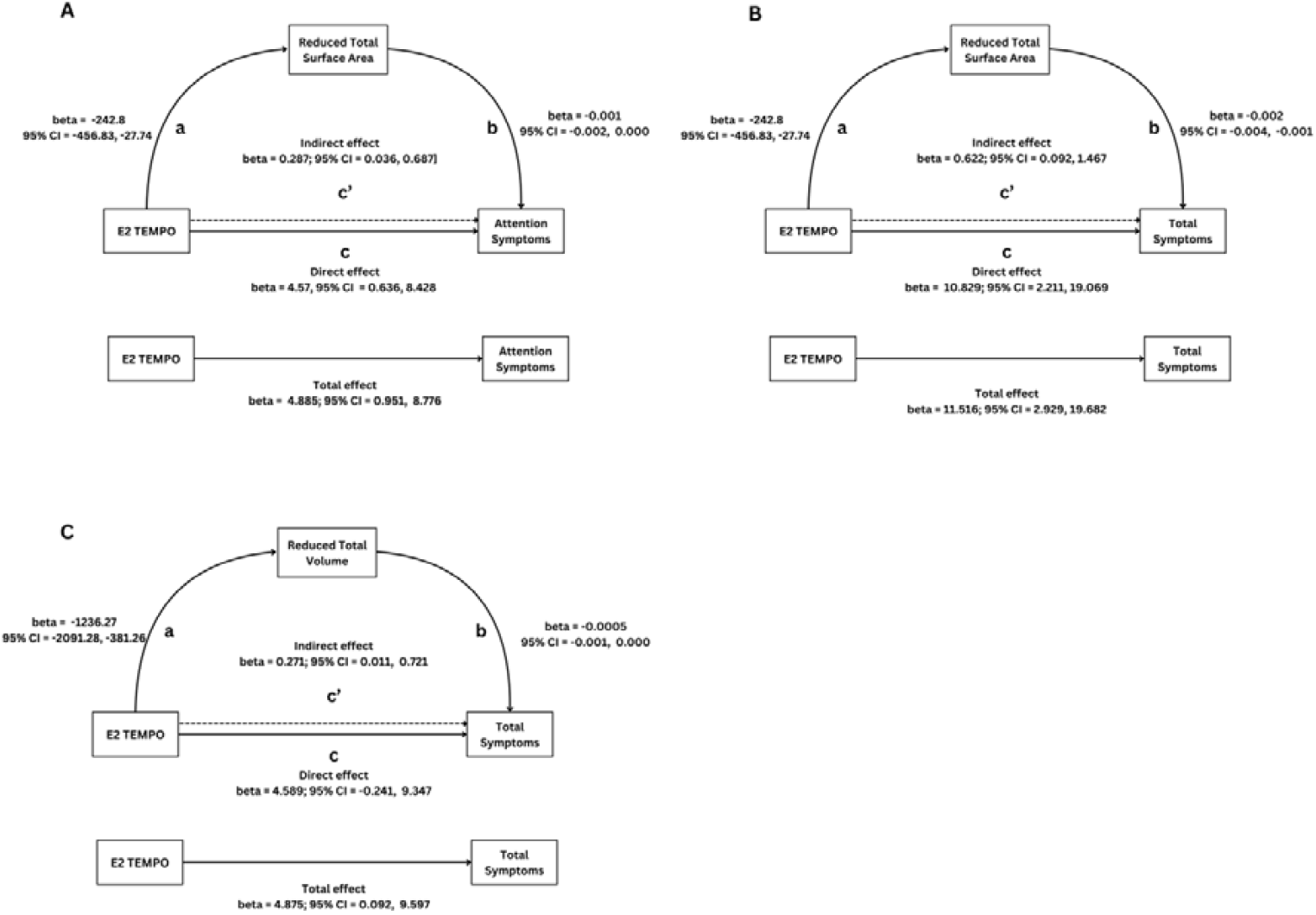
Indirect effect of E2 tempo on A) attention symptoms via greater reductions in surface area, B) total symptoms via greater reductions in surface area, and C) total symptoms via greater reductions in cortical volume.

### 3.5 Replication and Sensitivity Analyses

Effect sizes were comparable for significant models in the two split-half samples. However, effects were not always significant in each split-half sample (see *Supplementary Materials*).

Most of the significant findings remained significant after accounting for saliva covariates and including SES and race/ethnicity in models. However, no indirect association between E2 tempo and attention/total mental health symptoms survived when the BMIz score was included as a covariate. With winsorised data, the total effects of E2 tempo on mental health symptoms remained significant. However, no indirect effects remained significant.

### 3.6 Exploratory analyses

The interaction between E2 Timing and E2 Tempo was not significantly associated with any brain or mental health measures.

## DISCUSSION

In a large U.S.-based longitudinal sample of 9 – 15-year-old females, we found that faster E2 tempo was associated with an increase in attention, internalising and total mental health symptoms. This association was mediated via greater reductions in total volume and total surface area. While E2 timing showed no association with mental health symptoms, earlier E2 timing was associated with greater reductions in global and temporal GM.

We did not find the hypothesised association between E2 timing and mental health symptoms, suggesting that an early rise in E2 levels during late childhood may not predict emerging psychopathology two years later. While this contradicts our hypothesis, it aligns with previous studies that found no direct link between E2 levels and internalising symptoms, although these studies did not account for age (Barendse et al., 2022; Copeland et al., 2019; Hazell et al., 2023). In contrast, studies assessing timing of physical pubertal development have more consistently linked early maturation—particularly in females—to increased psychopathology risk (Copeland et al., 2019; Horvath et al., 2020; Ullsperger & Nikolas, 2017). As such, perhaps early maturation is more meaningful for psychopathology risk when assessed by physical changes rather than the E2 levels. This aligns with the “maturation disparity hypothesis,” which proposes that early maturers experience psychological challenges due to an asynchrony between physical and emotional development (Ge & Natsuaki, 2009), increasing their risk for mental health symptoms.

Consistent with hypotheses, a faster increase in E2 levels across two years was linked to higher attention, internalising, and total mental health symptoms. This aligns with the ‘stressful change’ and ‘maturation compression’ hypotheses, which suggest that rapid pubertal transitions shorten the time for developing adequate coping skills in response to the biological and psychosocial challenges of puberty, thereby heightening their vulnerability to psychopathology (Ge et al., 2001; Mendle et al., 2010). Extending these frameworks to hormonal changes, a faster increase in E2 may create a mismatch between biological maturation and cognitive, social, and emotional adaptation. Therefore, this suggests that rapid increases in E2 levels may leave children psychologically less equipped to navigate the demands of puberty and thus place them at greater risk for mental ill-health.

As hypothesised, higher E2 levels for one’s age in late childhood and a more rapid E2 increase with age were associated with decreased GM structure across early adolescence. This likely reflects accelerated maturation, as both surface area and volume typically decline during this period (Mills et al., 2021; see Supplementary Figure 3 for age-related effects in the sample). While no prior longitudinal studies have examined E2 timing and brain development, some cross-sectional research suggests the link between higher E2 levels and lower global GM volume in females (Herting et al., 2014; Peper et al., 2009). Additionally, earlier physical maturation has been associated with accelerated patterns of brain structure maturation (Goddings et al., 2019; Holm et al., 2023; Vijayakumar et al., 2018), including findings from the ABCD dataset (Beck et al., 2023; Dehestani et al., 2023; MacSweeney et al., 2023). Whether, collectively, these findings support a direct hormonal influence (as opposed to an effect reflecting social or cultural responses to visible physical changes associated with puberty) on brain development is unclear. While Dehestani et al. (2023) found that physical maturation had a stronger impact on brain development than hormonal levels, they did not examine E2. Given the weak correlations between E2 and physical development in ABCD participants (Supplementary Table 1), our findings suggest that early E2 increases may shape brain development even before overt physical changes become apparent.

Regionally, earlier E2 timing was associated with reduced surface area in the superior and middle temporal cortices (MTC) and the banks of the superior temporal sulcus, aligning with prior findings of smaller MTC GM density with higher E2 (Peper et al., 2009) and MTC thinning with faster E2 increases (Herting et al., 2015). Furthermore, physically more mature girls have shown smaller MTC volumes (Bramen et al., 2011). The temporal cortex has a high density of E2 receptors (Bixo et al., 1995b), and early physically maturing females with higher E2 levels exhibit altered temporal region function during social-emotional processing (Goddings et al., 2012; Klapwijk et al., 2013; Sumner et al., 2018). While speculative, our findings suggest neural reorganisation of regions supporting social-emotional processing in response to earlier E2 increases. However, these regional effects became non-significant after controlling for whole-brain surface area reductions, raising questions about whether they reflect local specificity or broader cortical changes. The field lacks clarity on the optimal approach for parsing regional from global longitudinal change (for a detailed discussion, see Vijayakumar et al., 2018), and as such, the regional effects seen here may reflect a particular involvement of the temporal cortex.

A faster increase in E2 levels (or tempo) was linked to greater reductions in global cortical volume and surface area, partially aligning with Herting et al. (2015), who found cortical thinning with faster E2 increases. This remains the only study to explore E2 tempo and structural brain development in adolescent females, though it did not examine global measures or cortical volume. Findings on pubertal tempo (based on physical changes) and brain development have also found faster tempo to be associated with greater cortical surface area reduction (Beck et al., 2023; Herting et al., 2015), with findings for other cortical metrics (i.e., thickness) inconsistent (Beck et al., 2023; Herting et al., 2015; Vijayakumar et al., 2021). Additionally, animal studies show that E2 increases impact synaptogenesis and synaptic plasticity in the female rat brain (Mukai et al., 2010). Thus, our findings may suggest that a faster increase in E2 influences synaptic plasticity, leading to a faster decrease in GM volume and surface area with age.

Notably, our findings further suggest that the link between faster E2 tempo and elevated mental health symptoms was mediated by accelerated reductions in whole-brain surface area and volume. Greater reductions in GM structure during adolescence may reflect atypical synaptic pruning processes, which have been associated with increased risk for psychopathology and disrupted functional connectivity, potentially impairing cognitive, social and emotional functioning (Anderson, 2011; Germann et al., 2021; Li et al., 2017; Shen et al., 2021). Taken together, a rapid rise in E2 levels from late childhood to early adolescence may contribute to neurobiological alterations that compromise the development of emotional and cognitive functioning, increasing vulnerability to mental health challenges in females. However, as our study captured E2 changes at only two time points during early adolescence, it provides a limited snapshot rather than a comprehensive trajectory across the full pubertal period. Future research is needed to examine these hormonal, neurobiological, and psychosocial pathways over the entire course of puberty.

There were no significant associations between E2 timing/tempo and mental health symptoms via WM development. A lack of associations between E2 timing/tempo and WM contrasts with prior studies (Herting et al., 2012; Ho et al., 2020). One explanation is that E2 timing/tempo may more strongly influence mental health via GM rather than WM development. Alternatively, our WM measures—derived from diffusion tensor models—may be less reliable, as they fail to capture complex fiber architecture (Alexander et al., 2002; Frank, 2001; Tuch et al., 2002) compared to approaches like sparse fascicle models (SFM; (Rokem et al., 2015). Future studies should explore these associations using advanced WM metrics.

### Limitations

The current study has provided some of the first evidence of associations between E2 timing and tempo, brain development and mental health. Nevertheless, this study also has some limitations that need to be considered. First, not all findings remained significant after winsorising, although it is important to note that winsorising may introduce bias and is not considered an appropriate method for addressing outliers by some (Dyckman & Zeff, 2019) (hence why it was not employed in our primary analyses). In addition, effects did not persist when BMI was included. This is not surprising given the established link between BMI and puberty (Li et al., 2017; Reinehr & Roth, 2019). Since BMI and E2 were assessed at the same time, BMI likely accounted for relevant variance related to E2 timing and tempo. Further research is needed to clarify the overlapping and independent roles of E2 and BMI in shaping brain development and mental health.

Second, E2 levels fluctuate across the menstrual cycle, even before menarche (Legro et al., 2000). By follow-up, nearly 50% of the sample had reached menarche, but menstrual cycle phase at saliva collection was not recorded. This likely introduced noise into the E2 measures, contributing to the small effect sizes observed. Future studies should consider cycle phase, though this remains challenging given cycle irregularity in the early post-menarche years (Treloar et al., 1967).

Third, while effects were similar in split-half analysis, many were not significant, likely due to the small effect sizes (<0.1). While the small effect size could be due to noise in the E2 measure, it may be the case that effects are very small, meaning that clinical meaningfulness should be interpreted with caution.

While the ABCD study aimed to include a demographically diverse sample, those included in analyses (i.e., without missing data) had higher SES and were more likely to be White. Further, there is a relatively reduced representation of individuals with higher mental health symptoms, potentially contributing to smaller effect sizes. Future research should examine these associations in populations with more severe mental health symptoms to establish clinical relevance.

## Conclusion

In conclusion, the current study highlights significant effects of tempo of E2 changes on mental health symptoms in early adolescent females, highlighting the mediating role of structural brain development. Findings suggest that a greater increase in E2 levels during early adolescence may accelerate gray matter structure development, increasing the risk of psychopathology. However, limitations of this work underscore the need for future research to further examine how individual differences in E2 timing and tempo influence both neurodevelopmental trajectories and mental health symptoms across adolescence.

## DECLARATIONS

## Acknowledgements

MK is supported by the Melbourne Research Scholarship at the University of Melbourne. The authors have declared that they have no competing or potential conflicts of interest and have agreed to the submitted version.

## Data Sharing

This study used data from the Adolescent Brain Cognitive Development (ABCD) Study, a large-scale, longitudinal project tracking over 11,000 children (aged 9–10) across 10 years (Reinehr & Roth, 2019). Data are available via the NIMH Data Archive (NDA). Details on funding partners and study sites can be found at abcdstudy.org/federal-partners and abcdstudy.org/scientists/workgroups. ABCD consortium investigators designed and conducted the study but were not involved in this analysis or manuscript preparation. The views expressed here are solely those of the authors and do not necessarily reflect those of the NIH or ABCD consortium.

