## Supplementary Materials for "Linking Oestradiol Timing and Tempo, Brain Development, and Mental Health in Adolescent Females"

___________________________________________________________________________

### **Methods:**

##### Sample sizes (N’s) across different analyses

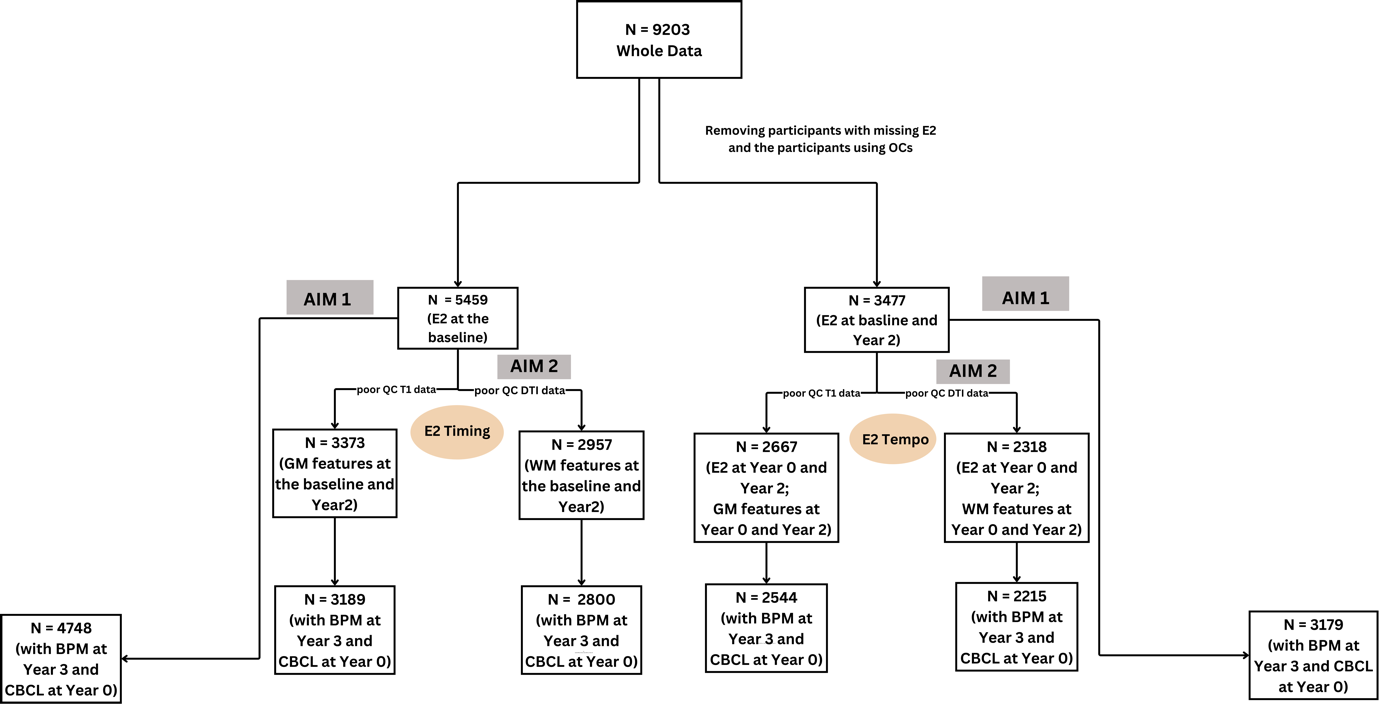

### **Results:**

#### 1. Descriptives:

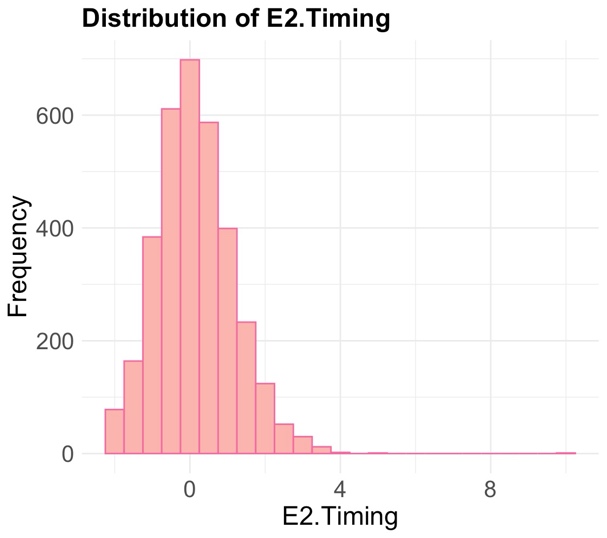

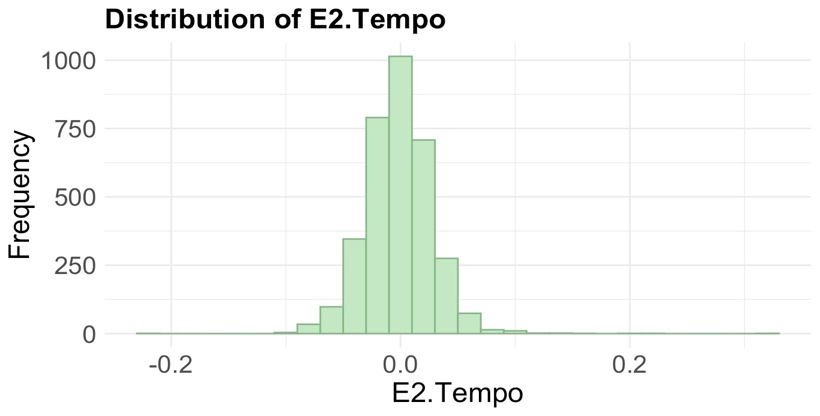

##### Figure 1: Frequency distribution of E2 Timing and E2 Tempo

##### Table 1: Correlation between E2 Timing and Tempo and puberty

|  | E2 Timing | E2 Tempo and baseline Tanner stages* | E2 Tempo and Wave 2 tanner stages* |
| --- | --- | --- | --- |
| Adrenal score | 0.07 (p = 3.309e-06) | 0.02 (p = 0.32) | 0.08 (p<0.001) |
| Gonadal score | 0.1 (p = 3.121e-07) | 0.07 (p = 0.01) | 0.07 (p=0.0003) |
| Body hair | 0.06 (p = 9.379e-06) | 0.018 (p = 0.32) | 0.06 (p = 0.002) |
| Skin changes | 0.05 (p=0.0005) | 0.03 (p = 0.16) | 0.07 (p<0.001) |
| height | 0.06 (p= 0.0002) | 0.003 (p = 0.89) | 0.06 (p < 0.002) |
| Breast development | 0.08 (p = 1.223e-07) | 0.05 (p = 0.008) | 0.04 (p = 0.03) |
| Menarche status (yes/no) | 0.075 (p= 1.437e-07) | 0.02 (p = 0.38) | 0.06 (p < 0.001) |

*Tanner stages were calculated using Pubertal Developmental Scale (PDS; Shirtcliff algorithm)

##### Table 2: Covariates at baseline (N = 5493)

| BMI (percentile) | **n (%)** |
| --- | --- |
| Underweight (<5^th^ percentile) | 232 (0.04) |
| Normal weight (>5^th^ to <85^th^ ) | 3531 (64.28) |
| Overweight (>85^th^ to <95^th^ ) | 844 (0.15) |
| Obese (>=95^th^ ) | 886 (0.16) |
| Race/Ethnicity^a^ | **n (%)** |
| White | 2803 (51.03) |
| Black | 874 (15.9) |
| Hispanic | 1100 (20.02) |
| Asian | 128 (2.3) |
| Other | 588 (10.7) |
| Parent Education^a^ | **n (%)** |
| <HS | 397 (7.2) |
| HS/GED | 567 (10.32) |
| Some college | 872 (15.87) |
| Associate degree (occupational/academic program) | 390 (7.09) |
| Bachelor’s degree | 1547 (28.16) |
| Postgraduate degree | 1099 (20.00) |
| Higher Professional/Doctoral degree | 306 (5.57) |
| Missing | 7 (0.12) |
| Family combined income^a^ | **n (%)** |
| < $5000 | 188 (3.42) |
| $5,000-$11,999 | 207 (3.76) |
| $12,000-$15,999 | 127 (2.31) |
| $16,000-$24,999 | 225 (4.09) |
| $25,000-$34,999 | 330 (6.00) |
| $35,000-$49,999 | 435 (7.91) |
| $50,000-$74,999 | 674 (12.27) |
| $75,000-$99,999 | 746 (13.58) |
| $100,000-$199,999 | 1530 (27.85) |
| > $200,000 | 575 (10.46) |
| Do not know | 230 (4.18) |
| Missing | 226 (4.11) |

A Indicates parent-reported measure

Table 3: Correlations between E2 levels and saliva covariates at each wave

|  | Baseline | 2-Year-Follow-Up |
| --- | --- | --- |
| **E2 correlations** | r (p-value) | r (p-value) |
| Caffeine intake | 0.073 (< 0.001) | 0.14 (< 0.001) |
| Physical activity | 0.04 (< 0.001) | 0.03 (0.07) |
| Collection time (in minutes) | 0.114 (< 0.001) | 0.07 (< 0.001) |
| Collection duration (in minutes) | -0.018 (0.380) | -0.0278 (0.101) |

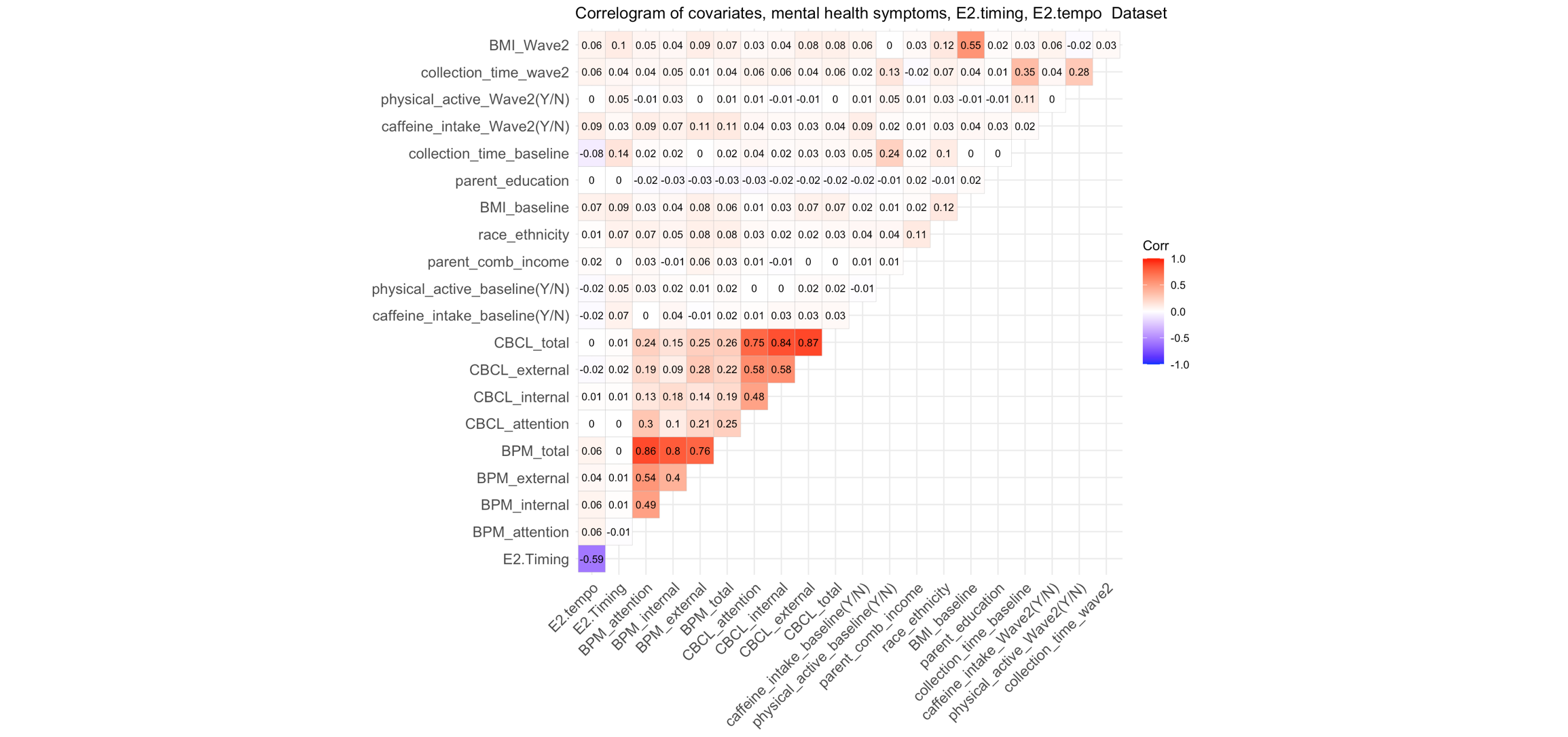

##### Figure 2: Correlation of E2 timing and E2 tempo with demographic variables and mental health symptoms

##### Table 4: Missing DataAnalyses for each sub-sample

All the missed data analyses were performed based on a comparison between the included individuals and excluded individuals (with missing E2 and imaging data, poor imaging data)

sMRI Timing

| **Variables** | **T-value (p)** |
| --- | --- |
| Race | -1.53 (0.126) |
| Parent combined income | -2.47 (0.013) |
| Parent education | 0.041 (0.967) |
| BMI- zscore | -0.758 (0.450) |

dMRI Timing

| **Variables** | **T-value (p)** |
| --- | --- |
| Race | -3.53 (0.0004168) |
| Parent combined income | -3.0949 (0.002) |
| Parent education | 0.60395 (0.545) |
| BMI - zscore | -0.190 (0.850) |

sMRI_Tempo

| **Variables** | **T-value (p)** |
| --- | --- |
| Race | 2.246 (0.024) |
| Parent combined income | -0.11963 (0.904) |
| Parent education | -0.833 (0.404) |
| BMI-zscore | 1.44 (0.149) |

dMRI_Tempo

| **Variables** | **T-value (p)** |
| --- | --- |
| Race | -0.38698 (0.698) |
| Parent combined income | -1.5459 (0.122) |
| Parent education | -0.23268 (0.816) |
| BMI-zscore | 2.3 (0.021) |

A)

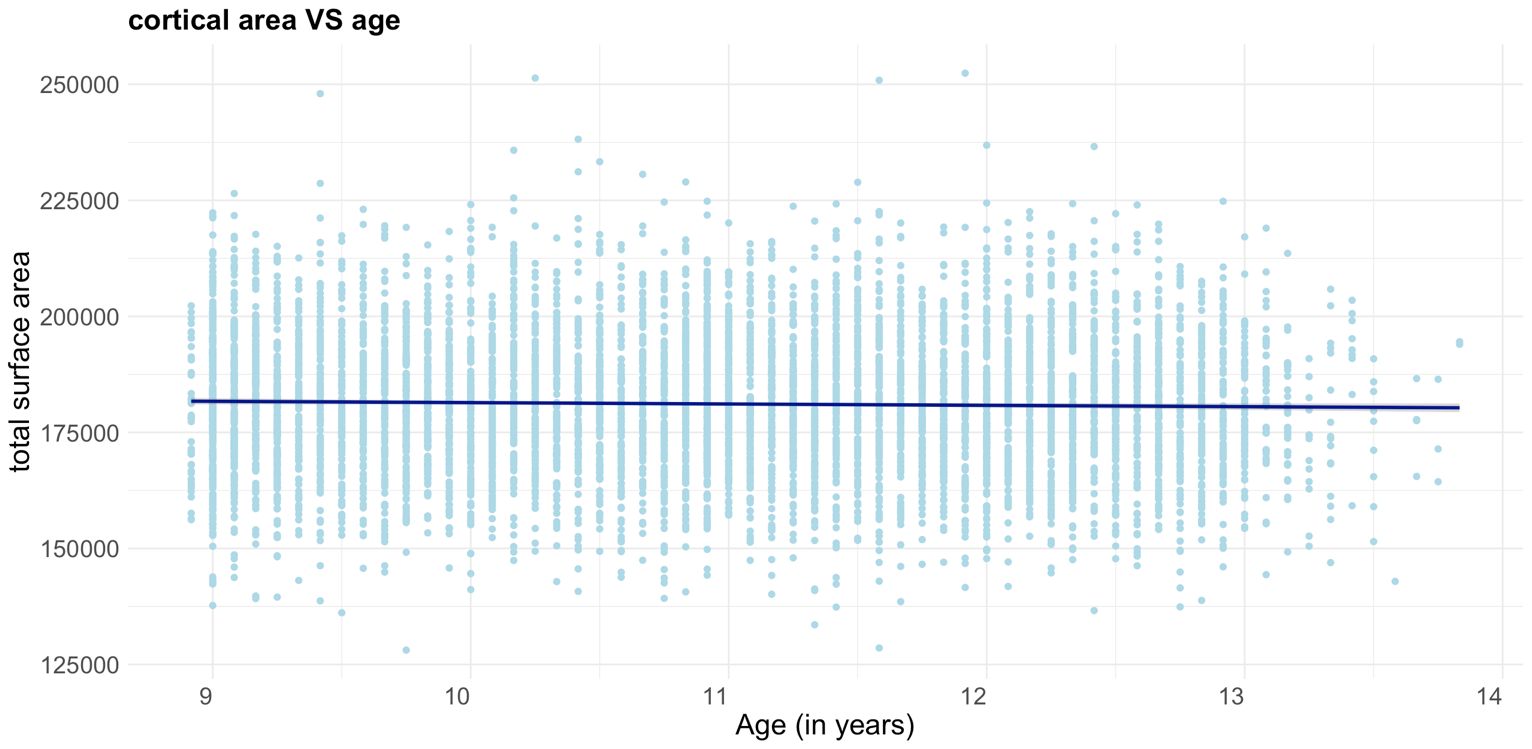

B)

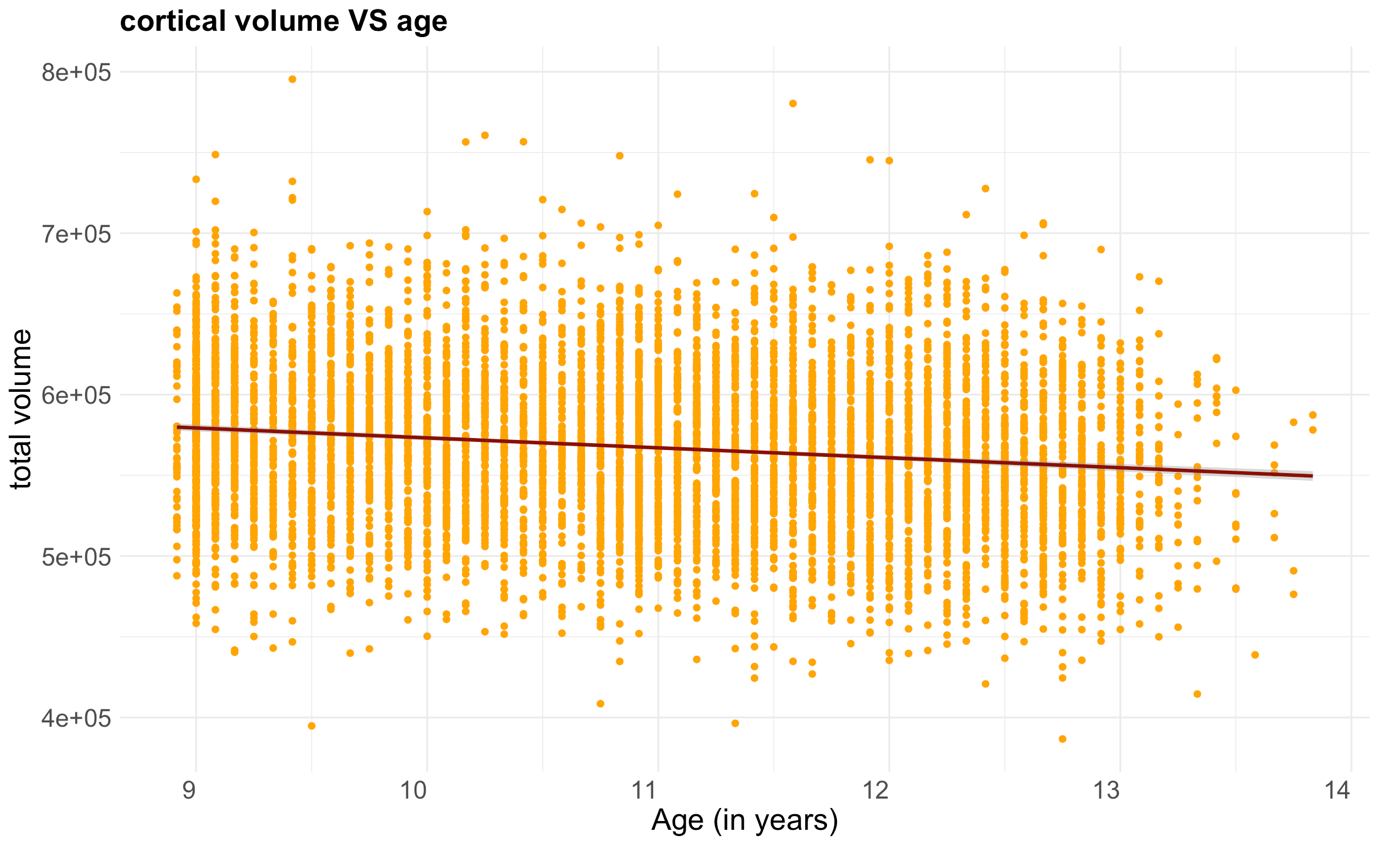

##### Figure 3: Distribution of A) total surface area and B) volume across age

#### 2. E2 timing, tempo, and brain structure

*mentioned p-values in all the tables are before the FDR-correction

##### *E2 Timing and cortical GM measures across the whole brain:*

Table 5: Beta coefficient, uncorrected p-value, and effect-sizes from the models investigating associations between cortical GM features and E2 timing

| **ROI** | **Thickness** | | | **Sulcal Depth** | | | **Surface area** | | | **Cortical volume** | | |
| --- | --- | --- | --- | --- | --- | --- | --- | --- | --- | --- | --- | --- |
|  | Beta coef | p-value | Cohens’d | Beta coef | p-value | Cohens’d | Beta coef | p-value | Cohens’d | Beta coef | p-value | Cohens’d |
| bankssts | 0.000023 | 0.677 | 0.007 | -0.000133 | 0.401 | -0.014 | -0.12211 | 0.002 | -0.054 | -0.284545 | 0.022 | -0.040 |
| caudal anterior cingulate | 0.00005 | 0.478 | 0.012 | 0.000075 | 0.507 | 0.011 | -0.038586 | 0.14 | -0.025 | -0.049344 | 0.623 | -0.008 |
| caudal middle frontal | -0.00002 | 0.777 | -0.005 | -0.000062 | 0.634 | -0.008 | -0.179498 | 0.113 | -0.027 | -1.027953 | 0.005 | -0.049 |
| cuneus | 0.000012 | 0.838 | 0.004 | 0.000178 | 0.146 | 0.025 | -0.016444 | 0.725 | -0.006 | 0.001674 | 0.991 | 0.000 |
| entorhinal | 0.000157 | 0.339 | 0.016 | 0.000053 | 0.869 | 0.003 | 0.024539 | 0.432 | 0.014 | 0.256427 | 0.093 | 0.029 |
| fusiform | -0.000013 | 0.805 | -0.004 | -0.000041 | 0.654 | -0.008 | -0.144778 | 0.031 | -0.037 | -0.644842 | 0.029 | -0.038 |
| inferior parietal | 0.000007 | 0.91 | 0.002 | 0.000074 | 0.292 | 0.018 | -0.171496 | 0.289 | -0.018 | -0.67893 | 0.183 | -0.023 |
| inferior temporal | -0.000045 | 0.472 | -0.012 | -0.000054 | 0.527 | -0.011 | -0.19383 | 0.034 | -0.037 | -0.795293 | 0.032 | -0.037 |
| isthmus cingulate | -0.000032 | 0.525 | -0.011 | -0.000028 | 0.806 | -0.004 | -0.011308 | 0.715 | -0.006 | -0.095127 | 0.351 | -0.016 |
| lateral occipital | -0.000036 | 0.516 | -0.011 | -0.000064 | 0.375 | -0.015 | -0.1529 | 0.193 | -0.022 | -0.675892 | 0.087 | -0.030 |
| lateral orbitofrontal | -0.000053 | 0.446 | -0.013 | -0.000122 | 0.248 | -0.020 | -0.048558 | 0.694 | -0.007 | -0.261556 | 0.462 | -0.013 |
| lingual | 0.000019 | 0.729 | 0.006 | 0.000056 | 0.513 | 0.011 | -0.037461 | 0.602 | -0.009 | 0.106641 | 0.631 | 0.008 |
| Medial orbitofrontal | -0.000089 | 0.231 | -0.021 | 0.000201 | 0.1 | 0.028 | -0.144594 | 0.115 | -0.027 | -0.60235 | 0.033 | -0.037 |
| middle temporal | 0.000022 | 0.734 | 0.006 | 0.000029 | 0.733 | 0.006 | -0.360266 | 0.0004^#^ | -0.061 | -0.710273 | 0.048 | -0.034 |
| parahippocampal | -0.000078 | 0.317 | -0.017 | 0.000003 | 0.985 | 0.0003 | 0.012425 | 0.63 | 0.008 | -0.048537 | 0.618 | -0.009 |
| paracentral | -0.000044 | 0.513 | -0.011 | 0.000006 | 0.956 | 0.001 | 0.005847 | 0.895 | 0.002 | -0.113617 | 0.537 | -0.011 |
| pars opercularis | -0.000034 | 0.522 | -0.011 | 0.000083 | 0.634 | 0.008 | -0.081449 | 0.187 | -0.023 | -0.377898 | 0.091 | -0.029 |
| pars orbitalis | -0.000009 | 0.903 | -0.002 | -0.000045 | 0.741 | -0.006 | -0.030922 | 0.233 | -0.021 | -0.173874 | 0.129 | -0.026 |
| pars triangularis | -0.000027 | 0.662 | -0.008 | 0.000089 | 0.434 | 0.013 | -0.090318 | 0.12 | -0.027 | -0.333015 | 0.104 | -0.028 |
| pericalcarine | 0.000095 | 0.183 | 0.023 | -0.00005 | 0.722 | -0.006 | -0.078271 | 0.13 | -0.026 | 0.075896 | 0.51 | 0.011 |
| postcentral | -0.00013 | 0.042 | -0.035 | 0.000064 | 0.628 | 0.008 | 0.184233 | 0.184 | 0.023 | -0.241901 | 0.551 | -0.010 |
| posterior cingulate | -0.000047 | 0.304 | -0.018 | 0.000057 | 0.597 | 0.009 | -0.027869 | 0.375 | -0.015 | -0.076365 | 0.478 | -0.012 |
| precentral | 0.000031 | 0.664 | 0.007 | 0.000133 | 0.088 | 0.029 | -0.059357 | 0.722 | -0.006 | -0.219226 | 0.656 | -0.008 |
| precuneus | -0.000029 | 0.566 | -0.010 | 0.000075 | 0.304 | 0.018 | -0.089668 | 0.353 | -0.016 | -0.334743 | 0.372 | -0.015 |
| Rostral anterior cingulate | 0.000102 | 0.31 | 0.017 | -0.000034 | 0.818 | -0.004 | -0.059908 | 0.118 | -0.027 | 0.024154 | 0.871 | 0.003 |
| rostral middle frontal | 0.00003 | 0.629 | 0.008 | -0.000172 | 0.022 | -0.040 | -0.312897 | 0.191 | -0.023 | -1.197487 | 0.106 | -0.028 |
| superior frontal | 0.000025 | 0.687 | 0.007 | -0.00004 | 0.477 | -0.012 | -0.599675 | 0.016 | -0.042 | -2.318867 | 0.007 | -0.046 |
| superior parietal | -0.000011 | 0.87 | -0.003 | -0.000058 | 0.482 | -0.012 | 0.066531 | 0.758 | 0.005 | -0.488873 | 0.487 | -0.012 |
| Superior temporal | -0.000034 | 0.568 | -0.010 | 0.000044 | 0.498 | 0.012 | -0.31522 | 0.001 | -0.055 | -0.86412 | 0.019 | -0.041 |
| supramarginal | 0.00002 | 0.749 | 0.006 | 0.000063 | 0.519 | 0.011 | -0.168558 | 0.32 | -0.017 | -0.048831 | 0.926 | -0.002 |
| frontal pole | 0.000349 | 0.005 | 0.049 | -0.000562 | 0.044 | -0.035 | -0.015537 | 0.378 | -0.015 | 0.069139 | 0.486 | 0.012 |
| temporal pole | 0.000372 | 0.042 | 0.035 | 0.000197 | 0.556 | 0.010 | -0.047135 | 0.073 | -0.031 | -0.038404 | 0.839 | -0.003 |
| transverse temporal | -0.000151 | 0.057 | -0.033 | 0.000109 | 0.532 | 0.011 | -0.010476 | 0.471 | -0.012 | -0.108669 | 0.058 | -0.033 |
| insula | -0.000008 | 0.926 | -0.002 | -0.000451 | 0.058 | -0.033 | 0.133618 | 0.19 | 0.023 | 0.398987 | 0.21 | 0.022 |
| Global | -0.000011 | 0.785 | -0.005 | 0.000004 | 0.535 | 0.011 | -6.577206 | 0.016^#^ | -0.042 | -26.167621 | 0.018^#^ | -0.041 |

##### *E2 Timing and subcortical GM measures across the whole brain:*

Table 6: Beta coefficient, uncorrected p-value, and effect sizes from the models investigating associations between subcortical GM features and E2 timing

| **Regions** | **Beta coefficient** | **p-value** | **Cohen’s d** |
| --- | --- | --- | --- |
| Thalamus proper | - 0.11 | 0.67 | -0.0073 |
| Caudate | -0.138 | 0.132 | -0.025 |
| Putamen | -0.522 | 0.972 | -0.00060 |
| Pallidum | -0.102 | 0.362 | -0.0157 |
| Hippocampus | -0.178 | 0.076 | -0.0305 |
| Amygdala | -0.050 | 0.536 | -0.0106 |
| Accumbens area | -0.005 | 0.905 | 0.00204 |
| Total subcortical volume | -0.3 | 0.77 | -0.0362 |

Table 7: Beta coefficient, uncorrected p-value, and effect-sizes from the models investigating associations between WM microstructures and E2 timing

| **ROI** | **FA** | | | **MD** | | |
| --- | --- | --- | --- | --- | --- | --- |
|  | **Beta coef** | **p-value** | **Cohens’d** | **Beta coef** | **p-value** | **Cohen’s d** |
| Atr | -0.00002071 | 0.147 | -0.0268 | -0.00000008 | 0.996 | -0.0001 |
| Cgc | -0.00002155 | 0.302 | -0.0190 | 0.00000235 | 0.907 | 0.0021 |
| Cgh | -0.00002833 | 0.187 | -0.0243 | -0.00000894 | 0.714 | -0.0067 |
| Cst | 0.00000795 | 0.554 | 0.0109 | -0.00000441 | 0.757 | -0.0057 |
| Fscs | -0.00001073 | 0.393 | -0.0157 | -0.00000070 | 0.951 | -0.0011 |
| Fxcut | 0.00001065 | 0.564 | 0.0106 | -0.00002495 | 0.611 | -0.0094 |
| Fx | -0.00000464 | 0.764 | -0.0055 | -0.00000283 | 0.937 | -0.0015 |
| Ifo | -0.00002933 | 0.030 | -0.0402 | -0.00000085 | 0.951 | -0.0011 |
| Ifsfc | -0.00000215 | 0.858 | -0.0033 | 0.00000081 | 0.950 | 0.0011 |
| Ilf | -0.00002823 | 0.042 | -0.0376 | 0.00000094 | 0.950 | 0.0011 |
| Pscs | 0.00000016 | 0.990 | 0.0002 | 0.00000298 | 0.805 | 0.0046 |
| Pslf | -0.00000323 | 0.782 | -0.0051 | -0.00000535 | 0.674 | -0.0077 |
| Scs | -0.00000606 | 0.607 | -0.0095 | 0.00000129 | 0.911 | 0.0021 |
| Sifc | -0.00003880 | 0.011 | -0.0469 | 0.00000883 | 0.561 | 0.0107 |
| Slf | -0.00000773 | 0.488 | -0.0128 | -0.00000216 | 0.859 | -0.0033 |
| Tslf | -0.00001014 | 0.363 | -0.0168 | 0.00000022 | 0.985 | 0.0003 |
| Unc | 0.00000061 | 0.968 | 0.0007 | -0.00000451 | 0.766 | -0.0055 |
| allfibers | -0.00001427 | 0.137 | -0.0274 | -0.00000051 | 0.966 | -0.0008 |

##### *Split half analysis - E2 Timing and cortical GM measures across the whole brain:*

Table 8: T-value and uncorrected p-value from the models investigating associations between cortical GM features and E2 timing

|  | **Sample - 1** | | | | | | | | **Sample - 2** | | | | | | | |
| --- | --- | --- | --- | --- | --- | --- | --- | --- | --- | --- | --- | --- | --- | --- | --- | --- |
| **ROI** | **Thickness** | | **Sulcal Depth** | | **Surface area** | | **Cortical volume** | | **Thickness** | | **Sulcal Depth** | | **Surface area** | | **Cortical volume** | |
|  | T-value | p-value | T-value | p-value | T-value | p-value | T-value | p-value | T-value | p-value | T-value | p-value | T-value | p-value | T-value | p-value |
| bankssts | 0.166 | 0.868 | -0.948 | 0.344 | -1.470 | 0.143 | -0.750 | 0.454 | 0.378 | 0.706 | -0.078 | 0.938 | -2.961 | 0.003 | -2.662 | 0.008 |
| caudal anterior cingulate | 1.046 | 0.297 | 0.617 | 0.538 | -1.254 | 0.211 | -0.245 | 0.806 | -0.190 | 0.850 | 0.186 | 0.853 | -0.785 | 0.433 | -0.508 | 0.612 |
| caudal middle frontal | -0.773 | 0.440 | -0.098 | 0.922 | -0.792 | 0.429 | -1.719 | 0.087 | 0.291 | 0.771 | -0.707 | 0.480 | -1.447 | 0.150 | -2.256 | 0.025 |
| cuneus | 0.397 | 0.692 | 1.150 | 0.252 | -1.244 | 0.215 | -0.326 | 0.745 | -0.145 | 0.885 | 1.084 | 0.280 | 0.855 | 0.394 | 0.283 | 0.778 |
| entorhinal | 1.829 | 0.069 | 0.082 | 0.935 | 0.713 | 0.477 | 1.999 | 0.047 | -0.255 | 0.799 | 0.188 | 0.851 | 0.202 | 0.840 | 0.502 | 0.617 |
| fusiform | 0.002 | 0.998 | -0.101 | 0.919 | -0.996 | 0.321 | -1.408 | 0.161 | -0.289 | 0.773 | -0.512 | 0.610 | -1.812 | 0.072 | -1.501 | 0.135 |
| inferior parietal | 0.741 | 0.459 | 1.066 | 0.288 | -1.366 | 0.173 | -0.590 | 0.556 | -0.580 | 0.563 | 0.529 | 0.598 | -0.062 | 0.951 | -1.536 | 0.126 |
| inferior temporal | -1.021 | 0.309 | -0.830 | 0.408 | -2.138 | 0.034 | -2.976 | 0.003 | -0.097 | 0.923 | 0.125 | 0.901 | -0.855 | 0.394 | -0.310 | 0.757 |
| isthmus cingulate | -0.245 | 0.807 | 0.291 | 0.771 | 0.023 | 0.982 | -0.893 | 0.373 | -0.653 | 0.515 | -0.892 | 0.374 | -0.478 | 0.633 | -0.406 | 0.686 |
| lateral occipital | -0.559 | 0.577 | -1.049 | 0.295 | -0.444 | 0.657 | -1.240 | 0.216 | -0.200 | 0.841 | -0.222 | 0.824 | -1.329 | 0.186 | -1.095 | 0.275 |
| lateral orbitofrontal | -0.850 | 0.396 | -0.772 | 0.441 | -0.784 | 0.434 | -0.903 | 0.368 | 0.037 | 0.970 | -1.189 | 0.236 | -0.014 | 0.989 | -0.268 | 0.789 |
| lingual | 0.390 | 0.697 | -0.315 | 0.753 | -1.033 | 0.303 | -0.275 | 0.783 | 0.232 | 0.817 | 1.521 | 0.130 | 0.203 | 0.840 | 1.063 | 0.289 |
| Medial orbitofrontal | -1.913 | 0.057 | 0.784 | 0.434 | -1.049 | 0.295 | -2.222 | 0.027 | 0.294 | 0.769 | 1.434 | 0.153 | -1.144 | 0.254 | -0.959 | 0.339 |
| middle temporal | -0.347 | 0.729 | -0.133 | 0.894 | -2.928 | 0.004 | -1.764 | 0.079 | 0.698 | 0.486 | 0.499 | 0.619 | -2.180 | 0.031 | -1.053 | 0.294 |
| parahippocampal | -0.650 | 0.516 | 0.430 | 0.668 | 0.418 | 0.677 | -0.805 | 0.422 | -0.864 | 0.389 | -0.202 | 0.840 | 0.430 | 0.668 | 0.129 | 0.898 |
| paracentral | -0.139 | 0.889 | -0.026 | 0.979 | 0.792 | 0.429 | 0.469 | 0.639 | -0.646 | 0.519 | -0.014 | 0.989 | -0.565 | 0.573 | -1.423 | 0.156 |
| pars opercularis | -0.904 | 0.367 | -0.437 | 0.663 | -0.867 | 0.387 | -1.141 | 0.255 | -0.003 | 0.998 | 1.310 | 0.192 | -1.069 | 0.287 | -1.262 | 0.208 |
| pars orbitalis | -0.357 | 0.721 | -0.393 | 0.694 | -0.955 | 0.341 | -0.831 | 0.407 | 0.049 | 0.961 | -0.339 | 0.735 | -0.466 | 0.642 | -1.137 | 0.257 |
| pars triangularis | -0.848 | 0.397 | 1.800 | 0.073 | -2.099 | 0.037 | -1.870 | 0.063 | 0.151 | 0.880 | -0.377 | 0.707 | -0.055 | 0.956 | -0.522 | 0.602 |
| pericalcarine | 0.299 | 0.765 | -0.688 | 0.493 | -1.630 | 0.105 | -0.551 | 0.582 | 1.476 | 0.142 | 0.297 | 0.767 | -0.332 | 0.740 | 1.449 | 0.149 |
| postcentral | -1.174 | 0.242 | 0.064 | 0.949 | 0.853 | 0.395 | -0.178 | 0.859 | -1.601 | 0.111 | 0.855 | 0.394 | 0.987 | 0.325 | -0.718 | 0.474 |
| posterior cingulate | -1.379 | 0.169 | 1.776 | 0.077 | -0.098 | 0.922 | -0.459 | 0.646 | -0.399 | 0.691 | -1.024 | 0.307 | -1.103 | 0.271 | -0.468 | 0.640 |
| precentral | -0.075 | 0.941 | 0.887 | 0.376 | -0.104 | 0.918 | -0.716 | 0.475 | 0.664 | 0.507 | 1.402 | 0.163 | -0.391 | 0.696 | 0.135 | 0.893 |
| precuneus | 0.150 | 0.881 | 0.281 | 0.779 | -1.217 | 0.225 | -0.652 | 0.515 | -0.929 | 0.354 | 0.908 | 0.365 | -0.126 | 0.900 | -0.627 | 0.531 |
| Rostral anterior cingulate | 0.817 | 0.415 | -0.059 | 0.953 | -0.654 | 0.514 | 0.576 | 0.565 | 0.823 | 0.412 | -0.736 | 0.463 | -1.460 | 0.146 | -0.197 | 0.844 |
| rostral middle frontal | -0.308 | 0.759 | -0.988 | 0.324 | -1.042 | 0.299 | -0.976 | 0.330 | 1.283 | 0.201 | -2.352 | 0.020 | -0.842 | 0.401 | -1.252 | 0.212 |
| superior frontal | -0.348 | 0.729 | -0.966 | 0.335 | -2.133 | 0.034 | -2.097 | 0.037 | 0.867 | 0.387 | -0.403 | 0.687 | -1.235 | 0.218 | -1.646 | 0.101 |
| superior parietal | 0.475 | 0.635 | -1.411 | 0.160 | -0.381 | 0.704 | -0.332 | 0.740 | -0.622 | 0.535 | 0.411 | 0.682 | 0.518 | 0.605 | -0.815 | 0.416 |
| Superior temporal | -0.436 | 0.663 | 1.484 | 0.139 | -2.802 | 0.006 | -2.319 | 0.021 | -0.403 | 0.687 | -0.499 | 0.619 | -1.552 | 0.123 | -1.014 | 0.312 |
| supramarginal | 0.915 | 0.362 | 0.876 | 0.382 | -1.398 | 0.164 | -0.278 | 0.781 | -0.713 | 0.477 | -0.165 | 0.869 | 0.030 | 0.976 | -0.237 | 0.813 |
| frontal pole | 1.693 | 0.092 | -1.491 | 0.137 | -0.138 | 0.890 | 0.699 | 0.485 | 2.282 | 0.024 | -1.588 | 0.114 | -0.983 | 0.327 | 0.338 | 0.736 |
| temporal pole | 1.745 | 0.083 | 0.659 | 0.511 | 0.929 | 0.354 | 1.557 | 0.121 | 1.375 | 0.171 | 0.280 | 0.780 | -3.184 | 0.002 | -1.844 | 0.067 |
| transverse temporal | -2.234 | 0.027 | 0.769 | 0.443 | -1.128 | 0.261 | -2.902 | 0.004 | -0.276 | 0.783 | 0.256 | 0.798 | 0.108 | 0.914 | 0.255 | 0.799 |
| insula | 0.555 | 0.579 | -0.301 | 0.763 | 0.396 | 0.693 | 0.907 | 0.366 | -0.354 | 0.723 | -2.215 | 0.028 | 1.459 | 0.146 | 0.986 | 0.326 |
| Global | -0.160 | 0.873 | 0.137 | 0.891 | -2.222 | 0.027 | -1.746 | 0.082 | -0.053 | 0.958 | 0.741 | 0.460 | -1.381 | 0.169 | -1.473 | 0.143 |

##### *Split half analysis E2 Timing and subcortical GM measures across the whole brain:*

Table 9: T-value and uncorrected p-value from the models investigating associations between sub-cortical GM features and E2 timing

|  | **Sample-1** | | **Sample-2** | |
| --- | --- | --- | --- | --- |
| **Regions** | **T-value** | **p-value** | **T-value** | **p-value** |
| Thalamus proper | -0.645 | 0.520 | 0.145 | 0.885 |
| Caudate | -0.922 | 0.357 | -1.100 | 0.273 |
| Putamen | 0.150 | 0.881 | -0.258 | 0.796 |
| Pallidum | -0.486 | 0.627 | -0.602 | 0.548 |
| Hippocampus | -2.143 | 0.033 | -0.440 | 0.661 |
| Amygdala | -0.452 | 0.652 | -0.453 | 0.651 |
| Accumbens area | 0.096 | 0.924 | 0.271 | 0.787 |
| Total subcortical volume | -2.148 | 0.033 | -0.664 | 0.508 |

##### *Simple half analysis - E2 Timing and WM measures across the whole brain:*

Table 10: T-value and uncorrected p-value from the models investigating associations between WM features and E2 timing

|  | **Sample – 1** | | | | **Sample – 2** | | | |
| --- | --- | --- | --- | --- | --- | --- | --- | --- |
| **ROI** | **FA** | | **MD** | | **FA** | | **MD** | |
|  | **T-value** | **p-value** | **T-value** | **p-value** | **T-value** | **p-value** | **T-value** | **p-value** |
| Atr | -1.962 | 0.052 | 0.536 | 0.593 | 0.029 | 0.977 | -0.439 | 0.661 |
| Cgc | -1.156 | 0.250 | 0.988 | 0.325 | -0.330 | 0.742 | -0.836 | 0.404 |
| Cgh | -0.270 | 0.788 | -0.125 | 0.900 | -1.698 | 0.091 | -0.304 | 0.762 |
| Cst | 0.304 | 0.761 | -0.530 | 0.597 | 0.238 | 0.812 | 0.176 | 0.861 |
| Fscs | -1.684 | 0.094 | 0.355 | 0.723 | 0.324 | 0.746 | -0.445 | 0.657 |
| Fxcut | -0.706 | 0.481 | 0.452 | 0.652 | 1.453 | 0.148 | -1.058 | 0.292 |
| Fx | -0.313 | 0.755 | -0.169 | 0.866 | -0.175 | 0.861 | 0.107 | 0.915 |
| Ifo | -2.070 | 0.040 | 0.123 | 0.902 | -0.938 | 0.350 | -0.248 | 0.804 |
| Ifsfc | -0.919 | 0.360 | 0.783 | 0.435 | 0.621 | 0.536 | -0.686 | 0.494 |
| Ilf | -1.959 | 0.052 | 0.289 | 0.773 | -0.948 | 0.344 | -0.225 | 0.822 |
| Pscs | -0.831 | 0.407 | 0.619 | 0.537 | 0.656 | 0.513 | -0.220 | 0.827 |
| Pslf | -0.855 | 0.394 | 0.105 | 0.916 | 0.487 | 0.627 | -0.723 | 0.471 |
| Scs | -1.493 | 0.138 | 0.591 | 0.556 | 0.607 | 0.545 | -0.411 | 0.681 |
| Sifc | -2.227 | 0.027 | 0.684 | 0.495 | -1.134 | 0.258 | 0.205 | 0.838 |
| Slf | -1.033 | 0.303 | 0.270 | 0.788 | 0.052 | 0.958 | -0.542 | 0.588 |
| Tslf | -1.146 | 0.254 | 0.484 | 0.629 | -0.196 | 0.845 | -0.472 | 0.638 |
| Unc | -0.897 | 0.371 | -0.144 | 0.886 | 1.035 | 0.302 | -0.616 | 0.539 |
| allfibers | -1.814 | 0.072 | 0.099 | 0.921 | -0.302 | 0.763 | -0.170 | 0.865 |

##### *E2 Tempo and cortical GM measures across the whole brain:*

Table 11: Beta coefficient, uncorrected p-value and effect size from investigating associations between corticalGM features and E2 tempo

| **ROI** | **Thickness** | | | **Sulcal Depth** | | | **Surface area** | | | **Cortical volume** | | |
| --- | --- | --- | --- | --- | --- | --- | --- | --- | --- | --- | --- | --- |
|  | Beta coef | p-value | Cohen’s d | Beta coef | p-value | Cohen’s d | Beta coef | p-value | Cohen’s d | Beta coef | p-value | Cohen’s d |
| bankssts | -0.0034 | 0.117 | -0.030 | 0.002 | 0.731 | 0.006 | 1.035 | 0.506 | 0.0128 | -0.833 | 0.865 | -0.003 |
| caudal anterior cingulate | 0.0002 | 0.941 | 0.001 | 0.008 | 0.055 | 0.037 | 0.140 | 0.892 | 0.002 | 1.882 | 0.640 | 0.0090 |
| caudal middle frontal | -0.0056 | 0.039 | -0.040 | 0.004 | 0.419 | 0.015 | 2.070 | 0.649 | 0.008 | -7.493 | 0.611 | -0.009 |
| cuneus | -0.0017 | 0.467 | -0.014 | -0.005 | 0.293 | -0.020 | -2.852 | 0.126 | -0.029 | -9.596 | 0.106 | -0.031 |
| entorhinal | 0.0037 | 0.566 | 0.011 | 0.006 | 0.601 | 0.010 | -1.322 | 0.287 | -0.02 | -3.978 | 0.514 | -0.012 |
| fusiform | -0.0039 | 0.065 | -0.035 | -0.00018 | 0.958 | -0.001 | -3.203 | 0.233 | -0.023 | -25.08 | 0.034 | -0.041 |
| inferior parietal | -0.0066 | 0.004 | -0.055 | -0.006 | 0.026 | -0.043 | -4.303 | 0.503 | -0.012 | -58.555 | 0.004 | -0.055 |
| inferior temporal | -0.0022 | 0.368 | -0.017 | -0.0005 | 0.866 | -0.003 | -3.598 | 0.327 | -0.019 | -24.065 | 0.106 | -0.031 |
| isthmus cingulate | -0.0023 | 0.244 | -0.022 | 0.009 | 0.032 | 0.04 | -2.240 | 0.074 | -0.034 | -9.070 | 0.026 | -0.043 |
| lateral occipital | -0.000015 | 0.994 | -0.0001 | -0.003 | 0.210 | -0.024 | -1.223 | 0.794 | -0.005 | -5.754 | 0.715 | -0.007 |
| lateral orbitofrontal | -0.0012 | 0.649 | -0.0088 | -0.0001 | 0.973 | -0.0006 | 0.905 | 0.853 | 0.003 | -10.339 | 0.468 | -0.014 |
| lingual | -0.0004 | 0.822 | -0.0043 | 0.004 | 0.240 | 0.022 | -7.732 | 0.007 | -0.05 | -16.382 | 0.062 | -0.036 |
| Medial orbitofrontal | 0.0012 | 0.669 | 0.0082 | -0.006 | 0.202 | -0.024 | -1.990 | 0.587 | -0.010 | -4.200 | 0.711 | -0.007 |
| middle temporal | -0.0045 | 0.087 | -0.033 | -0.001 | 0.567 | -0.011 | -1.104 | 0.787 | -0.005 | -25.83 | 0.074 | -0.034 |
| parahippocampal | 0.0044 | 0.154 | 0.027 | 0.002 | 0.675 | 0.008 | -1.013 | 0.321 | -0.019 | 0.5509 | 0.887 | 0.002 |
| paracentral | -0.0030 | 0.262 | -0.021 | 0.001 | 0.788 | 0.005 | -2.133 | 0.222 | -0.023 | -10.172 | 0.160 | -0.027 |
| pars opercularis | -0.0045 | 0.036 | -0.040 | -0.010 | 0.129 | -0.029 | 1.399 | 0.569 | 0.011 | -0.268 | 0.975 | -0.0005 |
| pars orbitalis | -0.0024 | 0.420 | -0.015 | -0.006 | 0.264 | -0.021 | -2.811 | 0.007 | -0.052 | -9.677 | 0.034 | -0.041 |
| pars triangularis | -0.00226 | 0.369 | -0.017 | -0.0004 | 0.916 | -0.002 | -4.798 | 0.043 | -0.039 | -21.45 | 0.009 | -0.050 |
| pericalcarine | -0.00202 | 0.471 | -0.013 | 0.001 | 0.843 | 0.003 | -2.147 | 0.299 | -0.020 | -8.861 | 0.051 | -0.037 |
| postcentral | 0.00197 | 0.436 | 0.0150 | 0.003 | 0.445 | 0.014 | -12.108 | 0.029 | -0.042 | -26.57 | 0.095 | -0.032 |
| posterior cingulate | -0.00278 | 0.130 | -0.029 | -0.002 | 0.560 | -0.011 | -1.0340 | 0.407 | -0.016 | -8.085 | 0.059 | -0.036 |
| precentral | -0.00427 | 0.142 | -0.028 | -0.008 | 0.008 | -0.051 | -2.8644 | 0.672 | -0.0082 | -41.37 | 0.041 | -0.039 |
| precuneus | -0.00325 | 0.104 | -0.031 | -0.001 | 0.570 | -0.011 | -6.4855 | 0.084 | -0.033 | -31.27 | 0.033 | -0.041 |
| Rostral anterior cingulate | -0.00395 | 0.325 | -0.019 | 0.008 | 0.142 | 0.028 | 0.0694 | 0.963 | 0.0008 | -0.859 | 0.885 | -0.002 |
| rostral middle frontal | -0.00523 | 0.038 | -0.040 | 0.004 | 0.127 | 0.029 | -16.537 | 0.084 | -0.033 | -67.71 | 0.021 | -0.044 |
| superior frontal | -0.00417 | 0.097 | -0.032 | -0.0006 | 0.790 | -0.005 | 1.861 | 0.852 | 0.003 | -41.083 | 0.233 | -0.023 |
| superior parietal | -0.00327 | 0.212 | -0.024 | 0.005 | 0.081 | 0.033 | -12.473 | 0.144 | -0.028 | -41.616 | 0.138 | -0.028 |
| Superior temporal | -0.00185 | 0.442 | -0.014 | -0.003 | 0.138 | -0.028 | -2.113 | 0.595 | -0.010 | -18.576 | 0.202 | -0.024 |
| supramarginal | -0.00716 | 0.005 | -0.053 | -0.002 | 0.455 | -0.014 | -4.577 | 0.493 | -0.013 | -53.216 | 0.011 | -0.049 |
| frontal pole | -0.0126 | 0.010 | -0.050 | 0.005 | 0.640 | 0.009 | -0.753 | 0.287 | -0.020 | -7.116 | 0.070 | -0.035 |
| temporal pole | -0.0137 | 0.063 | -0.036 | 0.002 | 0.827 | 0.0042 | 0.602 | 0.567 | 0.011 | -10.165 | 0.182 | -0.025 |
| transverse temporal | 0.00241 | 0.444 | 0.0148 | -0.002 | 0.671 | -0.008 | -0.796 | 0.171 | -0.026 | -0.018 | 0.993 | -0.0001 |
| insula | -0.00059 | 0.871 | -0.003 | 0.008 | 0.381 | 0.0169 | -3.575 | 0.389 | -0.016 | -10.487 | 0.412 | -0.015 |
| Global | -0.002820 | 0.086 | -0.033 | -0.0003 | 0.251 | -0.022 | -242.288 | 0.027# | -0.043 | -1236.272 | 0.004# | -0.055 |

#remained significant after controlling for saliva covariates

##### *E2 Tempo and subcortical GM measures across the whole brain:*

Table 12: Beta coefficient, uncorrected p-value, and effect-sizes from the models investigating associations between sub-cortical GM features and E2 tempo

| **Regions** | **Beta coefficient** | **p-value** | **Cohen’s d** |
| --- | --- | --- | --- |
| Thalamus proper | 0.342 | 0.973 | 0.0006 |
| Caudate | -1.425 | 0.700 | -0.007 |
| Putamen | -11.185 | 0.064 | -0.0359 |
| Pallidum | 5.161 | 0.260 | 0.021 |
| Hippocampus | 0.700 | 0.861 | 0.0033 |
| Amygdala | -0.287 | 0.929 | -0.001 |
| Accumbens area | -1.124 | 0.574 | -0.010 |
| Total subcortical volume | -16.057 | 0.625 | -0.009 |

##### *E2 Tempo and WM measures across the whole brain:*

Table 13: Effect size, beta coefficient and p-value from the models investigating associations WM features and E2 tempo

| **ROI** | **FA** | | | **MD** | | |
| --- | --- | --- | --- | --- | --- | --- |
|  | Beta coef | p-value | Cohens’d | Beta coef | p-value | Cohens’d |
| Atr | 0.00059 | 0.307 | 0.0212 | -0.00005 | 0.937 | -0.0016 |
| Cgc | 0.00024 | 0.776 | 0.0059 | -0.00040 | 0.623 | -0.0102 |
| Cgh | -0.00056 | 0.519 | -0.0134 | 0.00058 | 0.562 | 0.0121 |
| Cst | 0.00086 | 0.115 | 0.0329 | 0.00107 | 0.062 | 0.0389 |
| Fscs | 0.00144 | 0.005 | 0.0589 | 0.00029 | 0.530 | 0.0131 |
| Fxcut | 0.00027 | 0.719 | 0.0075 | 0.00207 | 0.288 | 0.0221 |
| Fx | 0.00006 | 0.917 | 0.0022 | 0.00173 | 0.230 | 0.0250 |
| Ifo | 0.00016 | 0.764 | 0.0063 | 0.00082 | 0.140 | 0.0308 |
| Ifsfc | 0.00080 | 0.102 | 0.0341 | 0.00030 | 0.558 | 0.0122 |
| Ilf | 0.00055 | 0.323 | 0.0206 | 0.00032 | 0.591 | 0.0112 |
| Pscs | 0.00078 | 0.125 | 0.0320 | 0.00035 | 0.474 | 0.0149 |
| Pslf | 0.00014 | 0.760 | 0.0064 | 0.00051 | 0.320 | 0.0207 |
| Scs | 0.00109 | 0.023 | 0.0476 | 0.00037 | 0.422 | 0.0167 |
| Sifc | -0.00013 | 0.840 | -0.0042 | 0.00019 | 0.753 | 0.0065 |
| Slf | 0.00034 | 0.451 | 0.0157 | 0.00046 | 0.350 | 0.0195 |
| Tslf | 0.00033 | 0.461 | 0.0153 | 0.00048 | 0.324 | 0.0205 |
| Unc | 0.00022 | 0.724 | 0.0073 | 0.00075 | 0.214 | 0.0259 |
| allfibers | 0.00035 | 0.366 | 0.0188 | 0.00066 | 0.173 | 0.0284 |

##### *Split half analysis - E2 Tempo and cortical GM measures across the whole brain:*

Table 13: T-value and uncorrected p-value from the models investigating associations between cortical GM features and E2 timing

|  | **Sample - 1** | | | | | | | | **Sample - 2** | | | | | | | |
| --- | --- | --- | --- | --- | --- | --- | --- | --- | --- | --- | --- | --- | --- | --- | --- | --- |
| **ROI** | **Thickness** | | **Sulcal Depth** | | **Surface area** | | **Cortical volume** | | **Thickness** | | **Sulcal Depth** | | **Surface area** | | **Cortical volume** | |
|  | T-value | p-value | T-value | p-value | T-value | p-value | T-value | p-value | T-value | p-value | T-value | p-value | T-value | p-value | T-value | p-value |
| bankssts | -1.185 | 0.238 | -0.664 | 0.508 | 0.246 | 0.806 | -0.145 | 0.885 | -0.965 | 0.336 | 0.977 | 0.330 | 0.647 | 0.519 | -0.101 | 0.920 |
| caudal anterior cingulate | -0.384 | 0.702 | 2.097 | 0.038 | -0.504 | 0.615 | -0.064 | 0.949 | 0.269 | 0.788 | 0.698 | 0.486 | 0.806 | 0.422 | 0.692 | 0.490 |
| caudal middle frontal | -0.223 | 0.824 | 0.341 | 0.733 | 0.872 | 0.385 | 1.595 | 0.113 | -2.984 | 0.003 | 0.851 | 0.396 | -0.168 | 0.867 | -2.470 | 0.015 |
| cuneus | -0.098 | 0.922 | -0.758 | 0.449 | -0.538 | 0.591 | -0.865 | 0.388 | -1.015 | 0.312 | -0.941 | 0.349 | -1.543 | 0.125 | -1.559 | 0.121 |
| entorhinal | -1.099 | 0.273 | -0.137 | 0.891 | -0.959 | 0.339 | -1.370 | 0.173 | 1.865 | 0.064 | 0.867 | 0.387 | -0.385 | 0.701 | 0.220 | 0.826 |
| fusiform | -1.426 | 0.156 | -0.466 | 0.642 | -0.495 | 0.621 | -1.340 | 0.182 | -1.124 | 0.263 | 0.651 | 0.516 | -1.250 | 0.213 | -1.801 | 0.074 |
| inferior parietal | -2.170 | 0.032 | -2.249 | 0.026 | 0.058 | 0.954 | -1.652 | 0.101 | -1.743 | 0.084 | -0.962 | 0.338 | -1.121 | 0.264 | -2.392 | 0.018 |
| inferior temporal | -0.725 | 0.469 | 0.202 | 0.840 | -0.260 | 0.795 | -0.689 | 0.492 | -0.448 | 0.655 | -0.589 | 0.557 | -1.235 | 0.219 | -1.614 | 0.109 |
| isthmus cingulate | -0.547 | 0.585 | 1.492 | 0.138 | -1.623 | 0.107 | -1.180 | 0.240 | -1.162 | 0.247 | 1.505 | 0.135 | -0.921 | 0.359 | -2.066 | 0.041 |
| lateral occipital | 0.714 | 0.477 | -2.016 | 0.046 | 0.592 | 0.555 | 1.431 | 0.154 | -0.433 | 0.666 | 0.174 | 0.862 | -0.984 | 0.327 | -1.833 | 0.069 |
| lateral orbitofrontal | -0.207 | 0.837 | 0.337 | 0.737 | -0.413 | 0.680 | -0.804 | 0.422 | -0.596 | 0.552 | -0.266 | 0.791 | 0.640 | 0.523 | -0.168 | 0.867 |
| lingual | 0.556 | 0.579 | 0.525 | 0.600 | -1.759 | 0.081 | -0.463 | 0.644 | -0.920 | 0.359 | 0.998 | 0.320 | -1.843 | 0.068 | -2.132 | 0.035 |
| Medial orbitofrontal | 0.494 | 0.622 | -0.339 | 0.735 | -0.902 | 0.369 | -0.738 | 0.462 | -0.023 | 0.982 | -1.497 | 0.137 | 0.300 | 0.765 | 0.244 | 0.808 |
| middle temporal | -0.993 | 0.322 | 0.025 | 0.980 | 0.471 | 0.638 | -0.829 | 0.408 | -1.272 | 0.206 | -1.049 | 0.296 | -1.007 | 0.316 | -1.699 | 0.092 |
| parahippocampal | 0.431 | 0.667 | 1.511 | 0.133 | -1.147 | 0.253 | -0.410 | 0.682 | 1.643 | 0.103 | -0.834 | 0.406 | -0.258 | 0.797 | 0.621 | 0.535 |
| paracentral | -0.540 | 0.590 | -0.045 | 0.964 | -0.612 | 0.541 | 0.144 | 0.886 | -1.132 | 0.260 | 0.461 | 0.645 | -0.991 | 0.323 | -2.089 | 0.039 |
| pars opercularis | -1.205 | 0.230 | -0.325 | 0.745 | 1.113 | 0.267 | 0.625 | 0.533 | -1.652 | 0.101 | -1.845 | 0.067 | -0.122 | 0.903 | -0.438 | 0.662 |
| pars orbitalis | 0.384 | 0.701 | -1.142 | 0.255 | -1.989 | 0.048 | 0.010 | 0.992 | -1.628 | 0.106 | -0.333 | 0.740 | -1.863 | 0.065 | -3.026 | 0.003 |
| pars triangularis | -0.099 | 0.921 | -0.509 | 0.612 | -1.296 | 0.197 | -1.327 | 0.186 | -0.973 | 0.332 | 0.358 | 0.721 | -1.422 | 0.157 | -2.110 | 0.037 |
| pericalcarine | 0.864 | 0.389 | 1.234 | 0.219 | -1.136 | 0.258 | -0.658 | 0.512 | -2.089 | 0.039 | -0.804 | 0.423 | -0.246 | 0.806 | -2.193 | 0.030 |
| postcentral | 1.661 | 0.099 | 0.333 | 0.740 | -1.550 | 0.123 | -0.252 | 0.802 | -0.656 | 0.513 | 0.673 | 0.502 | -1.676 | 0.096 | -2.272 | 0.025 |
| posterior cingulate | -1.254 | 0.212 | -0.515 | 0.607 | -0.213 | 0.832 | -1.253 | 0.212 | -0.761 | 0.448 | -0.202 | 0.840 | -0.959 | 0.340 | -1.407 | 0.162 |
| precentral | -1.250 | 0.213 | -3.259 | 0.001 | 0.421 | 0.674 | -0.586 | 0.558 | -0.837 | 0.404 | -0.423 | 0.673 | -1.096 | 0.275 | -2.410 | 0.017 |
| precuneus | -0.941 | 0.348 | 0.646 | 0.519 | -2.318 | 0.022 | -1.747 | 0.083 | -1.418 | 0.159 | -1.416 | 0.159 | -0.289 | 0.773 | -1.384 | 0.169 |
| Rostral anterior cingulate | -0.610 | 0.542 | 1.260 | 0.210 | -0.201 | 0.841 | -0.324 | 0.746 | -1.021 | 0.309 | 0.865 | 0.389 | 0.278 | 0.782 | -0.048 | 0.962 |
| rostral middle frontal | -0.877 | 0.382 | 1.537 | 0.126 | -1.357 | 0.177 | -1.005 | 0.317 | -2.191 | 0.030 | 0.814 | 0.417 | -0.951 | 0.343 | -2.109 | 0.037 |
| superior frontal | -0.476 | 0.635 | 0.092 | 0.927 | 0.578 | 0.564 | 0.757 | 0.450 | -2.045 | 0.043 | -0.217 | 0.828 | -0.269 | 0.789 | -2.335 | 0.021 |
| superior parietal | -0.539 | 0.591 | 1.997 | 0.048 | -1.305 | 0.194 | -0.467 | 0.641 | -1.288 | 0.200 | 0.473 | 0.637 | -0.777 | 0.439 | -1.688 | 0.094 |
| Superior temporal | -0.522 | 0.602 | -0.820 | 0.414 | 0.430 | 0.668 | -0.521 | 0.603 | -0.531 | 0.597 | -1.312 | 0.192 | -1.296 | 0.197 | -1.477 | 0.142 |
| supramarginal | -1.781 | 0.077 | -0.016 | 0.987 | -1.091 | 0.277 | -1.765 | 0.080 | -2.023 | 0.045 | -1.036 | 0.302 | -0.036 | 0.971 | -1.837 | 0.069 |
| frontal pole | -2.302 | 0.023 | 0.525 | 0.600 | -0.697 | 0.487 | -1.443 | 0.151 | -1.431 | 0.155 | 0.113 | 0.910 | -0.832 | 0.407 | -1.184 | 0.239 |
| temporal pole | -1.906 | 0.058 | -0.159 | 0.874 | -0.205 | 0.838 | -1.868 | 0.064 | -0.996 | 0.321 | 0.500 | 0.618 | 1.075 | 0.284 | 0.042 | 0.967 |
| transverse temporal | 0.474 | 0.636 | -1.317 | 0.190 | 0.623 | 0.534 | 0.856 | 0.393 | 0.553 | 0.581 | 0.719 | 0.473 | -2.606 | 0.010 | -1.133 | 0.259 |
| insula | -0.142 | 0.887 | 0.303 | 0.762 | 0.222 | 0.825 | 0.023 | 0.982 | -0.171 | 0.864 | 0.922 | 0.358 | -1.464 | 0.146 | -1.235 | 0.219 |
| Global | -0.819 | 0.414 | -0.496 | 0.621 | -1.283 | 0.201 | -1.110 | 0.269 | -1.660 | 0.099 | -1.071 | 0.286 | -1.927 | 0.056 | -2.749 | 0.007 |

##### *Split half analysis E2 Tempo and subcortical GM measures across the whole brain:*

Table 14: T-value and uncorrected p-value from the models investigating associations sub-cortical GM features and E2 timing

|  | **Sample-1** | | **Sample-2** | |
| --- | --- | --- | --- | --- |
| **Regions** | **T-value** | **p-value** | **T-value** | **p-value** |
| Thalamus proper | -0.310 | 0.757 | 0.431 | 0.667 |
| Caudate | -0.013 | 0.990 | -0.583 | 0.561 |
| Putamen | -0.449 | 0.654 | -2.186 | 0.031 |
| Pallidum | 0.825 | 0.410 | 0.894 | 0.373 |
| Hippocampus | 0.123 | 0.902 | 0.118 | 0.906 |
| Amygdala | 0.890 | 0.375 | -1.021 | 0.309 |
| Accumbens area | -0.348 | 0.728 | -0.521 | 0.603 |
| Total subcortical volume | 0.366 | 0.715 | -1.110 | 0.269 |

##### *Simple half analysis - E2 Tempo and WM measures across the whole brain:*

Table 15: T-value and uncorrected p-value from the models investigating associations between WM features and E2 timing

|  | **Sample – 1** | | | | **Sample – 2** | | | |
| --- | --- | --- | --- | --- | --- | --- | --- | --- |
| **ROI** | **FA** | | **MD** | | **FA** | | **MD** | |
|  | **T-value** | **p-value** | **T-value** | **p-value** | **T-value** | **p-value** | **T-value** | **p-value** |
| Atr | 0.108 | 0.914 | -2.426 | 0.017 | 1.465 | 0.146 | 2.220 | 0.029 |
| Cgc | -0.208 | 0.835 | -0.853 | 0.395 | 0.577 | 0.565 | -0.019 | 0.985 |
| Cgh | 0.182 | 0.856 | -0.442 | 0.659 | -1.035 | 0.303 | 1.055 | 0.294 |
| Cst | 0.785 | 0.434 | -0.538 | 0.591 | 1.569 | 0.120 | 3.149 | 0.002 |
| Fscs | 2.095 | 0.038 | -1.506 | 0.135 | 2.050 | 0.043 | 2.511 | 0.014 |
| Fxcut | 1.254 | 0.212 | 0.929 | 0.355 | -0.745 | 0.458 | 0.418 | 0.677 |
| Fx | 1.087 | 0.279 | 0.743 | 0.459 | -1.178 | 0.242 | 0.927 | 0.356 |
| Ifo | 0.464 | 0.643 | -1.571 | 0.119 | 0.032 | 0.974 | 3.721 | 0.000 |
| Ifsfc | 0.319 | 0.750 | -0.961 | 0.338 | 2.225 | 0.028 | 1.874 | 0.064 |
| Ilf | 1.202 | 0.232 | -0.536 | 0.593 | 0.065 | 0.948 | 1.338 | 0.184 |
| Pscs | 1.639 | 0.104 | -2.136 | 0.035 | 0.638 | 0.525 | 3.280 | 0.001 |
| Pslf | 0.164 | 0.870 | -1.372 | 0.173 | 0.229 | 0.820 | 2.962 | 0.004 |
| Scs | 2.192 | 0.030 | -1.991 | 0.049 | 1.163 | 0.247 | 3.248 | 0.002 |
| Sifc | 0.267 | 0.790 | -1.523 | 0.131 | -0.402 | 0.688 | 1.944 | 0.054 |
| Slf | 0.380 | 0.704 | -1.640 | 0.104 | 0.704 | 0.483 | 3.119 | 0.002 |
| Tslf | 0.398 | 0.691 | -1.870 | 0.064 | 0.734 | 0.464 | 3.365 | 0.001 |
| Unc | 0.376 | 0.707 | -1.075 | 0.285 | 0.151 | 0.880 | 2.829 | 0.006 |
| allfibers | 0.388 | 0.699 | -1.112 | 0.269 | 1.018 | 0.311 | 3.122 | 0.002 |

#### 3. Hemisphere-specific analyses

##### *E2 Timing and GM measures in the left hemisphere:*

Table 16: Beta coefficient, uncorrected p-value, and effect-sizes from the models investigating associations between cortical GM features in the left hemisphere and E2 timing

| **ROI** | **Thickness** | | | **Sulcal Depth** | | | **Surface area** | | | **Cortical volume** | | |
| --- | --- | --- | --- | --- | --- | --- | --- | --- | --- | --- | --- | --- |
|  | Beta coef | p-value | Cohen’s d | Beta coef | p-value | Cohen’s d | Beta coef | p-value | Cohen’s d | Beta coef | p-value | Cohen’s d |
| bankssts | 0.0000909 | 0.189 | 0.023 | -0.0000657 | 0.785 | -0.005 | -0.1362983 | 0.018 | -0.041 | -0.2912469 | 0.110 | -0.028 |
| caudal anterior cingulate | 0.0000591 | 0.543 | 0.010 | 0.0000548 | 0.698 | 0.007 | -0.0247462 | 0.475 | -0.012 | 0.0316297 | 0.804 | 0.004 |
| caudal middle frontal | -0.0000261 | 0.727 | -0.006 | -0.0000375 | 0.803 | -0.004 | -0.0385724 | 0.751 | -0.005 | -0.6868080 | 0.094 | -0.029 |
| cuneus | 0.0000611 | 0.353 | 0.016 | 0.0002571 | 0.118 | 0.027 | -0.0354840 | 0.491 | -0.012 | -0.0126779 | 0.935 | -0.001 |
| entorhinal | -0.0000014 | 0.994 | 0.000 | -0.0002329 | 0.584 | -0.009 | 0.0894715 | 0.048 | 0.034 | 0.4724044 | 0.019 | 0.041 |
| fusiform | -0.0000144 | 0.811 | -0.004 | -0.0000011 | 0.993 | 0.000 | -0.1079863 | 0.213 | -0.021 | -0.6304012 | 0.088 | -0.029 |
| inferior parietal | -0.0000276 | 0.673 | -0.007 | -0.0001500 | 0.094 | -0.029 | -0.3684374 | 0.033 | -0.037 | -1.4623365 | 0.006 | -0.047 |
| inferior temporal | -0.0000248 | 0.732 | -0.006 | -0.0001697 | 0.153 | -0.025 | -0.2413602 | 0.042 | -0.035 | -0.7408280 | 0.110 | -0.028 |
| isthmus cingulate | -0.0001261 | 0.055 | -0.033 | 0.0000924 | 0.533 | 0.011 | -0.0280620 | 0.500 | -0.012 | -0.2649175 | 0.034 | -0.037 |
| lateral occipital | -0.0000092 | 0.876 | -0.003 | -0.0001411 | 0.124 | -0.027 | -0.1147112 | 0.388 | -0.015 | -0.5769549 | 0.191 | -0.023 |
| lateral orbitofrontal | -0.0000885 | 0.265 | -0.019 | -0.0000880 | 0.468 | -0.013 | 0.0841685 | 0.473 | 0.012 | -0.1157790 | 0.748 | -0.006 |
| lingual | 0.0000409 | 0.504 | 0.012 | 0.0001743 | 0.097 | 0.029 | -0.0863737 | 0.311 | -0.017 | -0.0135222 | 0.956 | -0.001 |
| Medial orbitofrontal | -0.0001134 | 0.217 | -0.021 | 0.0002248 | 0.192 | 0.022 | -0.1384709 | 0.266 | -0.019 | -0.4796199 | 0.174 | -0.023 |
| middle temporal | 0.0000494 | 0.523 | 0.011 | 0.0001105 | 0.366 | 0.016 | -0.4225645 | 0.001 | -0.058 | -0.7850830 | 0.070 | -0.031 |
| parahippocampal | -0.0000773 | 0.430 | -0.014 | -0.0001303 | 0.553 | -0.010 | 0.0369950 | 0.311 | 0.017 | -0.0073260 | 0.956 | -0.001 |
| paracentral | -0.0001566 | 0.043 | -0.035 | 0.0000327 | 0.821 | 0.004 | -0.0389028 | 0.454 | -0.013 | -0.4512192 | 0.028 | -0.038 |
| pars opercularis | -0.0000179 | 0.792 | -0.005 | 0.0000281 | 0.861 | 0.003 | -0.0372454 | 0.676 | -0.007 | -0.2750576 | 0.371 | -0.015 |
| pars orbitalis | 0.0000243 | 0.798 | 0.004 | 0.0000727 | 0.697 | 0.007 | -0.0440155 | 0.167 | -0.024 | -0.1977813 | 0.152 | -0.025 |
| pars triangularis | -0.0000954 | 0.204 | -0.022 | 0.0000586 | 0.723 | 0.006 | -0.0126376 | 0.843 | -0.003 | -0.2823137 | 0.203 | -0.022 |
| pericalcarine | 0.0001320 | 0.094 | 0.029 | -0.0000913 | 0.624 | -0.008 | -0.1228659 | 0.037 | -0.036 | 0.0515190 | 0.685 | 0.007 |
| postcentral | -0.0000952 | 0.175 | -0.023 | 0.0002135 | 0.124 | 0.027 | 0.1014800 | 0.528 | 0.011 | -0.3437913 | 0.463 | -0.013 |
| posterior cingulate | -0.0000518 | 0.385 | -0.015 | 0.0001813 | 0.151 | 0.025 | -0.0392679 | 0.329 | -0.017 | -0.1009605 | 0.456 | -0.013 |
| precentral | -0.0000089 | 0.907 | -0.002 | -0.0000728 | 0.444 | -0.013 | 0.1218496 | 0.489 | 0.012 | 0.0735100 | 0.889 | 0.002 |
| precuneus | -0.0000609 | 0.266 | -0.019 | 0.0000741 | 0.398 | 0.015 | -0.0605002 | 0.591 | -0.009 | -0.3423775 | 0.408 | -0.014 |
| Rostral anterior cingulate | 0.0001617 | 0.203 | 0.022 | -0.0000083 | 0.966 | -0.001 | -0.0783568 | 0.174 | -0.023 | 0.0624433 | 0.757 | 0.005 |
| rostral middle frontal | 0.0000280 | 0.687 | 0.007 | -0.0001948 | 0.028 | -0.038 | -0.1990367 | 0.459 | -0.013 | -0.8323104 | 0.334 | -0.017 |
| superior frontal | 0.0000072 | 0.912 | 0.002 | -0.0000062 | 0.927 | -0.002 | -0.2964747 | 0.264 | -0.019 | -1.6407176 | 0.085 | -0.030 |
| superior parietal | -0.0000259 | 0.705 | -0.007 | 0.0000005 | 0.996 | 0.000 | -0.2947940 | 0.222 | -0.021 | -1.4197227 | 0.069 | -0.031 |
| Superior temporal | 0.0000181 | 0.795 | 0.004 | 0.0000804 | 0.362 | 0.016 | -0.3272234 | 0.010 | -0.044 | -0.8611464 | 0.057 | -0.033 |
| supramarginal | -0.0000046 | 0.952 | -0.001 | 0.0000970 | 0.383 | 0.015 | -0.5034392 | 0.017 | -0.041 | -1.0827981 | 0.096 | -0.029 |
| frontal pole | 0.0003014 | 0.054 | 0.033 | -0.0006502 | 0.067 | -0.032 | -0.0192183 | 0.351 | -0.016 | 0.0100567 | 0.927 | 0.002 |
| temporal pole | 0.0002359 | 0.267 | 0.019 | 0.0000305 | 0.948 | 0.001 | -0.0140737 | 0.690 | -0.007 | -0.0048773 | 0.983 | 0.000 |
| transverse temporal | -0.0002060 | 0.032 | -0.037 | 0.0002624 | 0.271 | 0.019 | -0.0261792 | 0.239 | -0.020 | -0.1841347 | 0.022 | -0.040 |
| insula | 0.0000557 | 0.606 | 0.009 | -0.0003880 | 0.272 | -0.019 | 0.0452978 | 0.747 | 0.006 | 0.3535354 | 0.352 | 0.016 |

Table 17: Beta coefficient, uncorrected p-value, and effect-sizes from the models investigating associations between sub-cortical GM features in the left hemisphere and E2 timing

| **Regions** | **Beta coefficient** | **p-value** | **Cohen’s d** |
| --- | --- | --- | --- |
| Thalamus proper | 0.0651659 | 0.850 | 0.003 |
| Caudate | -0.2034492 | 0.063 | -0.032 |
| Putamen | -0.0550173 | 0.777 | -0.005 |
| Pallidum | -0.0931354 | 0.541 | -0.011 |
| Hippocampus | -0.2418072 | 0.072 | -0.031 |
| Amygdala | -0.1173742 | 0.293 | -0.018 |
| Accumbens area | 0.0192445 | 0.772 | 0.005 |

##### *E2 Timing and GM measures in the right hemisphere:*

Table 18: Beta coefficient, uncorrected p-value, and effect-sizes from the models investigating associations between cortical GM features in the right hemisphere and E2 timing

| **ROI** | **Thickness** | | | **Sulcal Depth** | | | **Surface area** | | | **Cortical volume** | | |
| --- | --- | --- | --- | --- | --- | --- | --- | --- | --- | --- | --- | --- |
|  | Beta coef | p-value | Cohen’s d | Beta coef | p-value | Cohen’s d | Beta coef | p-value | Cohen’s d | Beta coef | p-value | Cohen’s d |
| bankssts | -0.0000454 | 0.504 | -0.012 | -0.0001686 | 0.407 | -0.014 | -0.1126854 | 0.012 | -0.044 | -0.3252704 | 0.021 | -0.040 |
| caudal anterior cingulate | 0.0000413 | 0.621 | 0.009 | 0.0000817 | 0.577 | 0.010 | -0.0394816 | 0.314 | -0.017 | -0.1217048 | 0.363 | -0.016 |
| caudal middle frontal | -0.0000139 | 0.864 | -0.003 | -0.0000778 | 0.697 | -0.007 | -0.2992348 | 0.061 | -0.032 | -1.3929140 | 0.006 | -0.048 |
| cuneus | -0.0000374 | 0.572 | -0.010 | 0.0001232 | 0.436 | 0.013 | -0.0025735 | 0.969 | -0.001 | -0.0151631 | 0.941 | -0.001 |
| entorhinal | 0.0003199 | 0.129 | 0.026 | 0.0003199 | 0.452 | 0.013 | -0.0384112 | 0.318 | -0.017 | 0.0696874 | 0.722 | 0.006 |
| fusiform | -0.0000170 | 0.782 | -0.005 | -0.0000848 | 0.486 | -0.012 | -0.1668751 | 0.051 | -0.034 | -0.6275977 | 0.084 | -0.030 |
| inferior parietal | 0.0000398 | 0.539 | 0.011 | 0.0002972 | 0.004 | 0.050 | -0.0664584 | 0.756 | -0.005 | -0.0780827 | 0.906 | -0.002 |
| inferior temporal | -0.0000575 | 0.415 | -0.014 | 0.0000587 | 0.613 | 0.009 | -0.1424871 | 0.186 | -0.023 | -0.8659224 | 0.037 | -0.036 |
| isthmus cingulate | 0.0000560 | 0.396 | 0.015 | -0.0001682 | 0.298 | -0.018 | 0.0155454 | 0.752 | 0.005 | 0.0988816 | 0.499 | 0.012 |
| lateral occipital | -0.0000618 | 0.299 | -0.018 | 0.0000182 | 0.840 | 0.003 | -0.2012419 | 0.156 | -0.024 | -0.8662785 | 0.058 | -0.033 |
| lateral orbitofrontal | -0.0000036 | 0.968 | -0.001 | -0.0001613 | 0.264 | -0.019 | -0.1317051 | 0.455 | -0.013 | -0.3855156 | 0.398 | -0.015 |
| lingual | -0.0000017 | 0.978 | 0.000 | -0.0000420 | 0.702 | -0.007 | 0.0038792 | 0.965 | 0.001 | 0.1533629 | 0.558 | 0.010 |
| Medial orbitofrontal | -0.0000595 | 0.502 | -0.012 | 0.0001543 | 0.280 | 0.019 | -0.1336005 | 0.216 | -0.021 | -0.6971117 | 0.044 | -0.035 |
| middle temporal | 0.0000065 | 0.929 | 0.002 | -0.0000255 | 0.812 | -0.004 | -0.3083804 | 0.010 | -0.045 | -0.8177783 | 0.055 | -0.033 |
| parahippocampal | -0.0000800 | 0.391 | -0.015 | 0.0001005 | 0.620 | 0.009 | -0.0134571 | 0.683 | -0.007 | -0.0897444 | 0.468 | -0.013 |
| paracentral | 0.0000567 | 0.449 | 0.013 | -0.0000192 | 0.903 | -0.002 | 0.0415906 | 0.485 | 0.012 | 0.1634864 | 0.487 | 0.012 |
| pars opercularis | -0.0000414 | 0.529 | -0.011 | 0.0001025 | 0.728 | 0.006 | -0.1182874 | 0.104 | -0.028 | -0.5402013 | 0.049 | -0.034 |
| pars orbitalis | -0.0000402 | 0.661 | -0.008 | -0.0001470 | 0.394 | -0.015 | -0.0118001 | 0.745 | -0.006 | -0.1591872 | 0.295 | -0.018 |
| pars triangularis | 0.0000499 | 0.524 | 0.011 | 0.0000998 | 0.518 | 0.011 | -0.1512570 | 0.087 | -0.030 | -0.3149975 | 0.307 | -0.018 |
| pericalcarine | 0.0000525 | 0.522 | 0.011 | -0.0000034 | 0.985 | 0.000 | -0.0371563 | 0.592 | -0.009 | 0.0943823 | 0.517 | 0.011 |
| postcentral | -0.0001487 | 0.061 | -0.032 | -0.0000635 | 0.742 | -0.006 | 0.2388892 | 0.174 | 0.023 | -0.2364173 | 0.614 | -0.009 |
| posterior cingulate | -0.0000507 | 0.382 | -0.015 | -0.0000640 | 0.652 | -0.008 | -0.0172646 | 0.689 | -0.007 | -0.0910867 | 0.508 | -0.011 |
| precentral | 0.0000693 | 0.438 | 0.013 | 0.0003139 | 0.004 | 0.050 | -0.2523262 | 0.265 | -0.019 | -0.7166015 | 0.281 | -0.019 |
| precuneus | 0.0000038 | 0.947 | 0.001 | 0.0000338 | 0.752 | 0.005 | -0.0885451 | 0.500 | -0.012 | -0.3586490 | 0.444 | -0.013 |
| Rostral anterior cingulate | 0.0000651 | 0.585 | 0.009 | -0.0000214 | 0.917 | -0.002 | -0.0269746 | 0.514 | -0.011 | -0.0187734 | 0.903 | -0.002 |
| rostral middle frontal | 0.0000406 | 0.570 | 0.010 | -0.0001731 | 0.097 | -0.029 | -0.3885844 | 0.223 | -0.021 | -1.5197945 | 0.125 | -0.026 |
| superior frontal | 0.0000406 | 0.563 | 0.010 | -0.0000661 | 0.369 | -0.015 | -0.8511444 | 0.011 | -0.044 | -3.0637004 | 0.006 | -0.048 |
| superior parietal | 0.0000004 | 0.996 | 0.000 | -0.0000823 | 0.496 | -0.012 | 0.2887870 | 0.318 | 0.017 | 0.2136057 | 0.816 | 0.004 |
| Superior temporal | -0.0000838 | 0.216 | -0.021 | 0.0000020 | 0.981 | 0.000 | -0.3185744 | 0.004 | -0.050 | -0.9835698 | 0.015 | -0.042 |
| supramarginal | 0.0000442 | 0.556 | 0.010 | 0.0000019 | 0.990 | 0.000 | 0.0380301 | 0.863 | 0.003 | 0.6080892 | 0.376 | 0.015 |
| frontal pole | 0.0003896 | 0.011 | 0.044 | -0.0005603 | 0.127 | -0.026 | -0.0147347 | 0.580 | -0.010 | 0.1038087 | 0.466 | 0.013 |
| temporal pole | 0.0005000 | 0.022 | 0.040 | 0.0002698 | 0.515 | 0.011 | -0.0811501 | 0.017 | -0.041 | -0.0775948 | 0.748 | -0.006 |
| transverse temporal | -0.0000961 | 0.331 | -0.017 | -0.0000195 | 0.933 | -0.001 | -0.0000447 | 0.998 | 0.000 | -0.0540972 | 0.382 | -0.015 |
| insula | -0.0000716 | 0.514 | -0.011 | -0.0004952 | 0.090 | -0.029 | 0.2330405 | 0.092 | 0.029 | 0.4874912 | 0.214 | 0.021 |

Table 19: Beta coefficient, uncorrected p-value, and effect-sizes from the models investigating associations between sub-cortical GM features in the right hemisphere and E2 timing

| **Regions** | **Beta coefficient** | **p-value** | **Cohen’s d** |
| --- | --- | --- | --- |
| Thalamus proper | -0.2610822 | 0.345 | -0.016 |
| Caudate | -0.0747875 | 0.498 | -0.012 |
| Putamen | 0.0186210 | 0.915 | 0.002 |
| Pallidum | -0.1059791 | 0.410 | -0.014 |
| Hippocampus | -0.1042353 | 0.379 | -0.015 |
| Amygdala | 0.0363763 | 0.700 | 0.007 |
| Accumbens area | -0.0085418 | 0.869 | -0.003 |

##### *E2 timing and WM measures in the left hemisphere*

Table 20: Beta coefficient, uncorrected p-value, and effect-sizes from the models investigating associations between WM features in the left hemisphere and E2 timing

| **ROI** | **FA** | | | **MD** | | |
| --- | --- | --- | --- | --- | --- | --- |
|  | **Beta coef** | **p-value** | **Cohens’d** | **Beta coef** | **p-value** | **Cohen’s d** |
| Atr | 0.0000009 | 0.960 | 0.001 | 0.0000039 | 0.826 | 0.004 |
| Cgc | -0.0000013 | 0.961 | -0.001 | 0.0000632 | 0.024 | 0.042 |
| Cgh | -0.0000511 | 0.048 | -0.037 | -0.0000086 | 0.757 | -0.006 |
| Cst | -0.0000011 | 0.946 | -0.001 | -0.0000085 | 0.588 | -0.010 |
| Fscs | -0.0000120 | 0.389 | -0.016 | -0.0000111 | 0.403 | -0.015 |
| Fxcut | -0.0000027 | 0.896 | -0.002 | -0.0000434 | 0.425 | -0.015 |
| Fx | -0.0000089 | 0.599 | -0.010 | -0.0000191 | 0.612 | -0.009 |
| Ifo | -0.0000324 | 0.039 | -0.038 | -0.0000159 | 0.355 | -0.017 |
| Ifsfc | -0.0000050 | 0.730 | -0.006 | -0.0000152 | 0.339 | -0.018 |
| Ilf | -0.0000209 | 0.210 | -0.023 | -0.0000122 | 0.523 | -0.012 |
| Pscs | -0.0000169 | 0.265 | -0.021 | -0.0000058 | 0.662 | -0.008 |
| Pslf | -0.0000111 | 0.432 | -0.014 | -0.0000105 | 0.482 | -0.013 |
| Scs | -0.0000162 | 0.211 | -0.023 | -0.0000091 | 0.477 | -0.013 |
| Sifc | -0.0000300 | 0.078 | -0.033 | -0.0000129 | 0.508 | -0.012 |
| Slf | -0.0000158 | 0.245 | -0.021 | -0.0000051 | 0.720 | -0.007 |
| Tslf | -0.0000163 | 0.241 | -0.022 | -0.0000038 | 0.787 | -0.005 |
| Unc | -0.0000092 | 0.627 | -0.009 | -0.0000176 | 0.373 | -0.016 |

##### *E2 timing and WM measure in the right hemisphere*

Table 21: Beta coefficient, uncorrected p-value, and effect-sizes from the models investigating associations between WM features in the right hemisphere and E2 timing

| **ROI** | **FA** | | | **MD** | | |
| --- | --- | --- | --- | --- | --- | --- |
|  | **Beta coef** | **p-value** | **Cohens’d** | **Beta coef** | **p-value** | **Cohen’s d** |
| Atr | -0.0000382 | 0.026 | -0.041 | -0.0000081 | 0.658 | -0.008 |
| Cgc | -0.0000368 | 0.157 | -0.026 | -0.0000593 | 0.025 | -0.042 |
| Cgh | -0.0000050 | 0.840 | -0.004 | -0.0000050 | 0.860 | -0.003 |
| Cst | 0.0000166 | 0.260 | 0.021 | -0.0000010 | 0.946 | -0.001 |
| Fscs | -0.0000066 | 0.651 | -0.008 | 0.0000087 | 0.478 | 0.013 |
| Fxcut | 0.0000225 | 0.308 | 0.019 | 0.0000010 | 0.987 | 0.0003 |
| Fx | 0.0000001 | 0.996 | 0.000 | 0.0000178 | 0.651 | 0.008 |
| Ifo | -0.0000242 | 0.100 | -0.030 | 0.0000136 | 0.406 | 0.015 |
| Ifsfc | 0.0000030 | 0.813 | 0.004 | 0.0000123 | 0.372 | 0.016 |
| Ilf | -0.0000317 | 0.048 | -0.037 | 0.0000175 | 0.357 | 0.017 |
| Pscs | 0.0000182 | 0.266 | 0.021 | 0.0000104 | 0.415 | 0.015 |
| Pslf | 0.0000077 | 0.542 | 0.011 | -0.0000027 | 0.841 | -0.004 |
| Scs | 0.0000063 | 0.676 | 0.008 | 0.0000113 | 0.366 | 0.017 |
| Sifc | -0.0000422 | 0.025 | -0.042 | 0.0000258 | 0.169 | 0.025 |
| Slf | 0.0000033 | 0.786 | 0.005 | -0.0000014 | 0.918 | -0.002 |
| Tslf | -0.0000003 | 0.980 | 0.0005 | 0.0000025 | 0.846 | 0.004 |
| Unc | 0.0000130 | 0.448 | 0.014 | 0.0000081 | 0.636 | 0.009 |

##### *E2 tempo and GM measures in the left hemisphere*

Table 22: Beta coefficient, uncorrected p-value, and effect-sizes from the models investigating associations between cortical GM features in the left hemi and E2 tempo

| **ROI** | **Thickness** | | | **Sulcal Depth** | | | **Surface area** | | | **Cortical volume** | | |
| --- | --- | --- | --- | --- | --- | --- | --- | --- | --- | --- | --- | --- |
|  | Beta coef | p-value | Cohen’s d | Beta coef | p-value | Cohen’s d | Beta coef | p-value | Cohen’s d | Beta coef | p-value | Cohen’s d |
| bankssts | -0.00664 | 0.015 | -0.048 | 0.00051 | 0.956 | 0.001 | -0.59050 | 0.796 | -0.005 | -6.53527 | 0.363 | -0.018 |
| caudal anterior cingulate | -0.00020 | 0.959 | -0.001 | 0.01493 | 0.009 | 0.051 | -0.01190 | 0.993 | 0.000 | 0.08157 | 0.987 | 0.000 |
| caudal middle frontal | -0.00504 | 0.099 | -0.032 | 0.00241 | 0.683 | 0.008 | -3.89546 | 0.423 | -0.016 | -22.60812 | 0.172 | -0.027 |
| cuneus | -0.00152 | 0.565 | -0.011 | -0.00714 | 0.280 | -0.021 | -0.25026 | 0.902 | -0.002 | -1.68380 | 0.786 | -0.005 |
| entorhinal | 0.01089 | 0.151 | 0.028 | -0.00293 | 0.864 | -0.003 | -1.78018 | 0.324 | -0.019 | -1.06672 | 0.894 | -0.003 |
| fusiform | -0.00548 | 0.024 | -0.044 | 0.00169 | 0.726 | 0.007 | -3.23544 | 0.356 | -0.018 | -27.58560 | 0.064 | -0.036 |
| inferior parietal | -0.00700 | 0.008 | -0.052 | -0.00425 | 0.236 | -0.023 | 3.05017 | 0.657 | 0.009 | -29.79268 | 0.160 | -0.027 |
| inferior temporal | -0.00373 | 0.202 | -0.025 | -0.00423 | 0.376 | -0.017 | -6.80917 | 0.155 | -0.028 | -38.99160 | 0.037 | -0.041 |
| isthmus cingulate | -0.00172 | 0.517 | -0.013 | 0.00884 | 0.129 | 0.029 | -2.61524 | 0.110 | -0.031 | -12.70734 | 0.010 | -0.051 |
| lateral occipital | -0.00009 | 0.970 | -0.001 | -0.00412 | 0.262 | -0.022 | -7.52611 | 0.154 | -0.028 | -19.76410 | 0.262 | -0.022 |
| lateral orbitofrontal | 0.00058 | 0.854 | 0.004 | 0.00634 | 0.186 | 0.026 | -5.94748 | 0.200 | -0.025 | -20.86144 | 0.149 | -0.028 |
| lingual | -0.00247 | 0.306 | -0.020 | 0.00101 | 0.808 | 0.005 | -6.23936 | 0.069 | -0.035 | -15.31082 | 0.112 | -0.031 |
| Medial orbitofrontal | 0.00545 | 0.141 | 0.029 | -0.00288 | 0.681 | -0.008 | -1.24707 | 0.804 | -0.005 | 7.65596 | 0.591 | 0.010 |
| middle temporal | -0.00659 | 0.037 | -0.041 | -0.00634 | 0.196 | -0.025 | -3.12463 | 0.542 | -0.012 | -42.95753 | 0.014 | -0.048 |
| parahippocampal | 0.00497 | 0.211 | 0.024 | 0.01522 | 0.080 | 0.034 | -3.72539 | 0.011 | -0.050 | -6.92399 | 0.198 | -0.025 |
| paracentral | -0.00058 | 0.852 | -0.004 | -0.00001 | 0.998 | 0.000 | -0.74296 | 0.716 | -0.007 | 0.00514 | 0.999 | 0.000 |
| pars opercularis | -0.00387 | 0.160 | -0.027 | -0.00829 | 0.200 | -0.025 | 1.44844 | 0.682 | 0.008 | 4.35519 | 0.720 | 0.007 |
| pars orbitalis | -0.00315 | 0.406 | -0.016 | -0.00837 | 0.262 | -0.022 | -0.77799 | 0.544 | -0.012 | -5.21660 | 0.341 | -0.018 |
| pars triangularis | 0.00077 | 0.799 | 0.005 | 0.00093 | 0.889 | 0.003 | -4.14991 | 0.097 | -0.032 | -13.97152 | 0.105 | -0.032 |
| pericalcarine | -0.00294 | 0.347 | -0.018 | -0.00647 | 0.390 | -0.017 | -0.45407 | 0.848 | -0.004 | -5.30067 | 0.296 | -0.020 |
| postcentral | 0.00093 | 0.737 | 0.006 | 0.00114 | 0.832 | 0.004 | -9.76035 | 0.129 | -0.030 | -25.13895 | 0.175 | -0.026 |
| posterior cingulate | -0.00307 | 0.200 | -0.025 | -0.00315 | 0.535 | -0.012 | -0.86191 | 0.585 | -0.011 | -8.59103 | 0.111 | -0.031 |
| precentral | -0.00521 | 0.088 | -0.033 | -0.00448 | 0.237 | -0.023 | -7.64271 | 0.277 | -0.021 | -60.64232 | 0.004 | -0.056 |
| precuneus | -0.00353 | 0.108 | -0.031 | -0.00225 | 0.513 | -0.013 | -7.32359 | 0.078 | -0.034 | -40.54541 | 0.010 | -0.050 |
| Rostral anterior cingulate | -0.00340 | 0.507 | -0.013 | 0.01518 | 0.046 | 0.039 | 1.23281 | 0.594 | 0.010 | 4.97975 | 0.541 | 0.012 |
| rostral middle frontal | -0.00361 | 0.199 | -0.025 | 0.00525 | 0.139 | 0.029 | -18.58925 | 0.076 | -0.035 | -64.42871 | 0.052 | -0.038 |
| superior frontal | -0.00392 | 0.136 | -0.029 | -0.00047 | 0.861 | -0.003 | -4.02226 | 0.702 | -0.007 | -41.10310 | 0.272 | -0.021 |
| superior parietal | -0.00377 | 0.173 | -0.026 | 0.00527 | 0.159 | 0.027 | -8.54103 | 0.361 | -0.018 | -36.08370 | 0.241 | -0.023 |
| Superior temporal | -0.00430 | 0.130 | -0.029 | -0.00245 | 0.491 | -0.013 | -2.38558 | 0.643 | -0.009 | -28.98750 | 0.108 | -0.031 |
| supramarginal | -0.00901 | 0.004 | -0.057 | 0.00004 | 0.992 | 0.000 | -1.16703 | 0.887 | -0.003 | -49.08118 | 0.057 | -0.037 |
| frontal pole | -0.01189 | 0.057 | -0.037 | 0.00936 | 0.505 | 0.013 | -0.59179 | 0.479 | -0.014 | -4.81266 | 0.274 | -0.021 |
| temporal pole | -0.01078 | 0.213 | -0.024 | -0.00505 | 0.788 | -0.005 | 0.41157 | 0.773 | 0.006 | -4.32826 | 0.646 | -0.009 |
| transverse temporal | 0.00348 | 0.368 | 0.017 | 0.00390 | 0.685 | 0.008 | -0.16864 | 0.850 | -0.004 | 2.06181 | 0.521 | 0.012 |
| insula | -0.00320 | 0.458 | -0.014 | 0.00021 | 0.988 | 0.000 | 1.03363 | 0.855 | 0.004 | -2.16904 | 0.888 | -0.003 |

Table 23: Beta coefficient, uncorrected p-value, and effect-sizes from the models investigating associations between sub-cortical GM features in the left hemi and E2 tempo

| **Regions** | **Beta coefficient** | **p-value** | **Cohen’s d** |
| --- | --- | --- | --- |
| Thalamus proper | -9.23120 | 0.500 | -0.013 |
| Caudate | 1.74091 | 0.690 | 0.008 |
| Putamen | -13.10517 | 0.097 | -0.032 |
| Pallidum | 5.50166 | 0.373 | 0.017 |
| Hippocampus | 3.66353 | 0.495 | 0.013 |
| Amygdala | -2.58112 | 0.566 | -0.011 |
| Accumbens area | -0.99208 | 0.709 | -0.007 |

##### *E2 tempo and GM measures in the right hemisphere*

Table 24: Beta coefficient, uncorrected p-value, and effect-sizes from the models investigating associations between cortical GM features in the right hemi and E2 tempo

| **ROI** | **Thickness** | | | **Sulcal Depth** | | | **Surface area** | | | **Cortical volume** | | |
| --- | --- | --- | --- | --- | --- | --- | --- | --- | --- | --- | --- | --- |
|  | Beta coef | p-value | Cohen’s d | Beta coef | p-value | Cohen’s d | Beta coef | p-value | Cohen’s d | Beta coef | p-value | Cohen’s d |
| bankssts | -0.00033 | 0.904 | -0.002 | 0.00300 | 0.712 | 0.007 | 2.85317 | 0.108 | 0.031 | 5.48870 | 0.328 | 0.019 |
| caudal anterior cingulate | 0.00007 | 0.984 | 0.000 | 0.00307 | 0.601 | 0.010 | 0.54435 | 0.729 | 0.007 | 3.32750 | 0.536 | 0.012 |
| caudal middle frontal | -0.00628 | 0.051 | -0.038 | 0.00573 | 0.487 | 0.013 | 6.80176 | 0.291 | 0.020 | 7.00572 | 0.731 | 0.007 |
| cuneus | -0.00221 | 0.403 | -0.016 | -0.00514 | 0.421 | -0.016 | -5.36890 | 0.046 | -0.039 | -17.09055 | 0.039 | -0.040 |
| entorhinal | -0.00380 | 0.652 | -0.009 | 0.01493 | 0.377 | 0.017 | -0.87091 | 0.570 | -0.011 | -9.50740 | 0.229 | -0.023 |
| fusiform | -0.00198 | 0.426 | -0.015 | -0.00175 | 0.718 | -0.007 | -3.40540 | 0.318 | -0.019 | -21.97052 | 0.134 | -0.029 |
| inferior parietal | -0.00605 | 0.021 | -0.045 | -0.00833 | 0.047 | -0.039 | -10.23329 | 0.233 | -0.023 | -89.12339 | 0.001 | -0.064 |
| inferior temporal | -0.00109 | 0.705 | -0.007 | 0.00382 | 0.408 | 0.016 | -0.11463 | 0.979 | -0.001 | -12.54681 | 0.455 | -0.014 |
| isthmus cingulate | -0.00274 | 0.304 | -0.020 | 0.01106 | 0.088 | 0.033 | -1.70707 | 0.400 | -0.016 | -6.68449 | 0.260 | -0.022 |
| lateral occipital | 0.00012 | 0.959 | 0.001 | -0.00326 | 0.371 | -0.017 | 4.56171 | 0.426 | 0.015 | 10.15606 | 0.579 | 0.011 |
| lateral orbitofrontal | -0.00322 | 0.365 | -0.018 | -0.00683 | 0.235 | -0.023 | 5.11871 | 0.467 | 0.014 | -0.10410 | 0.995 | 0.000 |
| lingual | 0.00129 | 0.598 | 0.010 | 0.00581 | 0.189 | 0.026 | -8.80462 | 0.014 | -0.048 | -17.38165 | 0.094 | -0.033 |
| Medial orbitofrontal | -0.00292 | 0.413 | -0.016 | -0.00993 | 0.084 | -0.034 | -2.65476 | 0.540 | -0.012 | -17.91995 | 0.198 | -0.025 |
| middle temporal | -0.00306 | 0.305 | -0.020 | 0.00083 | 0.847 | 0.004 | 1.27965 | 0.789 | 0.005 | -9.65781 | 0.572 | -0.011 |
| parahippocampal | 0.00430 | 0.247 | 0.022 | -0.00854 | 0.302 | -0.020 | 1.67491 | 0.205 | 0.025 | 8.07925 | 0.107 | 0.031 |
| paracentral | -0.00506 | 0.091 | -0.033 | 0.00253 | 0.684 | 0.008 | -4.06283 | 0.086 | -0.033 | -21.07497 | 0.024 | -0.044 |
| pars opercularis | -0.00537 | 0.043 | -0.039 | -0.01052 | 0.385 | -0.017 | 0.00281 | 0.999 | 0.000 | -5.33127 | 0.634 | -0.009 |
| pars orbitalis | -0.00153 | 0.680 | -0.008 | -0.00551 | 0.424 | -0.015 | -4.79695 | 0.001 | -0.064 | -14.25920 | 0.021 | -0.045 |
| pars triangularis | -0.00559 | 0.078 | -0.034 | -0.00145 | 0.818 | -0.004 | -5.25755 | 0.153 | -0.028 | -29.87530 | 0.019 | -0.046 |
| pericalcarine | -0.00110 | 0.735 | -0.007 | 0.00932 | 0.199 | 0.025 | -3.73959 | 0.184 | -0.026 | -13.23845 | 0.022 | -0.045 |
| postcentral | 0.00221 | 0.484 | 0.014 | 0.00478 | 0.518 | 0.013 | -13.95941 | 0.046 | -0.039 | -28.02985 | 0.126 | -0.030 |
| posterior cingulate | -0.00247 | 0.286 | -0.021 | -0.00153 | 0.787 | -0.005 | -1.50998 | 0.385 | -0.017 | -7.05867 | 0.197 | -0.025 |
| precentral | -0.00308 | 0.394 | -0.017 | -0.01169 | 0.008 | -0.052 | 0.14015 | 0.988 | 0.000 | -24.74844 | 0.372 | -0.017 |
| precuneus | -0.00315 | 0.162 | -0.027 | -0.00029 | 0.946 | -0.001 | -5.04213 | 0.339 | -0.019 | -24.10537 | 0.194 | -0.025 |
| Rostral anterior cingulate | -0.00576 | 0.229 | -0.023 | 0.00189 | 0.819 | 0.004 | -1.42447 | 0.389 | -0.017 | -6.91688 | 0.261 | -0.022 |
| rostral middle frontal | -0.00713 | 0.014 | -0.048 | 0.00533 | 0.211 | 0.024 | -14.90389 | 0.256 | -0.022 | -68.76733 | 0.091 | -0.033 |
| superior frontal | -0.00449 | 0.113 | -0.031 | -0.00009 | 0.975 | -0.001 | 2.59682 | 0.848 | 0.004 | -55.39443 | 0.216 | -0.024 |
| superior parietal | -0.00274 | 0.331 | -0.019 | 0.00500 | 0.304 | 0.020 | -13.88195 | 0.227 | -0.023 | -47.84770 | 0.189 | -0.025 |
| Superior temporal | 0.00025 | 0.926 | 0.002 | -0.00505 | 0.146 | -0.028 | -1.63676 | 0.709 | -0.007 | -11.07203 | 0.489 | -0.013 |
| supramarginal | -0.00523 | 0.085 | -0.033 | -0.00426 | 0.486 | -0.014 | -5.08006 | 0.561 | -0.011 | -50.49905 | 0.068 | -0.036 |
| frontal pole | -0.01322 | 0.031 | -0.042 | 0.00193 | 0.896 | 0.003 | -0.90960 | 0.391 | -0.017 | -9.22067 | 0.102 | -0.032 |
| temporal pole | -0.01691 | 0.054 | -0.037 | 0.01145 | 0.491 | 0.013 | 0.57776 | 0.672 | 0.008 | -15.08354 | 0.124 | -0.030 |
| transverse temporal | 0.00130 | 0.743 | 0.006 | -0.00940 | 0.306 | -0.020 | -1.51746 | 0.018 | -0.046 | -2.41041 | 0.333 | -0.019 |
| insula | 0.00184 | 0.675 | 0.008 | 0.01559 | 0.183 | 0.026 | -8.64589 | 0.121 | -0.030 | -19.99578 | 0.206 | -0.025 |

Table 25: Beta coefficient, uncorrected p-value, and effect-sizes from the models investigating associations between sub-cortical GM features in the right hemi and E2 tempo

| **Regions** | **Beta coefficient** | **p-value** | **Cohen’s d** |
| --- | --- | --- | --- |
| Thalamus proper | 11.09405 | 0.321 | 0.019 |
| Caudate | -4.31959 | 0.334 | -0.019 |
| Putamen | -9.18782 | 0.200 | -0.025 |
| Pallidum | 6.23509 | 0.237 | 0.023 |
| Hippocampus | -2.67042 | 0.572 | -0.011 |
| Amygdala | 1.60796 | 0.672 | 0.008 |
| Accumbens area | -0.83227 | 0.691 | -0.008 |

##### *E2 tempo and WM measures in the left hemisphere*

Table 26: Beta coefficient, uncorrected p-value, and effect-sizes from the models investigating associations between WM features in the left hemisphere and E2 tempo

| **ROI** | **FA** | | | **MD** | | |
| --- | --- | --- | --- | --- | --- | --- |
|  | **Beta coef** | **p-value** | **Cohens’d** | **Beta coef** | **p-value** | **Cohen’s d** |
| Atr | 0.00048 | 0.486 | 0.015 | -0.00039 | 0.590 | -0.011 |
| Cgc | 0.00048 | 0.661 | 0.009 | -0.00120 | 0.292 | -0.022 |
| Cgh | -0.00012 | 0.908 | -0.002 | 0.00118 | 0.300 | 0.022 |
| Cst | 0.00111 | 0.079 | 0.037 | 0.00110 | 0.086 | 0.036 |
| Fscs | 0.00112 | 0.047 | 0.041 | 0.00019 | 0.722 | 0.007 |
| Fxcut | 0.00007 | 0.937 | 0.002 | 0.00049 | 0.818 | 0.005 |
| Fx | 0.00015 | 0.827 | 0.005 | 0.00084 | 0.578 | 0.012 |
| Ifo | 0.00033 | 0.599 | 0.011 | 0.00034 | 0.624 | 0.010 |
| Ifsfc | 0.00084 | 0.150 | 0.030 | 0.00030 | 0.641 | 0.010 |
| Ilf | 0.00029 | 0.666 | 0.009 | -0.00039 | 0.616 | -0.010 |
| Pscs | 0.00038 | 0.542 | 0.013 | 0.00021 | 0.701 | 0.008 |
| Pslf | 0.00009 | 0.873 | 0.003 | 0.00030 | 0.625 | 0.010 |
| Scs | 0.00073 | 0.170 | 0.029 | 0.00026 | 0.607 | 0.011 |
| Sifc | 0.00037 | 0.596 | 0.011 | 0.00021 | 0.793 | 0.005 |
| Slf | 0.00052 | 0.347 | 0.020 | 0.00019 | 0.744 | 0.007 |
| Tslf | 0.00075 | 0.186 | 0.028 | 0.00018 | 0.760 | 0.006 |
| Unc | 0.00069 | 0.366 | 0.019 | 0.00038 | 0.635 | 0.010 |

##### *E2 tempo and WM measures in the right hemisphere*

Table 27: Beta coefficient, uncorrected p-value, and effect-sizes from the models investigating associations between WM features in the right hemisphere and E2 tempo

| **ROI** | **FA** | | | **MD** | | |
| --- | --- | --- | --- | --- | --- | --- |
|  | **Beta coef** | **p-value** | **Cohens’d** | **Beta coef** | **p-value** | **Cohen’s d** |
| Atr | 0.00065 | 0.346 | 0.020 | 0.00041 | 0.579 | 0.012 |
| Cgc | -0.00009 | 0.929 | -0.002 | 0.00039 | 0.714 | 0.008 |
| Cgh | -0.00099 | 0.320 | -0.021 | -0.00023 | 0.841 | -0.004 |
| Cst | 0.00066 | 0.266 | 0.023 | 0.00104 | 0.082 | 0.036 |
| Fscs | 0.00171 | 0.004 | 0.060 | 0.00036 | 0.461 | 0.015 |
| Fxcut | 0.00056 | 0.523 | 0.013 | 0.00344 | 0.138 | 0.031 |
| Fx | 0.00008 | 0.907 | 0.002 | 0.00255 | 0.110 | 0.033 |
| Ifo | 0.00006 | 0.925 | 0.002 | 0.00110 | 0.093 | 0.035 |
| Ifsfc | 0.00060 | 0.248 | 0.024 | 0.00042 | 0.444 | 0.016 |
| Ilf | 0.00078 | 0.231 | 0.025 | 0.00092 | 0.227 | 0.025 |
| Pscs | 0.00112 | 0.091 | 0.035 | 0.00052 | 0.314 | 0.021 |
| Pslf | 0.00011 | 0.826 | 0.005 | 0.00075 | 0.171 | 0.029 |
| Scs | 0.00137 | 0.026 | 0.046 | 0.00045 | 0.366 | 0.019 |
| Sifc | -0.00063 | 0.412 | -0.017 | 0.00013 | 0.861 | 0.004 |
| Slf | 0.00011 | 0.829 | 0.005 | 0.00073 | 0.166 | 0.029 |
| Tslf | -0.00015 | 0.759 | -0.006 | 0.00079 | 0.125 | 0.032 |
| Unc | -0.00022 | 0.754 | -0.007 | 0.00110 | 0.111 | 0.033 |

#### **4. Sensitivity Analyses:**

##### Models with covariates

Table 28: Beta coefficient, uncorrected p-value, and effect-sizes from the models investigating associations between cortical GM features and E2 timing – showing results only for the significant features from the main model

| **ROI** | Model with race/ethnicity | | | Model with parent education | | | Model with parent combine income | | | Model with BMIz | | |
| --- | --- | --- | --- | --- | --- | --- | --- | --- | --- | --- | --- | --- |
|  | Beta coef | p-value | Cohen’s d | Beta coef | p-value | Cohen’s d | Beta coef | p-value | Cohen’s d | Beta coef | p-value | Cohen’s d |
| Surface area_ bankssts | -0.117 | 0.0028 | -0.051 | -0.122 | 0.0019 | -0.053 | -0.121 | 0.0019 | -.053 | -0.106 | 0.007 | -0.046 |
| Surface area_ middle temporal | -0.348 | 0.00067 | -0.059 | -0.360 | 0.0004 | -0.061 | -0.359 | 0.00046 | -0.0608 | -0.286 | 0.005 | -0.048 |
| Surface area_ Superior temporal | -0.309 | 0.0018 | -0.054 | -0.315 | 0.0014 | -0.055 | -0.3146 | 0.0015 | -0.054 | -0.240 | 0.0148 | -0.042 |
| Total surface area | -6.226 | 0.022 | -0.039 | -6.579 | 0.015 | -0.041 | -6.560 | 0.0161 | -0.0416 | -4.279 | 0.113 | -0.027 |
| Total cortical volume | -23.662 | 0.0316 | -0.037 | -26.185 | 0.0178 | -0.0409 | -26.113 | 0.0181 | -0.0408 | -16.922 | 0.122 | -0.026 |

Table 29: Beta coefficient, uncorrected p-value, and effect-sizes from the models investigating associations between cortical GM features and E2 tempo - showing results only for the significant features from the main model

| **ROI** | Model with race/ethnicity | | | Model with parent education | | | Model with parent combine income | | | Model with BMIz | | |
| --- | --- | --- | --- | --- | --- | --- | --- | --- | --- | --- | --- | --- |
|  | Beta coef | p-value | Cohen’s d | Beta coef | p-value | Cohen’s d | Beta coef | p-value | Cohen’s d | Beta coef | p-value | Cohen’s d |
| Total surface area | -235.242 | 0.031 | -0.0418 | -242.23 | 0.027 | -o.043 | -239.609 | 0.028 | -0.042 | -186.870 | 0.085 | -0.034 |
| Total cortical volume | -1213.771 | 0.0054 | -0.054 | -1234.923 | 0.004 | -0.055 | -1228.734 | 0.005 | -0.054 | -1059.520 | 0.014 | -0.047 |

##### Models with winsorised data

Table 30: Beta coefficient, uncorrected p-value, and effect-sizes from the models investigating associations between cortical GM features and E2 timing across the whole brain

| **ROI** | **Thickness** | | | **Sulcal Depth** | | | **Surface area** | | | **Cortical volume** | | |
| --- | --- | --- | --- | --- | --- | --- | --- | --- | --- | --- | --- | --- |
|  | Beta coef | p-value | Cohen’s d | Beta coef | p-value | Cohen’s d | Beta coef | p-value | Cohen’s d | Beta coef | p-value | Cohen’s d |
| bankssts | 0.000024 | 0.66 | 0.0076 | -0.000115 | 0.464 | -0.0126 | -0.116147 | 0.003 | -0.0516 | -0.266012 | 0.031 | -0.0373 |
| caudal anterior cingulate | 0.000046 | 0.518 | 0.0112 | 0.000063 | 0.567 | 0.0099 | -0.03977 | 0.122 | -0.0267 | -0.070119 | 0.482 | -0.0121 |
| caudal middle frontal | -0.000016 | 0.813 | -0.0041 | -0.000057 | 0.659 | -0.0076 | -0.195123 | 0.08 | -0.0302 | -1.013702 | 0.005 | -0.0481 |
| cuneus | 0.000031 | 0.598 | 0.0091 | 0.000157 | 0.198 | 0.0222 | -0.014892 | 0.748 | -0.0055 | 0.028311 | 0.848 | 0.0033 |
| entorhinal | 0.000144 | 0.379 | 0.0152 | -0.00003 | 0.925 | -0.0016 | 0.017607 | 0.565 | 0.0099 | 0.26556 | 0.076 | 0.0307 |
| fusiform | -0.000016 | 0.752 | -0.0055 | -0.000036 | 0.693 | -0.0068 | -0.143257 | 0.032 | -0.037 | -0.624769 | 0.034 | -0.0366 |
| inferior parietal | 0.000002 | 0.972 | 0.0006 | 0.000075 | 0.281 | 0.0186 | -0.147883 | 0.356 | -0.0159 | -0.651698 | 0.2 | -0.0221 |
| inferior temporal | -0.000056 | 0.36 | -0.0158 | -0.000042 | 0.621 | -0.0085 | -0.198197 | 0.03 | -0.0374 | -0.785564 | 0.034 | -0.0366 |
| isthmus cingulate | -0.00003 | 0.548 | -0.0104 | -0.000034 | 0.77 | -0.005 | -0.004973 | 0.866 | -0.0029 | -0.067103 | 0.506 | -0.0115 |
| lateral occipital | -0.000029 | 0.594 | -0.0092 | -0.000072 | 0.318 | -0.0172 | -0.146957 | 0.21 | -0.0216 | -0.62361 | 0.113 | -0.0273 |
| lateral orbitofrontal | -0.000053 | 0.444 | -0.0132 | -0.000092 | 0.382 | -0.0151 | -0.050078 | 0.684 | -0.007 | -0.283321 | 0.423 | -0.0138 |
| lingual | 0.000035 | 0.517 | 0.0112 | 0.000084 | 0.324 | 0.017 | -0.041275 | 0.565 | -0.0099 | 0.183005 | 0.41 | 0.0142 |
| Medial orbitofrontal | -0.000095 | 0.195 | -0.0223 | 0.000192 | 0.115 | 0.0272 | -0.140332 | 0.125 | -0.0265 | -0.612435 | 0.029 | -0.0377 |
| middle temporal | 0.000021 | 0.735 | 0.0058 | 0.000021 | 0.801 | 0.0043 | -0.354542 | 0 | -0.0614 | -0.667008 | 0.061 | -0.0323 |
| parahippocampal | -0.000062 | 0.424 | -0.0138 | -0.000008 | 0.957 | -0.0009 | 0.003583 | 0.889 | 0.0024 | -0.058973 | 0.541 | -0.0105 |
| paracentral | -0.00004 | 0.552 | -0.0103 | 0.00002 | 0.858 | 0.0031 | 0.013332 | 0.761 | 0.0052 | -0.0772 | 0.674 | -0.0072 |
| pars opercularis | -0.000038 | 0.472 | -0.0124 | 0.000096 | 0.576 | 0.0096 | -0.05601 | 0.355 | -0.016 | -0.302868 | 0.168 | -0.0238 |
| pars orbitalis | -0.000012 | 0.872 | -0.0028 | -0.000029 | 0.833 | -0.0036 | -0.031761 | 0.219 | -0.0212 | -0.192544 | 0.091 | -0.0291 |
| pars triangularis | -0.000024 | 0.693 | -0.0068 | 0.00006 | 0.595 | 0.0091 | -0.085353 | 0.141 | -0.0254 | -0.299495 | 0.142 | -0.0253 |
| pericalcarine | 0.000116 | 0.102 | 0.0282 | -0.000057 | 0.671 | -0.0073 | -0.083929 | 0.101 | -0.0283 | 0.108665 | 0.342 | 0.0164 |
| postcentral | -0.000119 | 0.059 | -0.0326 | 0.000097 | 0.295 | 0.0181 | 0.207774 | 0.124 | 0.0266 | -0.152226 | 0.705 | -0.0065 |
| posterior cingulate | -0.000049 | 0.279 | -0.0187 | 0.000054 | 0.614 | 0.0087 | -0.018127 | 0.563 | -0.01 | -0.048547 | 0.65 | -0.0078 |
| precentral | 0.000021 | 0.763 | 0.0052 | 0.000121 | 0.089 | 0.0293 | -0.060817 | 0.711 | -0.0064 | -0.129784 | 0.789 | -0.0046 |
| precuneus | -0.000031 | 0.526 | -0.0109 | 0.000065 | 0.367 | 0.0155 | -0.095465 | 0.322 | -0.0171 | -0.311146 | 0.405 | -0.0144 |
| Rostral anterior cingulate | 0.00013 | 0.194 | 0.0224 | -0.000033 | 0.824 | -0.0038 | -0.061477 | 0.108 | -0.0277 | 0.041306 | 0.781 | 0.0048 |
| rostral middle frontal | 0.00003 | 0.622 | 0.0085 | -0.000168 | 0.024 | -0.0391 | -0.280068 | 0.241 | -0.0202 | -1.10896 | 0.134 | -0.0259 |
| superior frontal | 0.000022 | 0.727 | 0.006 | -0.000037 | 0.515 | -0.0112 | -0.571381 | 0.021 | -0.0401 | -2.230438 | 0.01 | -0.0447 |
| superior parietal | -0.000002 | 0.97 | -0.0006 | -0.000063 | 0.442 | -0.0133 | 0.076736 | 0.721 | 0.0062 | -0.420841 | 0.548 | -0.0104 |
| Superior temporal | -0.00003 | 0.613 | -0.0087 | 0.000035 | 0.589 | 0.0093 | -0.291569 | 0.002 | -0.0524 | -0.828595 | 0.024 | -0.0391 |
| supramarginal | 0.000017 | 0.781 | 0.0048 | 0.000018 | 0.846 | 0.0034 | -0.163117 | 0.332 | -0.0167 | 0.079264 | 0.88 | 0.0026 |
| frontal pole | 0.000343 | 0.005 | 0.0488 | -0.000584 | 0.034 | -0.0365 | -0.015042 | 0.391 | -0.0148 | 0.068421 | 0.486 | 0.012 |
| temporal pole | 0.00033 | 0.062 | 0.0323 | 0.000127 | 0.701 | 0.0066 | -0.047085 | 0.071 | -0.0312 | -0.059298 | 0.749 | -0.0055 |
| transverse temporal | -0.000145 | 0.066 | -0.0317 | 0.000099 | 0.567 | 0.0099 | -0.009556 | 0.506 | -0.0115 | -0.106517 | 0.062 | -0.0323 |
| insula | -0.000012 | 0.894 | -0.0023 | -0.000525 | 0.027 | -0.0381 | 0.163081 | 0.109 | 0.0277 | 0.454269 | 0.153 | 0.0247 |
| Global | -0.000009 | 0.816 | -0.004 | 0.000002 | 0.724 | 0.0061 | -6.205886 | 0.022 | -0.0396 | -24.164048 | 0.029 | -0.0378 |

Table 31: Beta coefficient, uncorrected p-value, and effect-sizes from the models investigating associations between sub-cortical GM features and E2 timing

| **Regions** | **Beta coefficient** | **p-value** | **Cohen’s d** |
| --- | --- | --- | --- |
| Thalamus proper | -0.152067 | 0.556 | -0.0101 |
| Caudate | -0.11949 | 0.193 | -0.0224 |
| Putamen | 0.063043 | 0.67 | 0.0073 |
| Pallidum | -0.126196 | 0.247 | -0.02 |
| Hippocampus | -0.180477 | 0.072 | -0.0311 |
| Amygdala | -0.05346 | 0.51 | -0.0114 |
| Accumbens area | 0.001746 | 0.972 | 0.0006 |
| Total subcortical volume | -1.719058 | 0.039 | -0.0357 |

Table 32: Beta coefficient, uncorrected p-value, and effect-sizes from the models investigating associations between WM features and E2 timing across the whole brain

| **ROI** | **FA** | | | **MD** | | |
| --- | --- | --- | --- | --- | --- | --- |
|  | **Beta coef** | **p-value** | **Cohens’d** | **Beta coef** | **p-value** | **Cohen’s d** |
| Atr | -0.0000227 | 0.108 | -0.0296 | -0.0000016 | 0.924 | -0.0018 |
| Cgc | -0.0000259 | 0.212 | -0.023 | -0.0000027 | 0.891 | -0.0025 |
| Cgh | -0.0000274 | 0.197 | -0.0238 | -0.0000092 | 0.695 | -0.0072 |
| Cst | 0.0000090 | 0.497 | 0.0125 | -0.0000028 | 0.839 | -0.0037 |
| Fscs | -0.0000138 | 0.269 | -0.0204 | -0.0000007 | 0.952 | -0.0011 |
| Fxcut | 0.0000068 | 0.71 | 0.0069 | -0.0000146 | 0.756 | -0.0057 |
| Fx | -0.0000043 | 0.781 | -0.0051 | -0.0000063 | 0.855 | -0.0034 |
| Ifo | -0.0000282 | 0.034 | -0.0393 | -0.0000006 | 0.964 | -0.0008 |
| Ifsfc | -0.0000039 | 0.741 | -0.0061 | -0.0000001 | 0.995 | -0.0001 |
| Ilf | -0.0000308 | 0.026 | -0.0413 | 0.0000024 | 0.872 | 0.003 |
| Pscs | -0.0000013 | 0.916 | -0.0019 | 0.0000038 | 0.748 | 0.0059 |
| Pslf | -0.0000047 | 0.682 | -0.0075 | -0.0000049 | 0.696 | -0.0072 |
| Scs | -0.0000085 | 0.467 | -0.0134 | 0.0000021 | 0.851 | 0.0035 |
| Sifc | -0.0000369 | 0.016 | -0.0447 | 0.0000107 | 0.472 | 0.0133 |
| Slf | -0.0000079 | 0.478 | -0.0131 | -0.0000020 | 0.868 | -0.0031 |
| Tslf | -0.0000101 | 0.36 | -0.0169 | 0.0000007 | 0.953 | 0.0011 |
| Unc | -0.0000010 | 0.945 | -0.0013 | -0.0000036 | 0.808 | -0.0045 |
| allfibers | -0.0000148 | 0.119 | -0.0288 | -0.0000010 | 0.931 | -0.0016 |

Table 33: Beta coefficient, uncorrected p-value, and effect-sizes from the models investigating associations between cortical GM features and E2 tempo

| **ROI** | **Thickness** | | | **Sulcal Depth** | | | **Surface area** | | | **Cortical volume** | | |
| --- | --- | --- | --- | --- | --- | --- | --- | --- | --- | --- | --- | --- |
|  | Beta coef | p-value | Cohen’s d | Beta coef | p-value | Cohen’s d | Beta coef | p-value | Cohen’s d | Beta coef | p-value | Cohen’s d |
| bankssts | -0.00373 | 0.107 | -0.0313 | 0.001504 | 0.816 | 0.0045 | 1.033018 | 0.525 | 0.0123 | -1.06317 | 0.835 | -0.004 |
| caudal anterior cingulate | 0.000321 | 0.915 | 0.0021 | 0.009757 | 0.036 | 0.0407 | 0.323443 | 0.763 | 0.0058 | 2.761052 | 0.512 | 0.0127 |
| caudal middle frontal | -0.00587 | 0.038 | -0.0404 | 0.003947 | 0.475 | 0.0139 | 2.593323 | 0.583 | 0.0107 | -5.33869 | 0.73 | -0.0067 |
| cuneus | -0.00271 | 0.272 | -0.0213 | -0.00517 | 0.316 | -0.0195 | -2.84678 | 0.144 | -0.0283 | -11.2022 | 0.069 | -0.0354 |
| entorhinal | 0.005013 | 0.467 | 0.0141 | 0.005346 | 0.691 | 0.0077 | -1.10896 | 0.388 | -0.0168 | -3.14544 | 0.616 | -0.0097 |
| fusiform | -0.00372 | 0.092 | -0.0327 | 0.000405 | 0.914 | 0.0021 | -3.61811 | 0.199 | -0.0249 | -26.283 | 0.035 | -0.0411 |
| inferior parietal | -0.00663 | 0.006 | -0.0541 | -0.00673 | 0.024 | -0.044 | -4.95481 | 0.461 | -0.0143 | -59.0965 | 0.006 | -0.0533 |
| inferior temporal | -0.00175 | 0.5 | -0.0131 | -0.00125 | 0.728 | -0.0067 | -2.87439 | 0.457 | -0.0144 | -21.9452 | 0.161 | -0.0272 |
| isthmus cingulate | -0.00185 | 0.379 | -0.0171 | 0.010194 | 0.034 | 0.0412 | -2.55248 | 0.041 | -0.0398 | -9.49258 | 0.026 | -0.0434 |
| lateral occipital | -0.00039 | 0.868 | -0.0032 | -0.00296 | 0.333 | -0.0188 | -0.46249 | 0.925 | -0.0018 | -7.76355 | 0.639 | -0.0091 |
| lateral orbitofrontal | -0.00109 | 0.706 | -0.0073 | -0.00046 | 0.916 | -0.0021 | 0.24093 | 0.963 | 0.0009 | -10.2535 | 0.492 | -0.0133 |
| lingual | -0.00131 | 0.567 | -0.0111 | 0.003637 | 0.31 | 0.0197 | -7.73223 | 0.011 | -0.0495 | -18.7345 | 0.042 | -0.0395 |
| Medial orbitofrontal | 0.001523 | 0.625 | 0.0095 | -0.00565 | 0.271 | -0.0213 | -1.64635 | 0.669 | -0.0083 | -1.87794 | 0.874 | -0.0031 |
| middle temporal | -0.0052 | 0.051 | -0.0379 | -0.0013 | 0.715 | -0.0071 | -0.34668 | 0.934 | -0.0016 | -27.8598 | 0.066 | -0.0358 |
| parahippocampal | 0.004533 | 0.168 | 0.0268 | 0.002525 | 0.694 | 0.0076 | -0.79878 | 0.454 | -0.0145 | 1.417818 | 0.728 | 0.0067 |
| paracentral | -0.00322 | 0.256 | -0.022 | -5E-06 | 0.999 | 0 | -2.61349 | 0.153 | -0.0277 | -11.8532 | 0.119 | -0.0303 |
| pars opercularis | -0.00429 | 0.057 | -0.037 | -0.01225 | 0.098 | -0.0321 | 0.680651 | 0.79 | 0.0052 | -2.61403 | 0.777 | -0.0055 |
| pars orbitalis | -0.00233 | 0.463 | -0.0142 | -0.00585 | 0.31 | -0.0197 | -2.72809 | 0.013 | -0.0485 | -9.00386 | 0.06 | -0.0365 |
| pars triangularis | -0.00272 | 0.298 | -0.0202 | 0.001289 | 0.79 | 0.0052 | -5.1115 | 0.041 | -0.0398 | -22.9201 | 0.008 | -0.0517 |
| pericalcarine | -0.00328 | 0.27 | -0.0214 | 0.003032 | 0.595 | 0.0103 | -2.17305 | 0.313 | -0.0196 | -11.2857 | 0.017 | -0.0464 |
| postcentral | 0.002025 | 0.442 | 0.0149 | 0.00133 | 0.732 | 0.0066 | -12.9557 | 0.023 | -0.0443 | -28.4413 | 0.087 | -0.0333 |
| posterior cingulate | -0.00287 | 0.133 | -0.0291 | -0.0018 | 0.691 | -0.0077 | -1.44366 | 0.27 | -0.0214 | -8.6775 | 0.053 | -0.0376 |
| precentral | -0.00368 | 0.215 | -0.0241 | -0.00817 | 0.006 | -0.0535 | -2.96008 | 0.673 | -0.0082 | -40.1495 | 0.056 | -0.0372 |
| precuneus | -0.00344 | 0.097 | -0.0323 | -0.00116 | 0.702 | -0.0074 | -6.35116 | 0.107 | -0.0313 | -32.6035 | 0.034 | -0.0413 |
| Rostral anterior cingulate | -0.00436 | 0.304 | -0.02 | 0.009172 | 0.136 | 0.0289 | 0.200944 | 0.901 | 0.0024 | -0.75343 | 0.905 | -0.0023 |
| rostral middle frontal | -0.00575 | 0.028 | -0.0427 | 0.004465 | 0.158 | 0.0274 | -19.0734 | 0.058 | -0.0368 | -74.6812 | 0.016 | -0.0471 |
| superior frontal | -0.00411 | 0.116 | -0.0305 | -0.0012 | 0.613 | -0.0098 | -1.2014 | 0.908 | -0.0022 | -42.4254 | 0.242 | -0.0227 |
| superior parietal | -0.00362 | 0.187 | -0.0256 | 0.00571 | 0.098 | 0.0322 | -13.3177 | 0.136 | -0.0289 | -44.6094 | 0.13 | -0.0294 |
| Superior temporal | -0.00198 | 0.435 | -0.0151 | -0.00368 | 0.182 | -0.0259 | -2.21139 | 0.586 | -0.0105 | -20.9408 | 0.171 | -0.0266 |
| supramarginal | -0.00693 | 0.009 | -0.0511 | -0.00146 | 0.708 | -0.0073 | -4.27378 | 0.541 | -0.0119 | -55.1052 | 0.013 | -0.0485 |
| frontal pole | -0.01355 | 0.008 | -0.0518 | 0.007 | 0.546 | 0.0117 | -0.68125 | 0.358 | -0.0178 | -6.90155 | 0.092 | -0.0327 |
| temporal pole | -0.0123 | 0.101 | -0.0319 | 0.006422 | 0.648 | 0.0089 | 0.593447 | 0.587 | 0.0105 | -9.95129 | 0.202 | -0.0248 |
| transverse temporal | 0.003135 | 0.343 | 0.0184 | -0.00296 | 0.683 | -0.0079 | -0.7487 | 0.215 | -0.0241 | 0.628678 | 0.792 | 0.0051 |
| insula | -0.00047 | 0.902 | -0.0024 | 0.009054 | 0.372 | 0.0173 | -4.56704 | 0.294 | -0.0203 | -9.28339 | 0.493 | -0.0133 |
| Global | -0.00293 | 0.087 | -0.0332 | -0.00025 | 0.392 | -0.0166 | -252.272 | 0.028 | -0.0428 | -1285.56 | 0.005 | -0.0545 |

Table 34: Beta coefficient, uncorrected p-value, and effect-sizes from the models investigating associations between sub-cortical GM features and E2 tempo

| **Regions** | **Beta coefficient** | **p-value** | **Cohen’s d** |
| --- | --- | --- | --- |
| Thalamus proper | 0.66441 | 0.951 | 0.0012 |
| Caudate | -1.47215 | 0.704 | -0.0074 |
| Putamen | -13.2608 | 0.037 | -0.0406 |
| Pallidum | 6.274313 | 0.181 | 0.026 |
| Hippocampus | 1.442447 | 0.731 | 0.0067 |
| Amygdala | -0.34043 | 0.92 | -0.0019 |
| Accumbens area | -0.97408 | 0.64 | -0.0091 |
| Total subcortical volume | -16.4284 | 0.634 | -0.0092 |

Table 35: Beta coefficient, uncorrected p-value, and effect-sizes from the models investigating associations between WM features and E2 tempo

| **ROI** | **FA** | | | **MD** | | |
| --- | --- | --- | --- | --- | --- | --- |
|  | **Beta coef** | **p-value** | **Cohens’d** | **Beta coef** | **p-value** | **Cohen’s d** |
| Atr | 0.000736 | 0.222 | 0.0255 | 0.000045 | 0.949 | 0.0013 |
| Cgc | 0.000378 | 0.671 | 0.0088 | -0.000363 | 0.662 | -0.0091 |
| Cgh | -0.000713 | 0.432 | -0.0164 | 0.000613 | 0.545 | 0.0126 |
| Cst | 0.000864 | 0.129 | 0.0317 | 0.000935 | 0.116 | 0.0328 |
| Fscs | 0.001608 | 0.003 | 0.0626 | 0.000289 | 0.549 | 0.0125 |
| Fxcut | 0.000533 | 0.492 | 0.0143 | 0.002033 | 0.308 | 0.0212 |
| Fx | 0.000077 | 0.906 | 0.0024 | 0.001829 | 0.212 | 0.026 |
| Ifo | 0.000101 | 0.86 | 0.0037 | 0.000889 | 0.125 | 0.032 |
| Ifsfc | 0.000954 | 0.061 | 0.0391 | 0.000393 | 0.47 | 0.015 |
| Ilf | 0.000613 | 0.297 | 0.0217 | 0.000256 | 0.687 | 0.0084 |
| Pscs | 0.000856 | 0.109 | 0.0335 | 0.000323 | 0.525 | 0.0132 |
| Pslf | 0.000253 | 0.611 | 0.0106 | 0.000542 | 0.315 | 0.0209 |
| Scs | 0.00121 | 0.017 | 0.0501 | 0.000362 | 0.454 | 0.0156 |
| Sifc | -0.000075 | 0.909 | -0.0024 | 0.000183 | 0.77 | 0.0061 |
| Slf | 0.000414 | 0.385 | 0.0181 | 0.000471 | 0.362 | 0.019 |
| Tslf | 0.00036 | 0.448 | 0.0158 | 0.000493 | 0.33 | 0.0203 |
| Unc | 0.000296 | 0.642 | 0.0097 | 0.000824 | 0.186 | 0.0276 |
| allfibers | 0.000398 | 0.326 | 0.0205 | 0.000732 | 0.146 | 0.0303 |

##### *Models with winsorised data and covariates*

Table 36: Beta coefficient, uncorrected p-value, and effect-sizes from the models investigating associations between cortical GM features and E2 timing

| **ROI** | **Thickness** | | | **Sulcal Depth** | | | **Surface area** | | | **Cortical volume** | | |
| --- | --- | --- | --- | --- | --- | --- | --- | --- | --- | --- | --- | --- |
|  | Beta coef | p-value | Cohen’s d | Beta coef | p-value | Cohen’s d | Beta coef | p-value | Cohen’s d | Beta coef | p-value | Cohen’s d |
| bankssts | 0.000024 | 0.668 | 0.007 | -0.000106 | 0.503 | -0.012 | -0.088205 | 0.023 | -0.039 | -0.209314 | 0.089 | -0.029 |
| caudal anterior cingulate | 0.000036 | 0.611 | 0.009 | 0.000044 | 0.688 | 0.007 | -0.034160 | 0.184 | -0.023 | -0.052763 | 0.597 | -0.009 |
| caudal middle frontal | -0.000010 | 0.882 | -0.003 | -0.000054 | 0.678 | -0.007 | -0.160447 | 0.151 | -0.025 | -0.848218 | 0.020 | -0.040 |
| cuneus | 0.000040 | 0.496 | 0.012 | 0.000155 | 0.205 | 0.022 | 0.007547 | 0.871 | 0.003 | 0.091942 | 0.535 | 0.011 |
| entorhinal | 0.000104 | 0.524 | 0.011 | 0.000029 | 0.927 | 0.002 | 0.024647 | 0.422 | 0.014 | 0.255464 | 0.088 | 0.029 |
| fusiform | -0.000020 | 0.695 | -0.007 | -0.000024 | 0.789 | -0.005 | -0.090736 | 0.174 | -0.023 | -0.519302 | 0.079 | -0.030 |
| inferior parietal | -0.000001 | 0.983 | 0.000 | 0.000076 | 0.278 | 0.019 | -0.028392 | 0.859 | -0.003 | -0.361592 | 0.475 | -0.012 |
| inferior temporal | -0.000049 | 0.417 | -0.014 | -0.000019 | 0.825 | -0.004 | -0.133495 | 0.143 | -0.025 | -0.552074 | 0.135 | -0.026 |
| isthmus cingulate | -0.000002 | 0.973 | -0.001 | -0.000072 | 0.533 | -0.011 | 0.007783 | 0.793 | 0.005 | 0.010913 | 0.913 | 0.002 |
| lateral occipital | -0.000020 | 0.713 | -0.006 | -0.000060 | 0.410 | -0.014 | -0.061890 | 0.597 | -0.009 | -0.401163 | 0.308 | -0.018 |
| lateral orbitofrontal | -0.000035 | 0.617 | -0.009 | -0.000105 | 0.321 | -0.017 | -0.036765 | 0.765 | -0.005 | -0.153737 | 0.664 | -0.007 |
| lingual | 0.000042 | 0.443 | 0.013 | 0.000065 | 0.448 | 0.013 | -0.000520 | 0.994 | 0.000 | 0.312916 | 0.158 | 0.024 |
| Medial orbitofrontal | -0.000082 | 0.266 | -0.019 | 0.000193 | 0.113 | 0.027 | -0.134606 | 0.142 | -0.025 | -0.521798 | 0.064 | -0.032 |
| middle temporal | 0.000014 | 0.823 | 0.004 | 0.000036 | 0.673 | 0.007 | -0.249405 | 0.012 | -0.044 | -0.403644 | 0.255 | -0.020 |
| parahippocampal | -0.000057 | 0.466 | -0.013 | 0.000012 | 0.935 | 0.001 | 0.013377 | 0.603 | 0.009 | -0.030036 | 0.756 | -0.005 |
| paracentral | -0.000008 | 0.911 | -0.002 | 0.000028 | 0.806 | 0.004 | 0.024543 | 0.577 | 0.010 | 0.033093 | 0.857 | 0.003 |
| pars opercularis | -0.000021 | 0.694 | -0.007 | 0.000117 | 0.497 | 0.012 | -0.038021 | 0.531 | -0.011 | -0.196755 | 0.372 | -0.015 |
| pars orbitalis | -0.000007 | 0.922 | -0.002 | -0.000045 | 0.741 | -0.006 | -0.017076 | 0.509 | -0.011 | -0.124328 | 0.275 | -0.019 |
| pars triangularis | -0.000010 | 0.875 | -0.003 | 0.000086 | 0.453 | 0.013 | -0.057701 | 0.320 | -0.017 | -0.178783 | 0.380 | -0.015 |
| pericalcarine | 0.000109 | 0.125 | 0.026 | -0.000020 | 0.882 | -0.003 | -0.054970 | 0.283 | -0.019 | 0.132964 | 0.246 | 0.020 |
| postcentral | -0.000108 | 0.088 | -0.029 | 0.000066 | 0.477 | 0.012 | 0.280411 | 0.038 | 0.036 | 0.078971 | 0.844 | 0.003 |
| posterior cingulate | -0.000031 | 0.494 | -0.012 | 0.000080 | 0.457 | 0.013 | -0.008719 | 0.781 | -0.005 | 0.018050 | 0.865 | 0.003 |
| precentral | 0.000025 | 0.719 | 0.006 | 0.000103 | 0.152 | 0.025 | -0.015932 | 0.923 | -0.002 | 0.040839 | 0.933 | 0.001 |
| precuneus | -0.000024 | 0.624 | -0.008 | 0.000116 | 0.109 | 0.028 | -0.043017 | 0.655 | -0.008 | -0.151188 | 0.686 | -0.007 |
| Rostral anterior cingulate | 0.000133 | 0.184 | 0.023 | -0.000075 | 0.615 | -0.009 | -0.049501 | 0.197 | -0.022 | 0.085895 | 0.564 | 0.010 |
| rostral middle frontal | 0.000053 | 0.392 | 0.015 | -0.000148 | 0.048 | -0.034 | -0.167723 | 0.483 | -0.012 | -0.620864 | 0.401 | -0.014 |
| superior frontal | 0.000035 | 0.572 | 0.010 | -0.000043 | 0.444 | -0.013 | -0.467402 | 0.058 | -0.033 | -1.585715 | 0.066 | -0.032 |
| superior parietal | 0.000003 | 0.967 | 0.001 | -0.000050 | 0.540 | -0.011 | 0.194545 | 0.365 | 0.016 | -0.067626 | 0.923 | -0.002 |
| Superior temporal | -0.000033 | 0.585 | -0.009 | 0.000045 | 0.487 | 0.012 | -0.184747 | 0.053 | -0.033 | -0.529263 | 0.145 | -0.025 |
| supramarginal | 0.000007 | 0.911 | 0.002 | -0.000005 | 0.960 | -0.001 | -0.065792 | 0.695 | -0.007 | 0.273360 | 0.601 | 0.009 |
| frontal pole | 0.000344 | 0.005 | 0.049 | -0.000560 | 0.043 | -0.035 | -0.010963 | 0.534 | -0.011 | 0.096351 | 0.328 | 0.017 |
| temporal pole | 0.000307 | 0.083 | 0.030 | 0.000107 | 0.748 | 0.006 | -0.035342 | 0.176 | -0.023 | -0.009823 | 0.958 | -0.001 |
| transverse temporal | -0.000107 | 0.175 | -0.023 | 0.000169 | 0.332 | 0.017 | -0.003214 | 0.823 | -0.004 | -0.076298 | 0.181 | -0.023 |
| insula | -0.000003 | 0.973 | -0.001 | -0.000593 | 0.013 | -0.043 | 0.200534 | 0.049 | 0.034 | 0.553969 | 0.082 | 0.030 |
| Global | -0.000002 | 0.952 | -0.001 | 0.000004 | 0.602 | 0.009 | -3.147380 | 0.240 | -0.020 | -13.089679 | 0.230 | -0.021 |

Table 37: Beta coefficient, uncorrected p-value, and effect-sizes from the models investigating associations between sub-cortical GM features and E2 timing

| **Regions** | **Beta coefficient** | **p-value** | **Cohen’s d** |
| --- | --- | --- | --- |
| Thalamus proper | -0.077019 | 0.766 | -0.005 |
| Caudate | -0.110902 | 0.228 | -0.021 |
| Putamen | 0.088178 | 0.552 | 0.010 |
| Pallidum | -0.086938 | 0.427 | -0.014 |
| Hippocampus | -0.134952 | 0.179 | -0.023 |
| Amygdala | -0.046183 | 0.570 | -0.010 |
| Accumbens area | 0.016373 | 0.740 | 0.006 |
| Total subcortical volume | -1.184773 | 0.153 | -0.025 |

Table 38: Beta coefficient, uncorrected p-value, and effect-sizes from the models investigating associations between WM features and E2 timing

| **ROI** | **FA** | | | **MD** | | |
| --- | --- | --- | --- | --- | --- | --- |
|  | **Beta coef** | **p-value** | **Cohens’d** | **Beta coef** | **p-value** | **Cohen’s d** |
| Atr | -0.000017 | 0.240 | -0.022 | 0.000001 | 0.951 | 0.001 |
| Cgc | -0.000021 | 0.304 | -0.019 | -0.000002 | 0.920 | -0.002 |
| Cgh | -0.000015 | 0.485 | -0.013 | -0.000021 | 0.374 | -0.016 |
| Cst | 0.000011 | 0.390 | 0.016 | -0.000012 | 0.392 | -0.016 |
| Fscs | -0.000015 | 0.223 | -0.022 | 0.000000 | 0.986 | 0.000 |
| Fxcut | -0.000002 | 0.902 | -0.002 | -0.000043 | 0.355 | -0.017 |
| Fx | -0.000008 | 0.610 | -0.009 | -0.000029 | 0.396 | -0.016 |
| Ifo | -0.000018 | 0.165 | -0.026 | -0.000002 | 0.909 | -0.002 |
| Ifsfc | -0.000002 | 0.875 | -0.003 | 0.000000 | 0.978 | 0.001 |
| Ilf | -0.000027 | 0.051 | -0.036 | -0.000004 | 0.772 | -0.005 |
| Pscs | -0.000005 | 0.682 | -0.008 | 0.000005 | 0.650 | 0.008 |
| Pslf | -0.000003 | 0.799 | -0.005 | -0.000003 | 0.819 | -0.004 |
| Scs | -0.000011 | 0.339 | -0.018 | 0.000003 | 0.773 | 0.005 |
| Sifc | -0.000027 | 0.072 | -0.033 | 0.000011 | 0.477 | 0.013 |
| Slf | -0.000006 | 0.601 | -0.010 | 0.0000003 | 0.984 | 0.000 |
| Tslf | -0.000007 | 0.518 | -0.012 | 0.000002 | 0.889 | 0.003 |
| Unc | 0.000003 | 0.848 | 0.004 | -0.000007 | 0.649 | -0.008 |
| allfibers | -0.000011 | 0.254 | -0.021 | -0.000003 | 0.830 | -0.004 |

Table 39: Beta coefficient, uncorrected p-value, and effect-sizes from the models investigating associations between cortical GM features and E2 tempo

| **ROI** | **Thickness** | | | **Sulcal Depth** | | | **Surface area** | | | **Cortical volume** | | |
| --- | --- | --- | --- | --- | --- | --- | --- | --- | --- | --- | --- | --- |
|  | Beta coef | p-value | Cohen’s d | Beta coef | p-value | Cohen’s d | Beta coef | p-value | Cohen’s d | Beta coef | p-value | Cohen’s d |
| bankssts | -0.001804 | 0.147 | -0.028 | 0.000249 | 0.943 | 0.001 | 1.177692 | 0.177 | 0.026 | 1.493421 | 0.587 | 0.011 |
| caudal anterior cingulate | 0.000044 | 0.978 | 0.001 | 0.005269 | 0.036 | 0.041 | 0.370471 | 0.521 | 0.012 | 1.695858 | 0.454 | 0.015 |
| caudal middle frontal | -0.003155 | 0.038 | -0.040 | 0.002362 | 0.428 | 0.015 | 2.523548 | 0.321 | 0.019 | -0.016605 | 0.998 | 0.000 |
| cuneus | -0.001496 | 0.260 | -0.022 | -0.002977 | 0.284 | -0.021 | -1.101120 | 0.293 | -0.020 | -4.971663 | 0.132 | -0.029 |
| entorhinal | 0.001656 | 0.656 | 0.009 | 0.004138 | 0.568 | 0.011 | -0.235593 | 0.733 | -0.007 | -0.830329 | 0.806 | -0.005 |
| fusiform | -0.001711 | 0.150 | -0.028 | 0.000282 | 0.890 | 0.003 | -1.729651 | 0.253 | -0.022 | -11.935120 | 0.074 | -0.035 |
| inferior parietal | -0.003243 | 0.012 | -0.049 | -0.003658 | 0.023 | -0.044 | -0.281099 | 0.938 | -0.002 | -24.269806 | 0.036 | -0.041 |
| inferior temporal | -0.000855 | 0.539 | -0.012 | 0.000097 | 0.960 | 0.001 | -0.796493 | 0.701 | -0.007 | -8.928873 | 0.288 | -0.021 |
| isthmus cingulate | -0.000606 | 0.593 | -0.010 | 0.004855 | 0.061 | 0.036 | -1.241821 | 0.065 | -0.036 | -3.864261 | 0.090 | -0.033 |
| lateral occipital | -0.000274 | 0.827 | -0.004 | -0.001334 | 0.418 | -0.016 | 0.658011 | 0.804 | 0.005 | -2.770669 | 0.756 | -0.006 |
| lateral orbitofrontal | -0.000344 | 0.825 | -0.004 | 0.000102 | 0.965 | 0.001 | 0.762022 | 0.783 | 0.005 | -2.606562 | 0.745 | -0.006 |
| lingual | -0.000832 | 0.498 | -0.013 | 0.001210 | 0.530 | 0.012 | -3.697251 | 0.024 | -0.044 | -9.555173 | 0.053 | -0.038 |
| Medial orbitofrontal | 0.000726 | 0.666 | 0.008 | -0.002504 | 0.365 | -0.018 | -0.700979 | 0.735 | -0.007 | -0.250735 | 0.969 | -0.001 |
| middle temporal | -0.002664 | 0.064 | -0.036 | -0.001195 | 0.532 | -0.012 | 1.265496 | 0.574 | 0.011 | -10.470264 | 0.196 | -0.025 |
| parahippocampal | 0.002635 | 0.136 | 0.029 | 0.000229 | 0.947 | 0.001 | -0.323188 | 0.574 | -0.011 | 1.676791 | 0.445 | 0.015 |
| paracentral | -0.001359 | 0.372 | -0.017 | 0.000045 | 0.986 | 0.000 | -0.908409 | 0.355 | -0.018 | -4.313052 | 0.290 | -0.021 |
| pars opercularis | -0.002233 | 0.065 | -0.036 | -0.006201 | 0.121 | -0.030 | 0.824018 | 0.550 | 0.012 | 0.469642 | 0.925 | 0.002 |
| pars orbitalis | -0.000669 | 0.695 | -0.008 | -0.002168 | 0.485 | -0.014 | -1.451538 | 0.014 | -0.048 | -4.252982 | 0.098 | -0.032 |
| pars triangularis | -0.001258 | 0.372 | -0.017 | -0.000142 | 0.957 | -0.001 | -2.195481 | 0.102 | -0.032 | -9.890945 | 0.033 | -0.042 |
| pericalcarine | -0.002017 | 0.207 | -0.024 | 0.002035 | 0.508 | 0.013 | -0.692641 | 0.548 | -0.012 | -5.922470 | 0.020 | -0.045 |
| postcentral | 0.000963 | 0.498 | 0.013 | 0.000147 | 0.944 | 0.001 | -6.040887 | 0.048 | -0.038 | -14.320301 | 0.108 | -0.031 |
| posterior cingulate | -0.000781 | 0.444 | -0.015 | -0.000850 | 0.727 | -0.007 | -0.511715 | 0.467 | -0.014 | -2.827271 | 0.236 | -0.023 |
| precentral | -0.001878 | 0.240 | -0.023 | -0.004129 | 0.010 | -0.050 | -0.734540 | 0.846 | -0.004 | -17.958683 | 0.111 | -0.031 |
| precuneus | -0.001672 | 0.134 | -0.029 | -0.000385 | 0.814 | -0.005 | -2.625056 | 0.215 | -0.024 | -14.674885 | 0.076 | -0.035 |
| Rostral anterior cingulate | -0.002705 | 0.236 | -0.023 | 0.004613 | 0.165 | 0.027 | 0.268754 | 0.757 | 0.006 | -0.227693 | 0.946 | -0.001 |
| rostral middle frontal | -0.002785 | 0.048 | -0.039 | 0.002419 | 0.155 | 0.028 | -9.297720 | 0.086 | -0.033 | -35.022679 | 0.035 | -0.041 |
| superior frontal | -0.001961 | 0.163 | -0.027 | -0.000395 | 0.756 | -0.006 | 0.634679 | 0.910 | 0.002 | -17.504349 | 0.368 | -0.017 |
| superior parietal | -0.001751 | 0.236 | -0.023 | 0.002580 | 0.166 | 0.027 | -5.316480 | 0.268 | -0.021 | -19.213425 | 0.225 | -0.024 |
| Superior temporal | -0.000829 | 0.544 | -0.012 | -0.001750 | 0.238 | -0.023 | -0.297746 | 0.891 | -0.003 | -8.981307 | 0.273 | -0.021 |
| supramarginal | -0.003241 | 0.022 | -0.044 | -0.000623 | 0.767 | -0.006 | -0.226603 | 0.952 | -0.001 | -21.936846 | 0.064 | -0.036 |
| frontal pole | -0.007607 | 0.006 | -0.054 | 0.004710 | 0.450 | 0.015 | -0.313681 | 0.432 | -0.015 | -3.460688 | 0.118 | -0.030 |
| temporal pole | -0.007654 | 0.058 | -0.037 | 0.004604 | 0.544 | 0.012 | 0.321099 | 0.586 | 0.011 | -5.988865 | 0.155 | -0.028 |
| transverse temporal | 0.001700 | 0.340 | 0.019 | -0.000803 | 0.838 | -0.004 | -0.428887 | 0.187 | -0.026 | 0.425007 | 0.740 | 0.006 |
| insula | 0.000013 | 0.995 | 0.0001 | 0.005063 | 0.356 | 0.018 | -2.465316 | 0.294 | -0.020 | -4.469773 | 0.540 | -0.012 |
| Global | -0.001454 | 0.115 | -0.031 | -0.000074 | 0.632 | -0.009 | -82.849386 | 0.175 | -0.026 | -538.868696 | 0.028 | -0.043 |

Table 40: Beta coefficient, uncorrected p-value, and effect-sizes from the models investigating associations between sub-cortical GM features and E2 tempo

| **Regions** | **Beta coefficient** | **p-value** | **Cohen’s d** |
| --- | --- | --- | --- |
| Thalamus proper | 1.524346 | 0.794 | 0.005 |
| Caudate | -0.315310 | 0.880 | -0.003 |
| Putamen | -6.843268 | 0.045 | -0.039 |
| Pallidum | 3.161791 | 0.211 | 0.024 |
| Hippocampus | 1.242583 | 0.582 | 0.011 |
| Amygdala | 0.093208 | 0.959 | 0.001 |
| Accumbens area | -0.638853 | 0.570 | -0.011 |
| Total subcortical volume | -3.726178 | 0.841 | -0.004 |

Table 41: Beta coefficient, uncorrected p-value, and effect-sizes from the models investigating associations between WM features and E2 tempo

| **ROI** | **FA** | | | **MD** | | |
| --- | --- | --- | --- | --- | --- | --- |
|  | **Beta coef** | **p-value** | **Cohens’d** | **Beta coef** | **p-value** | **Cohen’s d** |
| Atr | 0.000343 | 0.291 | 0.022 | 0.000136 | 0.721 | 0.007 |
| Cgc | 0.000080 | 0.868 | 0.003 | -0.000095 | 0.832 | -0.004 |
| Cgh | -0.000307 | 0.531 | -0.013 | 0.000231 | 0.673 | 0.009 |
| Cst | 0.000335 | 0.274 | 0.023 | 0.000466 | 0.144 | 0.030 |
| Fscs | 0.000676 | 0.020 | 0.049 | 0.000145 | 0.575 | 0.012 |
| Fxcut | 0.000341 | 0.415 | 0.017 | 0.000790 | 0.462 | 0.015 |
| Fx | 0.000043 | 0.903 | 0.003 | 0.000828 | 0.293 | 0.022 |
| Ifo | 0.000097 | 0.752 | 0.007 | 0.000415 | 0.184 | 0.028 |
| Ifsfc | 0.000360 | 0.190 | 0.027 | 0.000210 | 0.475 | 0.015 |
| Ilf | 0.000208 | 0.511 | 0.014 | -0.000023 | 0.947 | -0.001 |
| Pscs | 0.000343 | 0.232 | 0.025 | 0.000080 | 0.770 | 0.006 |
| Pslf | 0.000009 | 0.974 | 0.001 | 0.000281 | 0.334 | 0.020 |
| Scs | 0.000511 | 0.060 | 0.039 | 0.000136 | 0.603 | 0.011 |
| Sifc | 0.000019 | 0.958 | 0.001 | 0.000198 | 0.559 | 0.012 |
| Slf | 0.000107 | 0.677 | 0.009 | 0.000220 | 0.429 | 0.016 |
| Tslf | 0.000096 | 0.709 | 0.008 | 0.000195 | 0.475 | 0.015 |
| Unc | 0.000144 | 0.675 | 0.009 | 0.000507 | 0.132 | 0.031 |
| allfibers | 0.000154 | 0.480 | 0.015 | 0.000384 | 0.157 | 0.030 |

#### 5. Exploratory Analyses

##### *Interaction Model*

Table 42: Beta coefficient, uncorrected p-value, and effect-sizes from the models investigating associations between cortical GM features and E2 timing*tempo

| **ROI** | **Thickness** | | | **Sulcal Depth** | | | **Surface area** | | | **Cortical volume** | | |
| --- | --- | --- | --- | --- | --- | --- | --- | --- | --- | --- | --- | --- |
|  | Beta coef | p-value | Cohen’s d | Beta coef | p-value | Cohen’s d | Beta coef | p-value | Cohen’s d | Beta coef | p-value | Cohen’s d |
| bankssts | 0.001508 | 0.206 | 0.025 | 0.000361 | 0.915 | 0.002 | -0.434004 | 0.606 | -0.010 | -1.758911 | 0.507 | -0.013 |
| caudal anterior cingulate | -0.000886 | 0.570 | -0.011 | 0.000367 | 0.882 | 0.003 | -0.295122 | 0.601 | -0.010 | -3.856300 | 0.078 | -0.034 |
| caudal middle frontal | 0.000381 | 0.797 | 0.005 | -0.000951 | 0.741 | -0.006 | -1.389499 | 0.575 | -0.011 | -4.573612 | 0.568 | -0.011 |
| cuneus | 0.002646 | 0.038 | 0.040 | -0.002601 | 0.331 | -0.019 | -0.651100 | 0.521 | -0.012 | 3.688494 | 0.255 | 0.022 |
| entorhinal | 0.001162 | 0.744 | 0.006 | -0.009634 | 0.168 | -0.027 | -0.771926 | 0.253 | -0.022 | -0.665403 | 0.841 | -0.004 |
| fusiform | 0.001048 | 0.367 | 0.018 | 0.001043 | 0.594 | 0.010 | 0.084258 | 0.954 | 0.001 | 3.607748 | 0.572 | 0.011 |
| inferior parietal | 0.001599 | 0.205 | 0.025 | 0.000685 | 0.657 | 0.009 | -2.626286 | 0.450 | -0.015 | -0.688502 | 0.950 | -0.001 |
| inferior temporal | 0.000812 | 0.551 | 0.012 | 0.001751 | 0.347 | 0.018 | -0.422877 | 0.831 | -0.004 | 2.803055 | 0.728 | 0.007 |
| isthmus cingulate | 0.000626 | 0.563 | 0.011 | 0.000222 | 0.929 | 0.002 | 0.393177 | 0.568 | 0.011 | 2.429367 | 0.273 | 0.021 |
| lateral occipital | 0.001294 | 0.283 | 0.021 | -0.002836 | 0.074 | -0.035 | 0.170003 | 0.947 | 0.001 | 7.402453 | 0.388 | 0.017 |
| lateral orbitofrontal | 0.000818 | 0.585 | 0.011 | 0.004035 | 0.076 | 0.035 | -0.048713 | 0.985 | 0.000 | -0.325102 | 0.966 | -0.001 |
| lingual | 0.002961 | 0.013 | 0.048 | 0.004882 | 0.009 | 0.051 | -0.805582 | 0.605 | -0.010 | 10.742364 | 0.025 | 0.044 |
| Medial orbitofrontal | 0.001177 | 0.468 | 0.014 | -0.000673 | 0.801 | -0.005 | -1.197538 | 0.548 | -0.012 | -4.087092 | 0.511 | -0.013 |
| middle temporal | 0.003780 | 0.009 | 0.051 | 0.001948 | 0.289 | 0.021 | -2.999229 | 0.175 | -0.026 | 6.844782 | 0.380 | 0.017 |
| parahippocampal | 0.002917 | 0.089 | 0.033 | 0.000037 | 0.991 | 0.000 | -0.889961 | 0.113 | -0.031 | 0.625355 | 0.770 | 0.006 |
| paracentral | 0.000922 | 0.528 | 0.012 | 0.002325 | 0.346 | 0.018 | 0.395858 | 0.678 | 0.008 | 1.708361 | 0.663 | 0.008 |
| pars opercularis | 0.000069 | 0.953 | 0.001 | -0.002032 | 0.603 | -0.010 | 2.347059 | 0.079 | 0.034 | 6.257608 | 0.195 | 0.025 |
| pars orbitalis | 0.000417 | 0.802 | 0.005 | 0.003823 | 0.204 | 0.025 | 0.015231 | 0.978 | 0.001 | -0.517645 | 0.835 | -0.004 |
| pars triangularis | 0.000766 | 0.574 | 0.011 | -0.006232 | 0.014 | -0.048 | 1.606310 | 0.211 | 0.024 | 7.816430 | 0.080 | 0.034 |
| pericalcarine | 0.001935 | 0.205 | 0.025 | 0.000680 | 0.825 | 0.004 | -1.130151 | 0.313 | -0.020 | 1.173718 | 0.635 | 0.009 |
| postcentral | 0.000483 | 0.725 | 0.007 | 0.000167 | 0.954 | 0.001 | 1.037284 | 0.730 | 0.007 | 0.153804 | 0.986 | 0.000 |
| posterior cingulate | 0.000168 | 0.865 | 0.003 | -0.000334 | 0.888 | -0.003 | 0.026520 | 0.969 | 0.001 | -1.039750 | 0.653 | -0.009 |
| precentral | 0.000042 | 0.979 | 0.001 | 0.001030 | 0.547 | 0.012 | -3.408396 | 0.354 | -0.018 | -8.205435 | 0.453 | -0.015 |
| precuneus | 0.001205 | 0.266 | 0.022 | -0.000251 | 0.875 | -0.003 | -1.424720 | 0.486 | -0.014 | -0.584955 | 0.941 | -0.001 |
| Rostral anterior cingulate | 0.001908 | 0.382 | 0.017 | -0.003539 | 0.271 | -0.021 | 0.652244 | 0.431 | 0.015 | 4.875353 | 0.133 | 0.029 |
| rostral middle frontal | 0.000897 | 0.512 | 0.013 | 0.001985 | 0.224 | 0.024 | 8.815358 | 0.089 | 0.033 | 25.930683 | 0.103 | 0.032 |
| superior frontal | -0.000118 | 0.931 | -0.002 | -0.000504 | 0.680 | -0.008 | 0.476165 | 0.930 | 0.002 | -6.824280 | 0.714 | -0.007 |
| superior parietal | 0.002306 | 0.104 | 0.032 | -0.000412 | 0.819 | -0.004 | -3.656276 | 0.430 | -0.015 | 6.682036 | 0.661 | 0.009 |
| Superior temporal | 0.001189 | 0.364 | 0.018 | -0.001830 | 0.205 | -0.025 | 0.238926 | 0.911 | 0.002 | -0.212367 | 0.978 | -0.001 |
| supramarginal | 0.001970 | 0.157 | 0.027 | -0.002585 | 0.221 | -0.024 | -1.757797 | 0.626 | -0.009 | 11.216699 | 0.324 | 0.019 |
| frontal pole | 0.002343 | 0.383 | 0.017 | 0.005506 | 0.362 | 0.018 | 0.228783 | 0.554 | 0.011 | 2.092869 | 0.332 | 0.019 |
| temporal pole | -0.000081 | 0.984 | 0.000 | -0.006333 | 0.390 | -0.017 | -0.181768 | 0.751 | -0.006 | -2.055641 | 0.623 | -0.010 |
| transverse temporal | 0.000856 | 0.621 | 0.010 | -0.000979 | 0.796 | -0.005 | -0.095830 | 0.762 | -0.006 | -0.539937 | 0.663 | -0.008 |
| insula | 0.001378 | 0.488 | 0.013 | -0.005249 | 0.323 | -0.019 | 1.235561 | 0.585 | 0.011 | 7.705730 | 0.270 | 0.021 |
| Global | 0.001216 | 0.172 | 0.027 | -0.000045 | 0.767 | -0.006 | -12.288349 | 0.834 | -0.004 | 98.435868 | 0.675 | 0.008 |

Table 43: Beta coefficient, uncorrected p-value, and effect-sizes from the models investigating associations between sub-cortical GM features and E2 timing*tempo

| **Regions** | **Beta coefficient** | **p-value** | **Cohen’s d** |
| --- | --- | --- | --- |
| Thalamus proper | -6.113177 | 0.275 | -0.021 |
| Caudate | 1.905509 | 0.345 | 0.018 |
| Putamen | 6.262062 | 0.056 | 0.037 |
| Pallidum | -1.798385 | 0.470 | -0.014 |
| Hippocampus | 1.528327 | 0.483 | 0.014 |
| Amygdala | -1.733418 | 0.332 | -0.019 |
| Accumbens area | -0.280146 | 0.798 | -0.005 |
| Total subcortical volume | -3.157981 | 0.859 | -0.003 |

Table 44: Beta coefficient, uncorrected p-value, and effect-sizes from the models investigating associations between WM features and E2 timing*tempo

| **ROI** | **FA** | | | **MD** | | |
| --- | --- | --- | --- | --- | --- | --- |
|  | **Beta coef** | **p-value** | **Cohens’d** | **Beta coef** | **p-value** | **Cohen’s d** |
| Atr | -0.00014 | 0.643 | -0.010 | 0.00010 | 0.786 | 0.006 |
| Cgc | -0.00035 | 0.420 | -0.017 | 0.00018 | 0.665 | 0.009 |
| Cgh | 0.00003 | 0.955 | 0.001 | 0.00059 | 0.249 | 0.024 |
| Cst | -0.00009 | 0.760 | -0.006 | 0.00049 | 0.099 | 0.034 |
| Fscs | -0.00038 | 0.148 | -0.030 | 0.00025 | 0.298 | 0.022 |
| Fxcut | 0.00016 | 0.678 | 0.009 | 0.00074 | 0.467 | 0.015 |
| Fx | 0.00019 | 0.553 | 0.012 | 0.00066 | 0.378 | 0.018 |
| Ifo | 0.00015 | 0.591 | 0.011 | 0.00049 | 0.086 | 0.036 |
| Ifsfc | -0.00005 | 0.851 | -0.004 | 0.00037 | 0.168 | 0.029 |
| Ilf | -0.00016 | 0.573 | -0.012 | 0.00056 | 0.075 | 0.037 |
| Pscs | -0.00026 | 0.315 | -0.021 | 0.00030 | 0.231 | 0.025 |
| Pslf | -0.00024 | 0.333 | -0.020 | 0.00019 | 0.481 | 0.015 |
| Scs | -0.00034 | 0.173 | -0.028 | 0.00033 | 0.176 | 0.028 |
| Sifc | 0.00026 | 0.430 | 0.016 | 0.00067 | 0.032 | 0.045 |
| Slf | -0.00012 | 0.614 | -0.010 | 0.00018 | 0.482 | 0.015 |
| Tslf | -0.00005 | 0.830 | -0.004 | 0.00023 | 0.364 | 0.019 |
| Unc | 0.00001 | 0.964 | 0.001 | 0.00009 | 0.784 | 0.006 |
| allfibers | 0.000003 | 0.988 | 0.0003 | 0.00024 | 0.346 | 0.020 |

##### *Interaction Model with covariates*

Table 45: Beta coefficient, uncorrected p-value, and effect-sizes from the models investigating associations between cortical GM features and E2 timing*tempo

| **ROI** | **Thickness** | | | **Sulcal Depth** | | | **Surface area** | | | **Cortical volume** | | |
| --- | --- | --- | --- | --- | --- | --- | --- | --- | --- | --- | --- | --- |
|  | Beta coef | p-value | Cohen’s d | Beta coef | p-value | Cohen’s d | Beta coef | p-value | Cohen’s d | Beta coef | p-value | Cohen’s d |
| bankssts | 0.000817 | 0.202 | 0.025 | 0.000658 | 0.717 | 0.007 | -0.197448 | 0.661 | -0.009 | -0.875229 | 0.538 | -0.012 |
| caudal anterior cingulate | -0.000657 | 0.432 | -0.015 | 0.000269 | 0.840 | 0.004 | -0.150816 | 0.618 | -0.010 | -2.151882 | 0.067 | -0.036 |
| caudal middle frontal | 0.000473 | 0.552 | 0.012 | -0.000741 | 0.632 | -0.009 | -0.463120 | 0.727 | -0.007 | -0.486295 | 0.910 | -0.002 |
| cuneus | 0.001538 | 0.025 | 0.044 | -0.001367 | 0.340 | -0.018 | -0.367083 | 0.499 | -0.013 | 2.165835 | 0.211 | 0.024 |
| entorhinal | 0.000435 | 0.820 | 0.004 | -0.005041 | 0.178 | -0.026 | -0.427816 | 0.237 | -0.023 | -0.631082 | 0.722 | -0.007 |
| fusiform | 0.000540 | 0.386 | 0.017 | 0.000585 | 0.577 | 0.011 | 0.046869 | 0.952 | 0.001 | 1.892433 | 0.580 | 0.011 |
| inferior parietal | 0.000891 | 0.187 | 0.026 | 0.000371 | 0.654 | 0.009 | -0.981612 | 0.596 | -0.010 | 0.946806 | 0.873 | 0.003 |
| inferior temporal | 0.000469 | 0.520 | 0.012 | 0.001265 | 0.205 | 0.025 | -0.041749 | 0.969 | -0.001 | 2.199235 | 0.609 | 0.010 |
| isthmus cingulate | 0.000324 | 0.576 | 0.011 | -0.000323 | 0.809 | -0.005 | 0.262250 | 0.477 | 0.014 | 1.355369 | 0.253 | 0.022 |
| lateral occipital | 0.000765 | 0.236 | 0.023 | -0.001621 | 0.057 | -0.037 | 0.326952 | 0.810 | 0.005 | 5.081042 | 0.268 | 0.021 |
| lateral orbitofrontal | 0.000414 | 0.606 | 0.010 | 0.002093 | 0.085 | 0.033 | 0.028113 | 0.984 | 0.000 | -0.023656 | 0.995 | 0.000 |
| lingual | 0.001790 | 0.005 | 0.055 | 0.002816 | 0.005 | 0.055 | -0.511335 | 0.540 | -0.012 | 6.543831 | 0.011 | 0.050 |
| Medial orbitofrontal | 0.000580 | 0.505 | 0.013 | -0.000120 | 0.933 | -0.002 | -0.477347 | 0.655 | -0.009 | -1.494905 | 0.654 | -0.009 |
| middle temporal | 0.002085 | 0.007 | 0.053 | 0.001024 | 0.298 | 0.020 | -1.270561 | 0.281 | -0.021 | 5.271391 | 0.206 | 0.025 |
| parahippocampal | 0.001529 | 0.096 | 0.032 | 0.000104 | 0.954 | 0.001 | -0.484939 | 0.107 | -0.031 | 0.257903 | 0.822 | 0.004 |
| paracentral | 0.000741 | 0.344 | 0.018 | 0.001250 | 0.345 | 0.018 | 0.320950 | 0.529 | 0.012 | 1.765021 | 0.400 | 0.016 |
| pars opercularis | 0.000150 | 0.810 | 0.005 | -0.001023 | 0.625 | -0.009 | 1.286939 | 0.072 | 0.035 | 3.799203 | 0.142 | 0.028 |
| pars orbitalis | 0.000117 | 0.896 | 0.003 | 0.002265 | 0.161 | 0.027 | -0.002253 | 0.994 | 0.000 | -0.302833 | 0.820 | -0.004 |
| pars triangularis | 0.000502 | 0.493 | 0.013 | -0.003289 | 0.016 | -0.047 | 0.764291 | 0.267 | 0.022 | 4.189655 | 0.080 | 0.034 |
| pericalcarine | 0.001264 | 0.122 | 0.030 | 0.000417 | 0.800 | 0.005 | -0.649847 | 0.277 | -0.021 | 0.918846 | 0.488 | 0.013 |
| postcentral | 0.000434 | 0.555 | 0.011 | 0.000026 | 0.986 | 0.0001 | 0.639924 | 0.691 | 0.008 | 1.605128 | 0.728 | 0.007 |
| posterior cingulate | 0.000063 | 0.905 | 0.002 | -0.000366 | 0.773 | -0.006 | 0.087984 | 0.808 | 0.005 | -0.492214 | 0.690 | -0.008 |
| precentral | 0.000138 | 0.870 | 0.003 | 0.000426 | 0.643 | 0.009 | -1.532123 | 0.437 | -0.015 | -2.347411 | 0.688 | -0.008 |
| precuneus | 0.000700 | 0.227 | 0.023 | -0.000040 | 0.963 | -0.001 | -0.696065 | 0.525 | -0.012 | 0.170272 | 0.968 | 0.001 |
| Rostral anterior cingulate | 0.001126 | 0.336 | 0.019 | -0.001925 | 0.265 | -0.022 | 0.283011 | 0.524 | 0.012 | 2.580258 | 0.138 | 0.029 |
| rostral middle frontal | 0.000601 | 0.412 | 0.016 | 0.001090 | 0.213 | 0.024 | 4.375593 | 0.115 | 0.031 | 13.909456 | 0.102 | 0.032 |
| superior frontal | 0.000150 | 0.837 | 0.004 | -0.000170 | 0.795 | -0.005 | 0.781867 | 0.788 | 0.005 | 0.891875 | 0.929 | 0.002 |
| superior parietal | 0.001314 | 0.084 | 0.034 | -0.000062 | 0.949 | -0.001 | -1.924610 | 0.438 | -0.015 | 4.533197 | 0.578 | 0.011 |
| Superior temporal | 0.000686 | 0.328 | 0.019 | -0.000985 | 0.203 | -0.025 | 0.371730 | 0.746 | 0.006 | 0.955867 | 0.820 | 0.004 |
| supramarginal | 0.001199 | 0.107 | 0.031 | -0.001396 | 0.219 | -0.024 | -1.097290 | 0.570 | -0.011 | 6.951686 | 0.252 | 0.022 |
| frontal pole | 0.001041 | 0.470 | 0.014 | 0.002908 | 0.369 | 0.017 | 0.143664 | 0.489 | 0.013 | 1.221051 | 0.292 | 0.020 |
| temporal pole | -0.000025 | 0.991 | 0.000 | -0.003945 | 0.318 | -0.019 | -0.013229 | 0.966 | -0.001 | -0.646050 | 0.773 | -0.006 |
| transverse temporal | 0.000567 | 0.541 | 0.012 | -0.000113 | 0.956 | -0.001 | -0.034819 | 0.837 | -0.004 | -0.121746 | 0.855 | -0.004 |
| insula | 0.000709 | 0.506 | 0.013 | -0.003019 | 0.290 | -0.021 | 0.663104 | 0.584 | 0.011 | 3.953519 | 0.290 | 0.021 |
| Global | 0.000740 | 0.121 | 0.030 | -0.000018 | 0.820 | -0.004 | -0.377609 | 0.990 | 0.0002 | 96.727506 | 0.439 | 0.015 |

Table 46: Beta coefficient, uncorrected p-value, and effect-sizes from the models investigating associations between sub-cortical GM features and E2 timing*tempo

| **Regions** | **Beta coefficient** | **p-value** | **Cohen’s d** |
| --- | --- | --- | --- |
| Thalamus proper | -2.432839 | 0.417 | -0.016 |
| Caudate | 1.344839 | 0.214 | 0.024 |
| Putamen | 3.603294 | 0.040 | 0.040 |
| Pallidum | -0.881842 | 0.509 | -0.013 |
| Hippocampus | 0.882741 | 0.450 | 0.015 |
| Amygdala | -0.849620 | 0.375 | -0.017 |
| Accumbens area | 0.090200 | 0.878 | 0.003 |
| Total subcortical volume | 3.113733 | 0.744 | 0.006 |

Table 47: Beta coefficient, uncorrected p-value, and effect-sizes from the models investigating associations between WM features and E2 timing*tempo

| **ROI** | **FA** | | | **MD** | | |
| --- | --- | --- | --- | --- | --- | --- |
|  | **Beta coef** | **p-value** | **Cohens’d** | **Beta coef** | **p-value** | **Cohen’s d** |
| Atr | -0.000060 | 0.707 | -0.008 | 0.000012 | 0.950 | 0.001 |
| Cgc | -0.000108 | 0.645 | -0.010 | 0.000068 | 0.762 | 0.006 |
| Cgh | 0.000093 | 0.699 | 0.008 | 0.000291 | 0.290 | 0.022 |
| Cst | 0.000003 | 0.982 | 0.000 | 0.000212 | 0.181 | 0.028 |
| Fscs | -0.000183 | 0.194 | -0.027 | 0.000095 | 0.456 | 0.016 |
| Fxcut | 0.000108 | 0.598 | 0.011 | 0.000305 | 0.571 | 0.012 |
| Fx | 0.000149 | 0.385 | 0.018 | 0.000269 | 0.498 | 0.014 |
| Ifo | 0.000116 | 0.443 | 0.016 | 0.000215 | 0.162 | 0.029 |
| Ifsfc | 0.000019 | 0.887 | 0.003 | 0.000132 | 0.360 | 0.019 |
| Ilf | -0.000064 | 0.680 | -0.009 | 0.000286 | 0.087 | 0.036 |
| Pscs | -0.000143 | 0.309 | -0.021 | 0.000141 | 0.296 | 0.022 |
| Pslf | -0.000096 | 0.463 | -0.015 | 0.000075 | 0.599 | 0.011 |
| Scs | -0.000179 | 0.175 | -0.028 | 0.000148 | 0.249 | 0.024 |
| Sifc | 0.000176 | 0.307 | 0.021 | 0.000333 | 0.047 | 0.041 |
| Slf | -0.000043 | 0.732 | -0.007 | 0.000075 | 0.582 | 0.011 |
| Tslf | -0.000017 | 0.891 | -0.003 | 0.000105 | 0.431 | 0.016 |
| Unc | 0.000028 | 0.869 | 0.003 | 0.000039 | 0.816 | 0.005 |
| allfibers | 0.000039 | 0.717 | 0.008 | 0.000074 | 0.580 | 0.012 |

##### *Interaction Model with winsorised data*

Table 48: Beta coefficient, uncorrected p-value, and effect-sizes from the models investigating associations between cortical GM features and E2 timing*tempo

| **ROI** | **Thickness** | | | **Sulcal Depth** | | | **Surface area** | | | **Cortical volume** | | |
| --- | --- | --- | --- | --- | --- | --- | --- | --- | --- | --- | --- | --- |
|  | Beta coef | p-value | Cohen’s d | Beta coef | p-value | Cohen’s d | Beta coef | p-value | Cohen’s d | Beta coef | p-value | Cohen’s d |
| bankssts | 0.002220 | 0.227 | 0.023 | -0.003360 | 0.516 | -0.013 | -0.564799 | 0.661 | -0.009 | -3.778706 | 0.352 | -0.018 |
| caudal anterior cingulate | -0.000706 | 0.768 | -0.006 | 0.001843 | 0.621 | 0.010 | -0.220041 | 0.796 | -0.005 | -2.981140 | 0.373 | -0.017 |
| caudal middle frontal | -0.001001 | 0.656 | -0.009 | -0.002305 | 0.603 | -0.010 | -0.709542 | 0.850 | -0.004 | -8.988317 | 0.465 | -0.014 |
| cuneus | 0.001332 | 0.497 | 0.013 | 0.002167 | 0.598 | 0.010 | -1.832999 | 0.238 | -0.023 | -0.511084 | 0.917 | -0.002 |
| entorhinal | 0.004338 | 0.426 | 0.015 | -0.008213 | 0.444 | -0.015 | -1.458898 | 0.154 | -0.028 | -3.914483 | 0.433 | -0.015 |
| fusiform | 0.002682 | 0.127 | 0.030 | -0.001927 | 0.522 | -0.012 | -1.111866 | 0.617 | -0.010 | 6.479316 | 0.509 | 0.013 |
| inferior parietal | 0.001915 | 0.309 | 0.020 | 0.001467 | 0.536 | 0.012 | -8.092579 | 0.129 | -0.030 | -17.968178 | 0.293 | -0.020 |
| inferior temporal | 0.001338 | 0.514 | 0.013 | 0.001237 | 0.666 | 0.008 | -0.141999 | 0.963 | -0.001 | -0.433162 | 0.972 | -0.001 |
| isthmus cingulate | 0.000442 | 0.791 | 0.005 | 0.001614 | 0.675 | 0.008 | 0.302226 | 0.763 | 0.006 | 2.429770 | 0.471 | 0.014 |
| lateral occipital | 0.001058 | 0.567 | 0.011 | -0.003579 | 0.143 | -0.028 | -1.233935 | 0.752 | -0.006 | 1.856465 | 0.888 | 0.003 |
| lateral orbitofrontal | 0.001631 | 0.477 | 0.014 | 0.000932 | 0.789 | 0.005 | -0.379096 | 0.926 | -0.002 | -0.776673 | 0.948 | -0.001 |
| lingual | 0.002914 | 0.110 | 0.031 | 0.007255 | 0.011 | 0.049 | -1.673320 | 0.485 | -0.014 | 6.832643 | 0.354 | 0.018 |
| Medial orbitofrontal | 0.003871 | 0.119 | 0.030 | 0.000399 | 0.923 | 0.002 | -4.123313 | 0.179 | -0.026 | -8.385173 | 0.379 | -0.017 |
| middle temporal | 0.002980 | 0.157 | 0.027 | 0.002905 | 0.303 | 0.020 | -5.229147 | 0.116 | -0.031 | -4.480141 | 0.707 | -0.007 |
| parahippocampal | 0.003296 | 0.213 | 0.024 | 0.002111 | 0.680 | 0.008 | 0.071959 | 0.933 | 0.002 | 2.275518 | 0.489 | 0.013 |
| paracentral | -0.000073 | 0.974 | -0.001 | 0.000169 | 0.964 | 0.001 | -0.727384 | 0.619 | -0.010 | -4.377389 | 0.468 | -0.014 |
| pars opercularis | 0.000267 | 0.881 | 0.003 | -0.010438 | 0.079 | -0.034 | 0.352301 | 0.862 | 0.003 | -3.108017 | 0.672 | -0.008 |
| pars orbitalis | -0.000185 | 0.942 | -0.001 | 0.005673 | 0.220 | 0.024 | -0.238695 | 0.783 | -0.005 | 0.479384 | 0.900 | 0.002 |
| pars triangularis | -0.000116 | 0.955 | -0.001 | -0.006924 | 0.075 | -0.035 | 1.732073 | 0.381 | 0.017 | 4.410287 | 0.519 | 0.013 |
| pericalcarine | -0.000846 | 0.719 | -0.007 | 0.001518 | 0.739 | 0.006 | -3.221704 | 0.060 | -0.037 | -5.794900 | 0.125 | -0.030 |
| postcentral | 0.000534 | 0.798 | 0.005 | -0.001961 | 0.531 | -0.012 | -4.225874 | 0.347 | -0.018 | -16.258115 | 0.217 | -0.024 |
| posterior cingulate | -0.000713 | 0.637 | -0.009 | 0.002264 | 0.530 | 0.012 | -0.943856 | 0.363 | -0.018 | -5.965811 | 0.093 | -0.033 |
| precentral | -0.001520 | 0.519 | -0.012 | 0.003156 | 0.183 | 0.026 | -13.019412 | 0.020 | -0.045 | -39.564408 | 0.017 | -0.046 |
| precuneus | 0.002156 | 0.189 | 0.025 | -0.000224 | 0.927 | -0.002 | -4.399712 | 0.163 | -0.027 | -10.654292 | 0.379 | -0.017 |
| Rostral anterior cingulate | -0.002138 | 0.525 | -0.012 | -0.006023 | 0.223 | -0.024 | 2.175152 | 0.089 | 0.033 | 5.437522 | 0.276 | 0.021 |
| rostral middle frontal | 0.000163 | 0.937 | 0.002 | 0.003801 | 0.130 | 0.029 | 10.433517 | 0.190 | 0.025 | 17.991710 | 0.461 | 0.014 |
| superior frontal | -0.001436 | 0.489 | -0.013 | -0.001447 | 0.440 | -0.015 | -2.057764 | 0.803 | -0.005 | -28.743333 | 0.317 | -0.019 |
| superior parietal | 0.002701 | 0.214 | 0.024 | 0.000028 | 0.992 | 0.000 | -10.635690 | 0.134 | -0.029 | -10.647829 | 0.648 | -0.009 |
| Superior temporal | 0.001416 | 0.481 | 0.014 | -0.003644 | 0.101 | -0.032 | -2.104342 | 0.512 | -0.013 | -13.036802 | 0.281 | -0.021 |
| supramarginal | 0.001687 | 0.417 | 0.016 | 0.001940 | 0.536 | 0.012 | -6.726185 | 0.224 | -0.024 | -3.969179 | 0.820 | -0.004 |
| frontal pole | 0.002671 | 0.513 | 0.013 | 0.017221 | 0.063 | 0.036 | 0.240853 | 0.685 | 0.008 | 0.526070 | 0.873 | 0.003 |
| temporal pole | 0.002691 | 0.650 | 0.009 | 0.001286 | 0.909 | 0.002 | 0.298665 | 0.732 | 0.007 | -1.853980 | 0.768 | -0.006 |
| transverse temporal | 0.000430 | 0.872 | 0.003 | 0.003693 | 0.527 | 0.012 | -0.534047 | 0.267 | -0.022 | -3.519242 | 0.065 | -0.036 |
| insula | 0.006175 | 0.044 | 0.039 | 0.007700 | 0.347 | 0.018 | -3.202638 | 0.356 | -0.018 | 0.813310 | 0.939 | 0.001 |
| Global | 0.001231 | 0.364 | 0.018 | 0.000200 | 0.383 | 0.017 | -129.116449 | 0.152 | -0.028 | -284.951030 | 0.431 | -0.015 |

Table 49: Beta coefficient, uncorrected p-value, and effect-sizes from the models investigating associations between sub-cortical GM features and E2 timing*tempo

| **Regions** | **Beta coefficient** | **p-value** | **Cohen’s d** |
| --- | --- | --- | --- |
| Thalamus proper | -6.613351 | 0.442 | -0.015 |
| Caudate | 1.047840 | 0.736 | 0.007 |
| Putamen | -3.955349 | 0.432 | -0.015 |
| Pallidum | 2.703025 | 0.468 | 0.014 |
| Hippocampus | 2.491499 | 0.455 | 0.014 |
| Amygdala | -4.777512 | 0.082 | -0.034 |
| Accumbens area | -1.774483 | 0.288 | -0.021 |
| Total subcortical volume | 1402.255529 | 0.263 | 0.022 |

Table 50: Beta coefficient, uncorrected p-value, and effect-sizes from the models investigating associations between WM features and E2 timing*tempo

| **ROI** | **FA** | | | **MD** | | |
| --- | --- | --- | --- | --- | --- | --- |
|  | **Beta coef** | **p-value** | **Cohens’d** | **Beta coef** | **p-value** | **Cohen’s d** |
| Atr | -0.000205 | 0.663 | -0.009 | 0.000937 | 0.090 | 0.035 |
| Cgc | 0.000643 | 0.357 | 0.019 | 0.000707 | 0.279 | 0.023 |
| Cgh | 0.000070 | 0.922 | 0.002 | 0.001369 | 0.085 | 0.036 |
| Cst | -0.000200 | 0.653 | -0.009 | 0.000977 | 0.036 | 0.044 |
| Fscs | -0.000092 | 0.825 | -0.005 | 0.000570 | 0.131 | 0.031 |
| Fxcut | 0.000796 | 0.191 | 0.027 | 0.002216 | 0.157 | 0.029 |
| Fx | 0.000503 | 0.327 | 0.020 | 0.002299 | 0.046 | 0.042 |
| Ifo | 0.000561 | 0.209 | 0.026 | 0.000988 | 0.030 | 0.045 |
| Ifsfc | 0.000190 | 0.633 | 0.010 | 0.001105 | 0.010 | 0.054 |
| Ilf | 0.000441 | 0.339 | 0.020 | 0.001145 | 0.022 | 0.048 |
| Pscs | -0.000087 | 0.835 | -0.004 | 0.000469 | 0.238 | 0.025 |
| Pslf | 0.000035 | 0.928 | 0.002 | 0.000233 | 0.582 | 0.011 |
| Scs | -0.000053 | 0.892 | -0.003 | 0.000546 | 0.150 | 0.030 |
| Sifc | 0.000176 | 0.732 | 0.007 | 0.001305 | 0.008 | 0.055 |
| Slf | 0.000140 | 0.708 | 0.008 | 0.000207 | 0.609 | 0.011 |
| Tslf | 0.000242 | 0.516 | 0.014 | 0.000269 | 0.498 | 0.014 |
| Unc | 0.000790 | 0.115 | 0.033 | -0.000001 | 0.999 | 0.000 |
| allfibers | 0.000190 | 0.548 | 0.012 | 0.000817 | 0.038 | 0.043 |

##### *Interaction Model with winsorised data and covariates*

Table 51: Beta coefficient, uncorrected p-value, and effect-sizes from the models investigating associations between cortical GM features and E2 timing*tempo

| **ROI** | **Thickness** | | | **Sulcal Depth** | | | **Surface area** | | | **Cortical volume** | | |
| --- | --- | --- | --- | --- | --- | --- | --- | --- | --- | --- | --- | --- |
|  | Beta coef | p-value | Cohen’s d | Beta coef | p-value | Cohen’s d | Beta coef | p-value | Cohen’s d | Beta coef | p-value | Cohen’s d |
| bankssts | 0.001196 | 0.226 | 0.023 | -0.001102 | 0.692 | -0.008 | -0.236930 | 0.731 | -0.007 | -1.977849 | 0.365 | -0.018 |
| caudal anterior cingulate | -0.000724 | 0.574 | -0.011 | 0.001286 | 0.522 | 0.012 | -0.049350 | 0.914 | -0.002 | -1.686890 | 0.349 | -0.018 |
| caudal middle frontal | 0.000020 | 0.987 | 0.000 | -0.001782 | 0.456 | -0.014 | 0.432033 | 0.831 | 0.004 | -0.378901 | 0.954 | -0.001 |
| cuneus | 0.000929 | 0.379 | 0.017 | 0.001361 | 0.539 | 0.012 | -1.075844 | 0.197 | -0.025 | 0.130073 | 0.961 | 0.001 |
| entorhinal | 0.001966 | 0.503 | 0.013 | -0.004121 | 0.475 | -0.014 | -0.836089 | 0.129 | -0.029 | -2.800138 | 0.298 | -0.020 |
| fusiform | 0.001412 | 0.135 | 0.029 | -0.001081 | 0.505 | -0.013 | -0.687728 | 0.566 | -0.011 | 3.382645 | 0.522 | 0.012 |
| inferior parietal | 0.001132 | 0.264 | 0.022 | 0.000866 | 0.498 | 0.013 | -3.398998 | 0.233 | -0.023 | -6.956277 | 0.448 | -0.015 |
| inferior temporal | 0.000825 | 0.454 | 0.015 | 0.001362 | 0.377 | 0.017 | 0.229290 | 0.889 | 0.003 | 0.979438 | 0.883 | 0.003 |
| isthmus cingulate | 0.000200 | 0.824 | 0.004 | 0.000043 | 0.984 | 0.000 | 0.254803 | 0.637 | 0.009 | 1.430724 | 0.430 | 0.015 |
| lateral occipital | 0.000690 | 0.488 | 0.013 | -0.002165 | 0.100 | -0.032 | -0.175461 | 0.933 | -0.002 | 3.292078 | 0.642 | 0.009 |
| lateral orbitofrontal | 0.000934 | 0.450 | 0.015 | 0.000459 | 0.807 | 0.005 | -0.154296 | 0.944 | -0.001 | -0.065506 | 0.992 | 0.000 |
| lingual | 0.002040 | 0.037 | 0.040 | 0.004327 | 0.005 | 0.055 | -1.194723 | 0.355 | -0.018 | 5.126733 | 0.195 | 0.025 |
| Medial orbitofrontal | 0.002048 | 0.125 | 0.030 | 0.000984 | 0.657 | 0.009 | -1.946958 | 0.238 | -0.023 | -3.353005 | 0.514 | -0.013 |
| middle temporal | 0.001760 | 0.121 | 0.030 | 0.001448 | 0.341 | 0.018 | -2.156174 | 0.225 | -0.024 | 0.561266 | 0.930 | 0.002 |
| parahippocampal | 0.001818 | 0.201 | 0.025 | 0.001435 | 0.602 | 0.010 | 0.031580 | 0.946 | 0.001 | 1.172170 | 0.507 | 0.013 |
| paracentral | 0.000509 | 0.673 | 0.008 | 0.000043 | 0.983 | 0.000 | -0.160303 | 0.839 | -0.004 | -0.612007 | 0.850 | -0.004 |
| pars opercularis | 0.000366 | 0.703 | 0.007 | -0.005837 | 0.068 | -0.035 | 0.161149 | 0.883 | 0.003 | -1.231026 | 0.755 | -0.006 |
| pars orbitalis | -0.000297 | 0.827 | -0.004 | 0.003695 | 0.138 | 0.029 | -0.157841 | 0.735 | -0.007 | 0.331106 | 0.871 | 0.003 |
| pars triangularis | 0.000142 | 0.899 | 0.002 | -0.003694 | 0.078 | -0.034 | 0.611460 | 0.565 | 0.011 | 2.164022 | 0.556 | 0.011 |
| pericalcarine | 0.000074 | 0.953 | 0.001 | 0.000793 | 0.746 | 0.006 | -1.900111 | 0.038 | -0.040 | -2.568719 | 0.206 | -0.025 |
| postcentral | 0.000652 | 0.563 | 0.011 | -0.001306 | 0.438 | -0.015 | -1.999007 | 0.408 | -0.016 | -5.687178 | 0.421 | -0.016 |
| posterior cingulate | -0.000364 | 0.652 | -0.009 | 0.000638 | 0.743 | 0.006 | -0.400895 | 0.473 | -0.014 | -3.010042 | 0.113 | -0.031 |
| precentral | -0.000655 | 0.605 | -0.010 | 0.001606 | 0.208 | 0.024 | -6.540739 | 0.030 | -0.042 | -17.856870 | 0.046 | -0.039 |
| precuneus | 0.001331 | 0.132 | 0.029 | 0.000125 | 0.925 | 0.002 | -2.219903 | 0.190 | -0.025 | -4.780378 | 0.463 | -0.014 |
| Rostral anterior cingulate | -0.001041 | 0.565 | -0.011 | -0.003361 | 0.207 | -0.024 | 1.059075 | 0.124 | 0.030 | 2.772927 | 0.302 | 0.020 |
| rostral middle frontal | 0.000278 | 0.803 | 0.005 | 0.002143 | 0.113 | 0.031 | 4.580390 | 0.285 | 0.021 | 8.944920 | 0.496 | 0.013 |
| superior frontal | -0.000384 | 0.731 | -0.007 | -0.000535 | 0.596 | -0.010 | -0.004182 | 0.999 | 0.000 | -6.704029 | 0.664 | -0.008 |
| superior parietal | 0.001735 | 0.138 | 0.029 | 0.000306 | 0.838 | 0.004 | -5.751608 | 0.132 | -0.029 | -3.604954 | 0.773 | -0.006 |
| Superior temporal | 0.000873 | 0.419 | 0.016 | -0.002000 | 0.094 | -0.033 | -0.811163 | 0.638 | -0.009 | -5.459701 | 0.400 | -0.016 |
| supramarginal | 0.001331 | 0.233 | 0.023 | 0.001343 | 0.426 | 0.015 | -4.143434 | 0.163 | -0.027 | 0.463305 | 0.960 | 0.001 |
| frontal pole | 0.001083 | 0.622 | 0.010 | 0.009836 | 0.048 | 0.038 | 0.185993 | 0.561 | 0.011 | 0.459067 | 0.796 | 0.005 |
| temporal pole | 0.001418 | 0.657 | 0.009 | -0.000293 | 0.961 | -0.001 | 0.355780 | 0.449 | 0.015 | -0.090402 | 0.979 | -0.001 |
| transverse temporal | 0.000352 | 0.806 | 0.005 | 0.002993 | 0.342 | 0.018 | -0.271750 | 0.293 | -0.020 | -1.641905 | 0.109 | -0.031 |
| insula | 0.003338 | 0.043 | 0.039 | 0.003867 | 0.380 | 0.017 | -1.795097 | 0.337 | -0.019 | 0.125813 | 0.983 | 0.000 |
| Global | 0.000851 | 0.244 | 0.023 | 0.000139 | 0.263 | 0.022 | -59.040372 | 0.221 | -0.024 | -69.449578 | 0.720 | -0.007 |

Table 52: Beta coefficient, uncorrected p-value, and effect-sizes from the models investigating associations between sub-cortical GM features and E2 timing*tempo

| **Regions** | **Beta coefficient** | **p-value** | **Cohen’s d** |
| --- | --- | --- | --- |
| Thalamus proper | -1.818806 | 0.694 | -0.008 |
| Caudate | 1.248125 | 0.455 | 0.014 |
| Putamen | -1.651194 | 0.542 | -0.012 |
| Pallidum | 1.521301 | 0.449 | 0.015 |
| Hippocampus | 1.504129 | 0.403 | 0.016 |
| Amygdala | -2.571902 | 0.082 | -0.034 |
| Accumbens area | -0.552743 | 0.539 | -0.012 |
| Total subcortical volume | -5.330713 | 0.717 | -0.007 |

Table 53: Beta coefficient, uncorrected p-value, and effect-sizes from the models investigating associations between WM features and E2 timing*tempo

| **ROI** | **FA** | | | **MD** | | |
| --- | --- | --- | --- | --- | --- | --- |
|  | **Beta coef** | **p-value** | **Cohens’d** | **Beta coef** | **p-value** | **Cohen’s d** |
| Atr | -0.000077 | 0.762 | -0.006 | 0.000448 | 0.132 | 0.031 |
| Cgc | 0.000558 | 0.139 | 0.031 | 0.000356 | 0.311 | 0.021 |
| Cgh | 0.000196 | 0.609 | 0.011 | 0.000731 | 0.088 | 0.036 |
| Cst | 0.000005 | 0.984 | 0.000 | 0.000462 | 0.065 | 0.039 |
| Fscs | 0.000006 | 0.977 | 0.001 | 0.000222 | 0.275 | 0.023 |
| Fxcut | 0.000528 | 0.108 | 0.034 | 0.001055 | 0.210 | 0.026 |
| Fx | 0.000406 | 0.142 | 0.031 | 0.001152 | 0.062 | 0.039 |
| Ifo | 0.000403 | 0.093 | 0.035 | 0.000431 | 0.078 | 0.037 |
| Ifsfc | 0.000222 | 0.301 | 0.022 | 0.000474 | 0.039 | 0.043 |
| Ilf | 0.000318 | 0.202 | 0.027 | 0.000609 | 0.024 | 0.047 |
| Pscs | -0.000035 | 0.878 | -0.003 | 0.000193 | 0.368 | 0.019 |
| Pslf | 0.000094 | 0.653 | 0.009 | 0.000067 | 0.767 | 0.006 |
| Scs | -0.000015 | 0.942 | -0.002 | 0.000227 | 0.267 | 0.023 |
| Sifc | 0.000186 | 0.503 | 0.014 | 0.000681 | 0.010 | 0.054 |
| Slf | 0.000134 | 0.505 | 0.014 | 0.000058 | 0.789 | 0.006 |
| Tslf | 0.000171 | 0.393 | 0.018 | 0.000097 | 0.650 | 0.009 |
| Unc | 0.000508 | 0.060 | 0.039 | -0.000014 | 0.958 | -0.001 |
| allfibers | 0.000211 | 0.217 | 0.026 | 0.000338 | 0.111 | 0.033 |
